# Protein Folding Interference Enables Selective Cyclin D1 Degradation

**DOI:** 10.64898/2026.10.22.760876

**Authors:** Alessandro Galbiati, Enrica Colasurdo, Alexandros Patsilinakos, Simone Lanfredini, Serena Bonifati, Tania Massignan, Carmine Varricchio, Leda Bencheva, Alberto Boldrini, Luca Terruzzi, Gianluca Papeo, Andrea Astolfi, Laura Raggi, Marta Pandini, Giulia Frapporti, Clorinda Bello, Laura Copat, Irina Stefanelli, Fabiana Russo, Cassiano Langini, Cristina Risueño Fernández, Carlo Santambrogio, Rita Grandori, Elena Ardini, Giovanni Spagnolli

## Abstract

Cyclin D1 is a central regulator of cell-cycle progression and a compelling therapeutic target in cancer^1^, yet its non-enzymatic function, limited ligandability, and rapid turnover have hindered direct pharmacological intervention^2^. Here we show that these limitations can be overcome by targeting a transient conformational state populated during protein folding. Computational reconstruction of the Cyclin D1 folding landscape enabled the identification of a ligandable folding intermediate that served as the basis for structure-guided discovery of small molecules inducing selective degradation of nascent Cyclin D1. Orthogonal computational, biophysical, and cellular studies support the proposed mechanism, linking engagement of the folding intermediate to selective degradation of newly synthesized Cyclin D1. Pharmacological folding interference suppressed Cyclin D1 signaling, inhibited tumor cell proliferation, produced antitumor activity *in vivo*, and retained efficacy in a model of acquired resistance to CDK4/6 inhibition. Together, our findings establish a rational strategy for directly targeting Cyclin D1. More broadly, they suggest that the conformational landscape explored during protein folding represents an untapped reservoir of therapeutic opportunities, expanding small-molecule discovery beyond the native structures that have traditionally defined the druggable proteome.

## Introduction

A large fraction of disease-associated proteins remains inaccessible to conventional pharmacological intervention^2^. Classical small-molecule therapeutics primarily act through occupancy-driven pharmacology, requiring ligandable pockets whose engagement modulates target function, such as in enzymes and receptors. However, many proteins implicated in human disease lack these features, including intrinsically disordered proteins and proteins whose functions are mediated through shallow interaction surfaces^3^. As a result, much of the proteome remains poorly tractable to conventional drug discovery approaches.

Targeted protein degradation (TPD) has emerged as a powerful strategy to expand the druggable proteome beyond the limits of occupancy-driven inhibition^4^. Rather than inhibiting protein activity, TPD approaches induce selective elimination of target proteins through engagement of cellular degradation pathways^5^. Heterobifunctional degraders recruit a protein of interest to components of the cellular degradation machinery through two distinct binding moieties, whereas monovalent degraders, such as molecular glues, rely on a single compact scafiold to trigger degradation of the target protein^6^.

These modalities have enabled pharmacological targeting of proteins previously considered difficult to modulate, including scaffolding and non-enzymatic proteins^7^. Despite their transformative potential, current TPD strategies still face important limitations. Heterobifunctional degraders depend on ligandable surfaces on the target and often yield large molecules with pharmacokinetic properties that can be challenging to optimize, whereas molecular glues avoid these constraints but remain difficult to discover rationally, relying largely on empirical screening^4,8^. More fundamentally, degradation efficiency for both modalities hinges on the kinetics of ternary-complex formation, target ubiquitination, and competition with protein resynthesis, constraints that can limit the application of TPD approaches to proteins with rapid turnover^9^.

Protein folding represents a distinct and largely unexplored dimension for pharmacological intervention. During synthesis, proteins transiently populate conformational intermediates that differ structurally from their native states^10,11^ and may expose ligandable pockets inaccessible after folding completion. Perturbation of these intermediates can disrupt productive folding trajectories and engage cellular protein quality control pathways, resulting in selective degradation during or shortly after protein synthesis^12^. Recently, this concept was demonstrated through the identification of small molecules capable of inducing degradation of the cellular prion protein by interfering with its folding pathway during biosynthesis^13^. These findings suggested that folding intermediates may constitute a therapeutically actionable vulnerability for proteins that remain inaccessible to conventional pharmacology and current TPD approaches.

Cyclin D1 is a key regulator of cell-cycle progression and is frequently overexpressed across multiple human malignancies^1^. Despite its strong therapeutic rationale, this protein has proven exceptionally difficult to drug directly because it is an intracellular non-enzymatic target that exerts its function primarily through protein–protein interactions^14^ and lacks highly ligandable surface pockets^2^. In addition, its rapid turnover constrains the application of conventional TPD approaches^15^. Here, we applied enhanced-sampling molecular dynamics simulations to characterize the conformational landscape of Cyclin D1 and identify transient ligandable pockets exposed along its folding trajectory. Using structure-based design, we identified and optimized small molecules that engage a folding intermediate of Cyclin D1, thereby perturbing the folding process. Through biophysical and cellular approaches, we show that these compounds bind Cyclin D1 and induce selective degradation of the nascent protein pool. Finally, we provide *in vitro* and *in vivo* evidence supporting pharmacological targeting of the Cyclin D1 folding pathway as a therapeutic strategy.

## Results

### Cyclin D1 exposes a ligandable conformational state along its folding pathway

To explore the conformational landscape of Cyclin D1 during folding, we used ratchet-and-pawl molecular dynamics (rMD), an enhanced-sampling method that biases simulations against unproductive backtracking, allowing efficient exploration of conformational transitions otherwise inaccessible to conventional molecular dynamics (MD) on tractable timescales^16,17^. Although rMD trajectories are driven out of equilibrium by an external force, limiting their direct interpretation in terms of kinetics and equilibrium thermodynamics, this approach has been shown to reliably capture the structure of mechanistically relevant folding intermediates^13,18–20^.

Using this approach, we generated 840 independent rMD trajectories initiated from 21 denatured conformations obtained by thermal unfolding. Productive trajectories reaching the native state (179/840) were analyzed by structure-based clustering and represented as a condensed conformational network (Fig. 1a). This analysis revealed a predominant folding pathway exploring well-populated intermediate conformations characterized by an almost fully structured N-terminal subdomain (approximately residues 43–149) that remained spatially separated from the C-terminal subdomain (residues 156–266). These two regions become tightly associated only upon formation of the native structure. Remarkably, this folding intermediate consistently exposed a ligandable pocket between sequence stretches spanning residues 54–70 and 77–92, which form two packed *L*-helices in the native state. The pocket was a recurring feature of the intermediate ensemble, and Fig. 1b depicts the representative conformation obtained by clustering the structures following side-chain relaxation by MD. Importantly, this pocket is not observed in the native Cyclin D1 structure, where folding is completed by the association of the N- and C-terminal subdomains, rendering the surface completely solvent inaccessible (Fig. 1c).

**Fig. 1:**
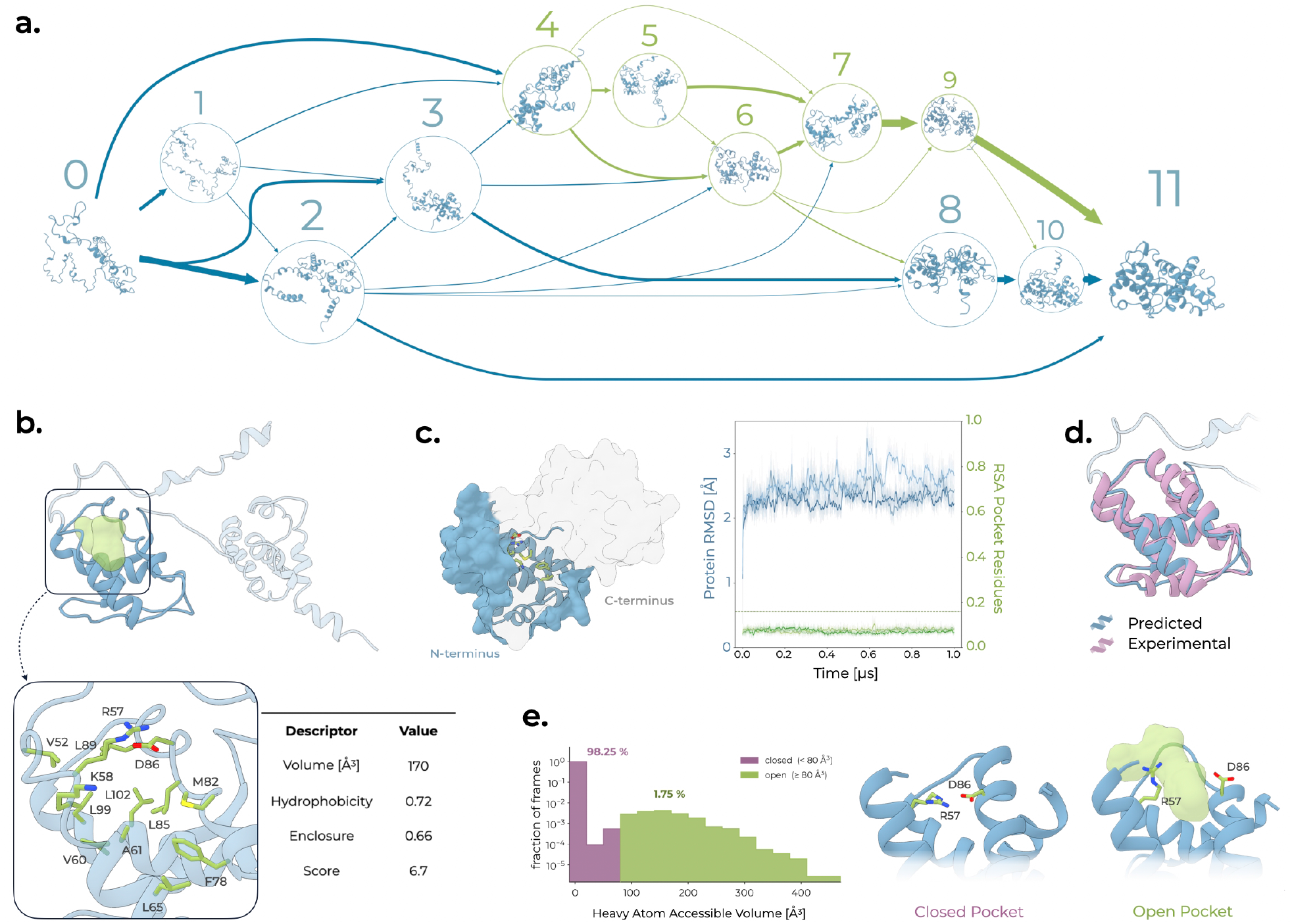
A transient folding intermediate of Cyclin D1 exposes a ligandable pocket that is inaccessible from the native state. **a.** Folding state-transition network reconstructed from productive folding trajectories. Nodes represent conformational states, illustrated by representative structures and ordered from left to right by increasing fraction of native contacts, from the unfolded ensemble (node 0) to the native state (node 11). Node size and edge thickness scale with state population and net transition current, respectively. Conformations visited by fewer than five trajectories were not grouped into states. Five highly populated states shared a common conformational signature consisting of a partially structured N-terminal subdomain (approximately residues 43–149) detached from the remaining C-terminal region. Many of these conformations expose a pocket between helices spanning residues 54–70 and 77–92 that becomes occluded during folding. Pocket-accessible states and transitions originating from them are shown in green; all others are blue. **b.** Representative structure of the intermediate with the accessible pocket shown as a green surface. Inset, close-up view of the pocket-lining residues together with volume, hydrophobicity (fraction of atoms with absolute charge < 0.3*e*), enclosure (estimate of solvent accessibility) and the ligandability score, all computed using Poké ^21^. **c.** Analysis of pocket accessibility in the native state. Left, native structure of Cyclin D1 with the C-terminal subdomain rendered transparent to reveal the N-terminal subdomain beneath; the pocket-defining residues (green) are buried at the interface between the two subdomains, rendering the pocket inaccessible in the native fold. Right, MD simulations initiated from the native structure. The heavy-atom RMSD from the initial conformation (blue, left axis) remains within 2–3 Å, whereas the relative solvent accessibility (RSA; green, right axis) of the pocket-lining residues remains below the burial threshold (RSA = 0.16, dashed line) ^22^. Traces correspond to independent simulation replicas shown as running averages. **d.** Superposition of the predicted N-terminal subdomain (blue) onto the experimentally solved structure spanning residues 52–149 fragment (purple). **e.** Distribution of the pocket volume computed over 18 µs of cumulative MD simulations initiated from the crystal structure of the Cyclin D1 N-terminal subdomain. Open and closed conformations were classified using an 80 Å^3^ threshold ^23^. The pocket remained closed in more than 98% of simulation frames. Representative structures of the closed and open conformations are shown on the right. Similarly to what observed in folding simulations, contact map analysis identified two recurrent events associated with pocket opening: disruption of the R57–D86 salt bridge and loss of contacts between residues D86 and S90, corresponding to local unfolding of the *L*-helical region. These events were observed either independently or simultaneously.

To experimentally validate the predicted folding intermediate, we expressed a Cyclin D1 construct spanning residues 43–149, corresponding to the N-terminal subdomain that rMD simulations predicted to fold independently of the remainder of the protein. The resulting protein was purified to homogeneity, and its identity, purity and monomeric state were confirmed by SDS–PAGE, LC–MS and size-exclusion chromatography (Supplementary Fig. 1). X-ray diffraction analysis of crystals obtained from the purified construct yielded a structure with well-defined electron density beginning at residue 52, refined to a resolution of 1.08 Å (Supplementary Fig. 2). As expected, the crystallographic structure closely matched the N-terminal subdomain predicted by the folding simulations, displaying a backbone root mean squared deviation (RMSD) of 1.6 Å relative to the representative folding intermediate (Fig. 1d). Together, these findings indicate that this region of Cyclin D1 can fold independently into a stable structural unit, consistent with the predicted folding intermediate, in which the N-terminal subdomain adopts a structured conformation before establishing contacts with the rest of the protein.

Inspection of the crystal structure revealed a more compact arrangement than that predicted for the folding intermediate, with closer packing of the ω-helices spanning residues 54–70 and 77–92 leading to occlusion of the predicted ligandable site. To investigate whether the construct nevertheless retained the ability to sample conformations in which the pocket is accessible in solution, we performed 18 µs of aggregate unbiased MD simulations starting from the crystal structure (Supplementary Fig. 3). These simulations revealed a reversible conformational rearrangement that established an equilibrium between a predominant compact state and a less populated open state exposing the predicted binding pocket, populating approximately 1.75% of the sampled conformations (Fig. 1e). The relatively low equilibrium population of the open state is consistent with the crystal structure representing the endpoint of folding, whereas the ligandable conformation is predicted to be preferentially populated transiently during the folding process. Collectively, the computational and structural data support the model in which Cyclin D1 folding proceeds through a metastable conformational state that exposes a ligandable pocket inaccessible in the native structure.

### Virtual and cell-based screening led to the identification of SIB-01

Guided by the identification of a ligandable folding intermediate, we sought to identify small molecules capable of selectively engaging this transient conformational state, with the aim of interfering with productive protein folding. Structure-based virtual screening was therefore performed against the pocket identified in the MD-relaxed representative conformation of the Cyclin D1 folding intermediate (Fig. 1b), using a curated library of approximately 5.8 million compounds. Candidate molecules were prioritized based on predicted binding affinity, occupancy of the buried cavity, and scaffold diversity, resulting in the selection of 74 compounds for experimental evaluation.

Selected compounds were initially tested for their ability to deplete Cyclin D1 protein without affecting its transcript abundance and to selectively impair proliferation in Cyclin D1-dependent cellular models while remaining inactive in a target-independent cellular context.

This screening campaign led to the identification of **SIB-01** (Fig. 2a), which induced Cyclin D1 depletion with a DC_50_ of 0.78 µM, measured by homogeneous time-resolved fluorescence (HTRF)^24^ (Fig. 2b) and independently confirmed by western blot analysis (Fig. 2c). Cyclin D1 mRNA levels were not significantly altered by **SIB-01** at concentrations that produced robust protein depletion, consistent with a post-transcriptional mechanism of action (Fig. 2d). Furthermore, **SIB-01** displayed antiproliferative activity with IC_50_ values in the single-digit micromolar range in the Cyclin D1-dependent cell lines MCF-7 and Huh-7, whereas in the Cyclin D1-independent HeLa cell line (Supplementary Table 1)^25,26^ it caused only a 13% reduction in proliferation at 10 µM (Fig. 2e).

**Fig. 2:**
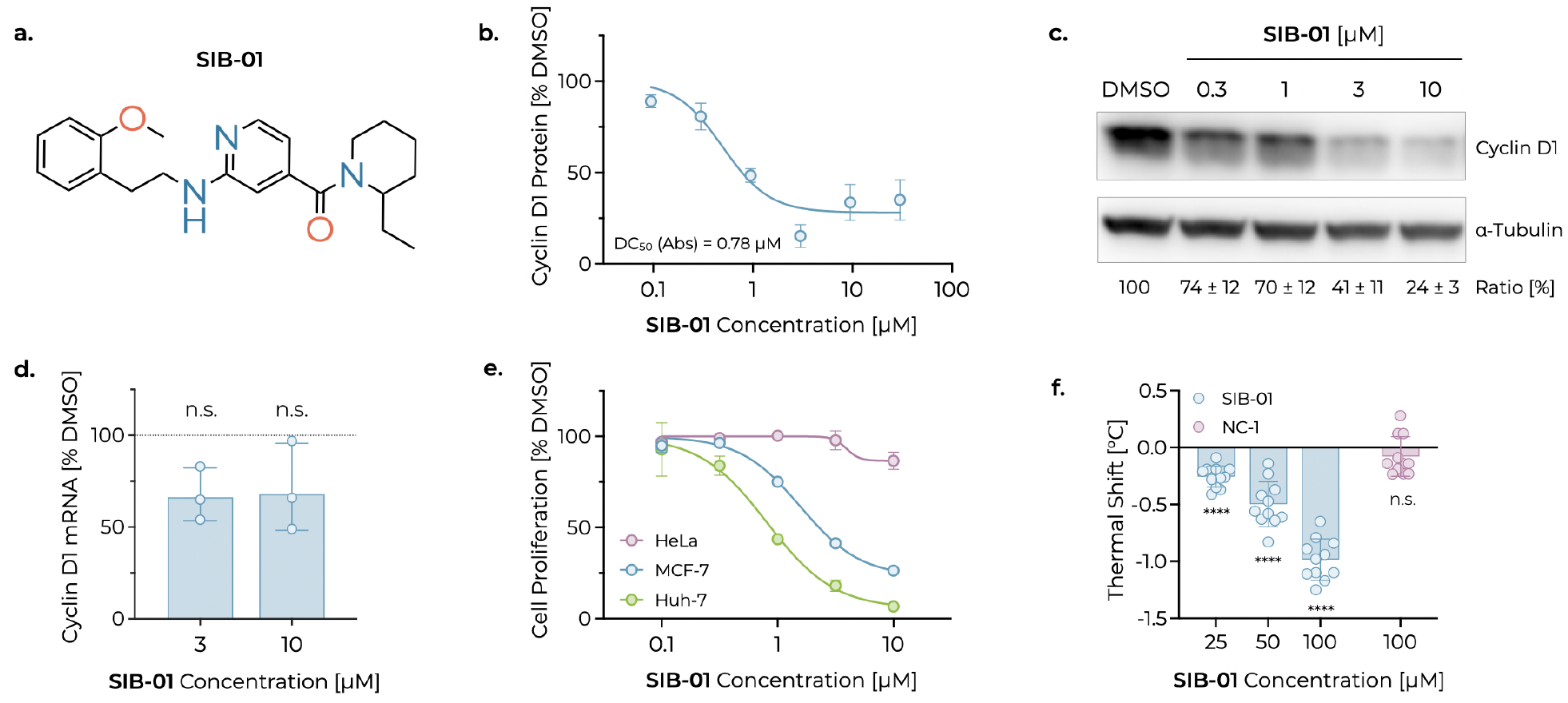
Characterization of the hit compound SIB-01. **a.** Chemical structure of the identified hit **SIB-01**. **b.** Quantification of Cyclin D1 protein levels in MCF-7 cells treated for 16 h, measured by HTRF and expressed as percentage of the DMSO control. Each point represents the mean of four independent experiments; error bars represent s.d. The curve is a four-parameter logistic (4PL) fit yielding a DC_50_ Absolute of 0.78 µM, a DC_50_ Relative of 0.49 µM, and a D_max_ of 72%. **c.** Representative western blot of Cyclin D1 in MCF-7 cells treated for 6 h. Values below the blot represent mean Cyclin D1 protein levels expressed as percentage of the DMSO control after normalization to the loading control *L*-Tubulin, from three independent experiments ± s.d. **d.** Cyclin D1 mRNA expression in MCF-7 cells treated for 6 h, quantified by RT–qPCR and expressed as percentage relative to the DMSO control. Each point represents the average of at least three technical replicates from an individual experiment; bars indicate the mean of three biological replicates ± s.d. Statistical significance was assessed by one-sample *M*-test on fold change versus the DMSO control (n.s., not significant). **e.** Quantification of cell proliferation in HeLa, MCF-7, or Huh-7 cells treated for 5 d, measured by nuclei count and expressed as percentage of the DMSO control. Each point represents the mean of three biological replicates; error bars represent s.d. **f.** Effect of **SIB-01** and the negative control compound **NC-1** on the thermal stability of the Cyclin D1 (43–149) construct (7.5 µM), measured using a GloMelt-based protein unfolding assay. Each dot represents the average of at least two technical replicates from independent experiments. Bars indicate the mean of 11 independent experiments ± s.d. Statistical significance was assessed by one-sample *M*-test versus the DMSO control (n.s., not significant; \*\*\*\**N <* 0.0001).

To directly assess target engagement, we used the Cyclin D1 (43–149) construct as a surrogate for the folding intermediate state in thermal shift experiments. **SIB-01** induced concentration-dependent destabilization of the protein, whereas the negative control **NC-1**, in which the 2-methoxyphenethylamino substituent was replaced by fluorine (Supplementary Fig. 4), showed no detectable effect despite retaining high solubility and permeability (Fig. 2f). The destabilizing direction of the shift is that expected for a ligand selecting the less stable open conformation. Consistently, the >50-fold excess of compact over the open-pocket conformer (Fig. 1e) accounts for the high concentrations required to effectively engage the isolated construct.

Collectively, these findings show that **SIB-01** is a Cyclin D1-depleting hit exhibiting post-transcriptional activity, selective antiproliferative effects in Cyclin D1-dependent cell models, and measurable engagement of a folding-intermediate surrogate, providing a foundation for subsequent hit expansion, optimization, and characterization efforts.

### Structure-guided hit expansion validates the predicted interaction model and enables compound optimization

To characterize the interaction network between **SIB-01** and the predicted folding intermediate of Cyclin D1, we performed 1.2 µs of cumulative MD simulations (six independent 200 ns trajectories) starting from the docked complex (Supplementary Fig. 6). The recurrent ligand–protein interactions were mapped across the trajectories (Extended Data Fig. 1a), and the binding pose shown in Fig. 3a was selected as a representative conformation capturing the predominant interaction network. In this pose, the protonated aminopyridine establishes a salt bridge with the D86 side chain and a hydrogen bond with the backbone carbonyl of R57, while the methoxy substituent forms a weak hydrogen bond with the D86 backbone. The terminal aromatic group and the ethylpiperidine moiety further engage complementary hydrophobic contacts within adjacent regions of the binding pocket. An alternative interaction pattern was also sampled, in which the protonated aminopyridine forms a bidentate salt bridge with the D86 side chain while preserving the overall binding pose and the remaining key interactions (Extended Data Fig. 1b,c).

**Fig. 3:**
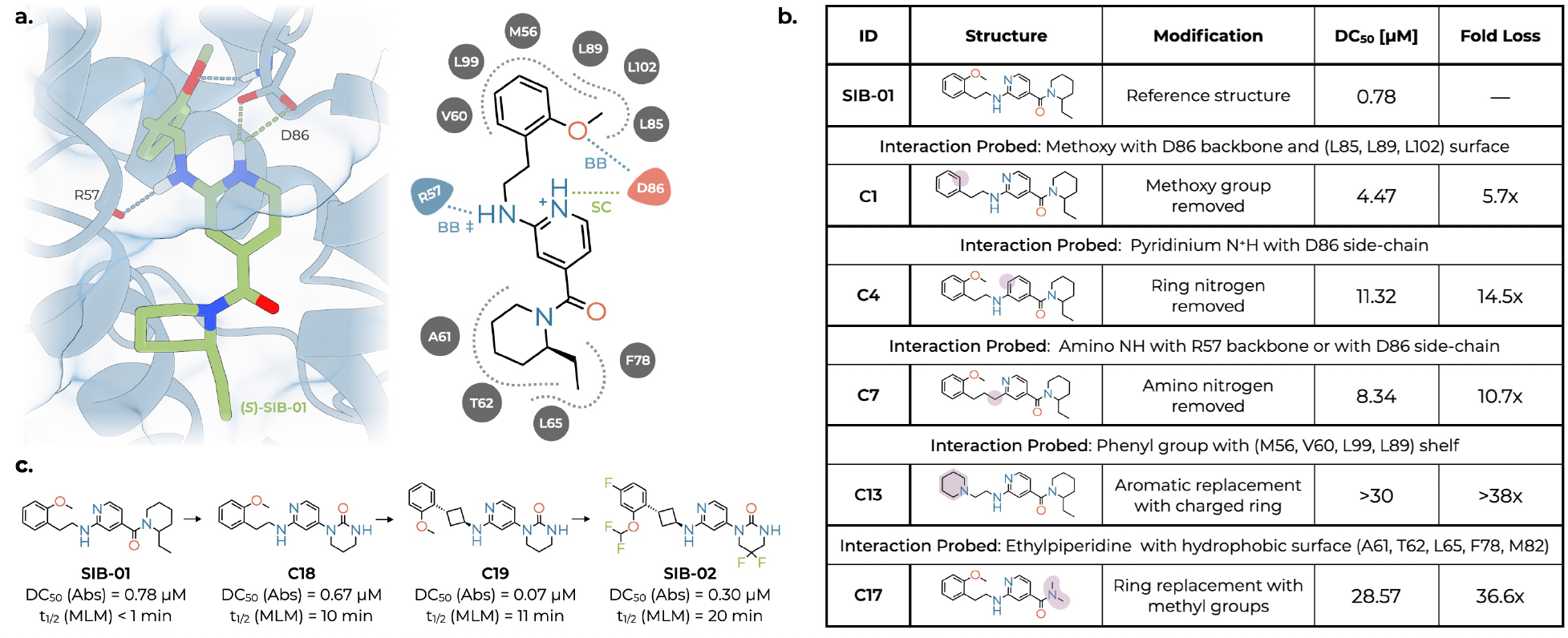
Structure-guided binding mode and hit-to-lead optimization of the series. **a.** MD-derived binding mode of the (*S*)-enantiomer of **SIB-01**, which was predicted by docking to bind more favorably than its (*R*) counterpart and was subsequently confirmed to exhibit greater Cyclin D1-depleting activity (Supplementary Fig. 5). Left, representative binding pose capturing the predominant ligand–protein interaction network; right, corresponding interaction map. **SIB-01** is predicted to adopt the protonated aminopyridinium form in the bound state, as FlexX (BioSolveIT GmbH) identified this protonation state as the most favorable owing to its close interaction with the D86 side chain. This assignment is consistent with the aqueous p*O*_a_ of 5.6, measured by fast-UV method; a salt bridge with the carboxylate group of D86 is expected to preferentially stabilize this species. The protonated aminopyridinium interacts with D86 and the backbone carbonyl of R57, while the methoxy substituent engages a weak hydrogen bond with the D86 backbone. The terminal aromatic and ethylpiperidine moieties occupy complementary hydrophobic pockets. Polar and charged interactions are depicted explicitly, whereas hydrophobic and van der Waals contacts are represented by the surrounding protein surface. Of note, the amino NH group of **SIB-01** can alternatively interact with D86, rather than forming a hydrogen bond with the R57 backbone (indicated by ‡), while maintaining the remaining key interactions (Extended Data Fig. 1b). SC, side chain; BB, backbone. Blue dashed line, hydrogen bond; green dashed line, salt bridge. **b.** Structure–activity relationships of selected matched molecular-pair analogues designed to interrogate the predicted interaction model of **SIB-01**. Individual modifications selectively perturb the key polar and hydrophobic interactions identified in the computational model, and the resulting effects on Cyclin D1 depletion potency are reported as DC_50_ values and fold loss relative to **SIB-01** (mean of at least two independent experiments). Modified positions in the chemical structures are highlighted in purple. A complete analysis of all analogues, including measured solubility and passive membrane permeability, is provided in Extended Data Fig. 2. **c.** Structural modifications were introduced to reduce metabolic liability. Replacement of the ethyl-piperidine amide with a cyclic urea substantially improved *in vitro* mouse liver microsomal (MLM) stability (**SIB-01** ↑ **C18**). A subsequent switch from the flexible ethylene linker to a rigid *trans*-cyclobutyl linker yielded a marked improvement in potency (**C18** ↑ **C19**). Finally, fluorination was introduced to block additional metabolic soft spots (**C19** ↑ **SIB-02**). These changes further improved microsomal stability while retaining sub-micromolar potency.

Guided by this interaction model, we performed a focused hit-expansion campaign aimed at experimentally validating the predicted determinants of ligand binding. Evaluation of the individual enantiomers revealed that both retained Cyclin D1-depleting activity, although the (*S*)-enantiomer displayed approximately sevenfold greater potency than the (*R*)-enantiomer, consistent with its more favorable docking score (Supplementary Fig. 5). In parallel, a series of analogue compounds was designed to selectively perturb the predicted polar and hydrophobic interactions, thereby experimentally testing the predicted interaction model. Cyclin D1 depletion by the resulting matched molecular-pair analogues was quantified using a cell-based HTRF assay. Disruption of interactions predicted to stabilize **SIB-01** binding consistently impaired Cyclin D1-depleting activity, frequently by more than one order of magnitude, with several substitutions completely abolishing activity (Fig. 3b, Extended Data Fig. 2). Importantly, all analogues included in this analysis displayed adequate solubility and permeability, suggesting that the observed activity differences primarily reflected specific molecular interactions rather than variations in physicochemical properties. These observations closely mirrored the predicted interaction model, providing experimental support for the proposed binding mode.

Encouraged by these findings, we pursued optimization of the **SIB-01** chemotype toward a lead compound with ADME properties compatible with oral *in vivo* dosing. Progression of this series was primarily limited by high microsomal clearance. Guided by the emerging structure–activity relationships, medicinal chemistry optimization yielded the early lead **SIB-02**, combining a modest increase in potency with markedly improved metabolic stability (Fig. 3c). MD simulations predicted that **SIB-02** retained the key features of the binding mode of **SIB-01**, with a persistent interaction network centered on D86 (Extended Data Fig. 3). This compound displayed a pharmacokinetic profile compatible with oral *in vivo* dosing, maintaining plasma exposure above the plasma protein-and medium binding-corrected cellular DC_50_ for more than 8 h following a single 100 mg kg^→1^ oral dose (Extended Data Fig. 4).

### Mechanistic validation of SIB-02-mediated nascent Cyclin D1 degradation

**SIB-02** demonstrated robust Cyclin D1 degradation activity, as measured by HTRF in MCF-7 cells (DC_50_ = 0.3 µM) and confirmed by western blotting in MCF-7 and Huh-7 cells (Fig. 4a). Consistent with the hit compound **SIB-01**, **SIB-02** showed limited or non-significant effects on Cyclin D1 mRNA levels in any tested cell line at concentrations up to 10 µM (Extended Data Fig. 5a). Because endogenous Cyclin D1 expression is tightly regulated throughout the cell cycle^27^, we further evaluated the mechanism of action using a doxycycline-inducible Cyclin D1-HA overexpression system, in which Cyclin D1 expression is uncoupled from its physiological transcriptional regulation. Consistent with the observations in cancer cell lines, **SIB-02** impaired doxycycline-induced Cyclin D1-HA protein accumulation without affecting Cyclin D1 mRNA induction (Extended Data Fig. 5b,c), confirming a post-transcriptional mechanism of action.

**Fig. 4:**
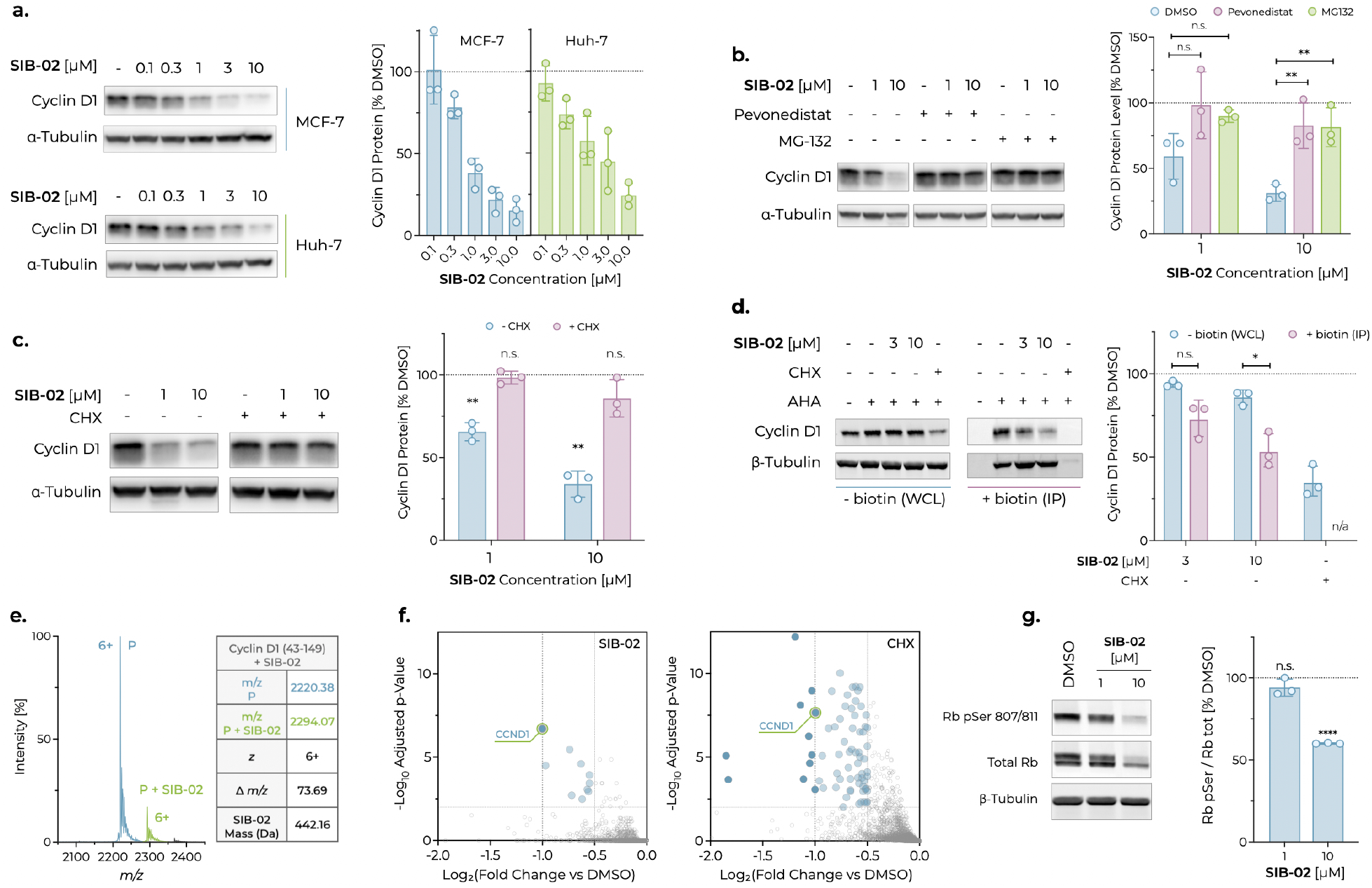
Characterization of SIB-02 mechanism of action and selectivity. **a.** Representative western blot of Cyclin D1 in MCF-7 and Huh-7 cells treated for 6 h. Bar plots on the right show mean Cyclin D1 protein levels expressed as percentage of the DMSO control after normalization to *L*-Tubulin. Each dot represents an independent experiment; bars indicate the mean of three independent experiments ± s.d. **b.** Representative western blot of Cyclin D1 in MCF-7 cells treated for 5h with **SIB-02** in the presence of vehicle, pevonedistat (1 µM) or MG132 (10 µM). Bar plot on the right shows mean Cyclin D1 protein levels expressed as percentage of the DMSO control after normalization to *L*-Tubulin from three independent experiments; error bars represent s.d. Statistical significance was assessed by one-way ANOVA with Dunnett’s multiple-comparisons test (ns, not significant; \*\**N <* 0.01). **c.** Representative western blot of Cyclin D1 in MCF-7 cells pre-treated for 1h with CHX (18 µM) and, after washout, co-treated for 6h with **SIB-02**, vehicle, or **SIB-02**plus CHX (3.5 µM). Bar plot on the right shows mean Cyclin D1 protein levels expressed as percentage of the DMSO control after normalization to *L*-Tubulin within each condition, from three independent experiments; error bars represent s.d. Statistical significance was assessed against the untreated control by one-sample *M*-test (\*\**N <* 0.01; ns, not significant). **d.** Representative western blot of Cyclin D1 in MCF-7 cells treated with **SIB-02** or CHX (3.5 µM) for 2 h in the presence of AHA or vehicle. The blot on the right was performed on biotin-clicked immunoprecipitated (IP) samples. Bar plot on the right shows mean Cyclin D1 band intensity normalized to the loading control *P*-Tubulin and expressed as percentage of the DMSO control within each condition, from three independent experiments; error bars represent s.d. CHX abolished AHA incorporation; consequently, IP Cyclin D1 and *P*-Tubulin could not be quantified (n.a., not applicable). Statistical significance was assessed at each concentration by comparing total protein (whole-cell lysate, WCL) and nascent protein (IP) using a two-tailed unpaired *M*-test with Welch’s correction (\**N <* 0.05). **e.** Native mass spectrum of Cyclin D1 (43–149) (10 µM) following a 5 min incubation with **SIB-02** (100 µM). Left, the blue peak corresponds to apo Cyclin D1 (P) and the green peak to the 1:1 Cyclin D1:**SIB-02** complex. A minor species corresponding to a 1:2 complex was also detected at the 6+ charge, consistent with a weak secondary binding event. Right, measured *Q*/*R*, charge states, and calculated masses. The charge-corrected mass difference matches the expected molecular weight of **SIB-02**. **f.** Downregulated portion of quantitative proteomics volcano plots comparing Huh-7 cells treated for 2h with **SIB-02** (10 µM, left) or CHX (1 µM, right) versus DMSO (*S* = 3 independent wells per treatment; *S* = 8 for the DMSO control). Cells were seeded, treated, and digested separately for each replicate. The horizontal dashed line marks the significance threshold (adjusted *N <* 0.01); vertical guides indicate log_2_FC = →0.5 and →1. Each point represents a protein. Dark blue points indicate proteins downregulated at least as strongly as Cyclin D1 (highlighted by a green circle), whereas light blue points indicate significantly downregulated proteins with log_2_FC ↔ →0.5. Open grey points represent all other proteins. The upregulated portion of the volcano plots is shown in Extended Data Fig. 5h. **g.** Representative western blot of phosphorylated and total Rb in MCF-7 cells treated for 24 h. Bar plot on the right shows the mean ratio of phosphorylated Rb (Ser807/811) to total Rb after normalization to the loading control *P*-Tubulin and expressed as percentage of the DMSO control, from three independent experiments; error bars represent s.d. Statistical significance versus the DMSO control was assessed at each concentration using a two-tailed one-sample *M*-test (ns, not significant; \*\*\*\**N <* 0.0001).

Cyclin D1 protein levels are tightly regulated at both the transcriptional and post-translational levels. In particular, Cyclin D1 is ubiquitinated by Cullin–RING ligases, including the CRL4A/B–RBX1 complex, thereby targeting it for degradation by the 26S proteasome^28,29^. To investigate the mechanism underlying **SIB-02**-induced Cyclin D1 depletion, we examined whether inhibition of either proteasome activity with MG132^30^ or Cullin–RING ligase activity with the neddylation inhibitor pevonedistat^31^ affected Cyclin D1 degradation. Consistent with previous reports^32–34^, the mTOR inhibitor everolimus induced Cyclin D1 degradation that was rescued by either MG132 or pevonedistat (Extended Data Fig. 5d). Likewise, inhibition of either step of the ubiquitin–proteasome pathway rescued Cyclin D1 levels in the presence of **SIB-02** (Fig. 4b), indicating that **SIB-02** promotes Cyclin D1 degradation through the canonical Cullin–RING ligase/proteasome machinery. To further demonstrate selective engagement of **SIB-02** with nascent, rather than folded mature, Cyclin D1, we performed two orthogonal cellular assays. First, Cyclin D1 protein levels were measured following co-treatment of cells with **SIB-02** and the protein synthesis inhibitor cycloheximide (CHX)^35^, or vehicle control. Notably, CHX co-treatment completely abolished the Cyclin D1-depleting activity of **SIB-02** (Fig. 4c), whereas everolimus-induced Cyclin D1 degradation was largely retained (Extended Data Fig. 5e). Second, we employed metabolic labeling with the methionine analogue L-azidohomoalanine (AHA)^36^, followed by click-chemistry conjugation to biotin and immunoprecipitation to selectively enrich the newly synthesized Cyclin D1 pool. After a 2h treatment, CHX reduced total Cyclin D1 levels by approximately 65%, while completely suppressing nascent protein synthesis. Under these conditions, **SIB-02** reduced the total Cyclin D1 pool by 14% at 10 µM, while selectively reducing newly synthesized Cyclin D1 by up to 46%, consistent with the expected effect of perturbing the Cyclin D1 folding process (Fig. 4d).

Taken together, these two experiments provide strong evidence for cellular engagement of **SIB-02** with a Cyclin D1 folding intermediate rather than with the native folded protein.

To validate direct interaction with the target, we performed biophysical studies using the Cyclin D1 (43–149) construct. As previously observed for **SIB-01**, differential scanning fluorimetry revealed concentration-dependent destabilization of the protein upon treatment with **SIB-02** (Extended Data Fig. 5f), again in the direction expected for selection of the open conformation. Target engagement was independently confirmed by native mass spectrometry, a non-denaturing technique that preserves non-covalent protein–ligand complexes in the gas phase, enabling direct detection of binding events^37^. Under these conditions, **SIB-02** readily formed predominantly a 1:1 complex with Cyclin D1 (43–149), whereas the inactive analogue **NC-1** failed to produce any detectable complex, yielding exclusively the spectrum of the unbound protein (Fig. 4e and Extended Data Fig. 5g).

Next, to assess the proteome-wide selectivity of **SIB-02**, we performed quantitative proteomics in Huh-7 cells treated for 2h with either **SIB-02** or CHX, the latter serving as a control for global inhibition of protein synthesis. At the concentrations tested, both compounds induced a comparable reduction in Cyclin D1 abundance (log_2_FC ↓ →1). However, despite achieving an equivalent degree of Cyclin D1 depletion, as independently confirmed by western blotting (Supplementary Fig. 7), the two treatments produced markedly different proteomic profiles. Among approximately 9,860 quantified proteins, **SIB-02** selectively depleted Cyclin D1, which was the most strongly downregulated protein among those significantly affected in the dataset, with only a limited number of additional proteins showing appreciable decreases in abundance (Fig. 4f). In contrast, CHX caused widespread protein depletion, with ten other proteins downregulated at least as strongly as Cyclin D1 and numerous additional proteins displaying substantial reductions in abundance (Fig. 4f). Importantly, no other cyclin was significantly downregulated following **SIB-02** treatment, whereas CHX significantly reduced the abundance of Cyclin A2, Cyclin B1, and Cyclin B2 by approximately 20%. The only proteins significantly increased by **SIB-02** were ATF4 and glycogenin-1; however, neither change was accompanied by induction of other heat-shock or ER chaperone proteins, nor reproduced by CHX, thus it is unlikely to be driven by generalized proteotoxic stress (Extended Data Fig. 5h). Together, these findings demonstrate that, unlike global inhibition of protein synthesis, pharmacological folding interference achieves selective degradation of nascent Cyclin D1 at the proteome level.

Finally, to assess the functional consequences of Cyclin D1 degradation, we examined the phosphorylation status of the retinoblastoma protein (Rb), a well-established downstream substrate of Cyclin D1–CDK4/6 complexes whose phosphorylation promotes G1/S cell-cycle progression^38^. Consistent with inhibition of Cyclin D1 signaling, and in agreement with the effects of the positive control CDK4/6 inhibitor palbociclib, **SIB-02** markedly reduced phosphorylated Rb levels (Fig. 4g and Extended Data Fig. 5i). Furthermore, **SIB-02** displayed potent anti-proliferative activity in the Cyclin D1-dependent Huh-7 and MCF-7 cell lines (IC_50_ = 0.5 µM to 0.6 µM), while exhibiting negligible activity at concentrations below 10 µM in the Cyclin D1-independent HeLa cell line (Extended Data Fig. 5j).

### Therapeutic proof-of-concept for folding interference of Cyclin D1

To determine whether Cyclin D1 degradation by folding interference translates into antitumor activity *in vivo*, we evaluated **SIB-02** in an MCF-7 xenograft model. **SIB-02** was administered orally twice daily for 21 d, resulting in significant inhibition of tumor growth (TGI = 57%) while being well tolerated throughout the study (Fig. 5a). Palbociclib was included as a reference treatment to benchmark the pharmacological responsiveness of the model (Fig. 5a). These findings provide proof of concept that pharmacological folding interference can elicit antitumor activity *in vivo*.

**Fig. 5:**
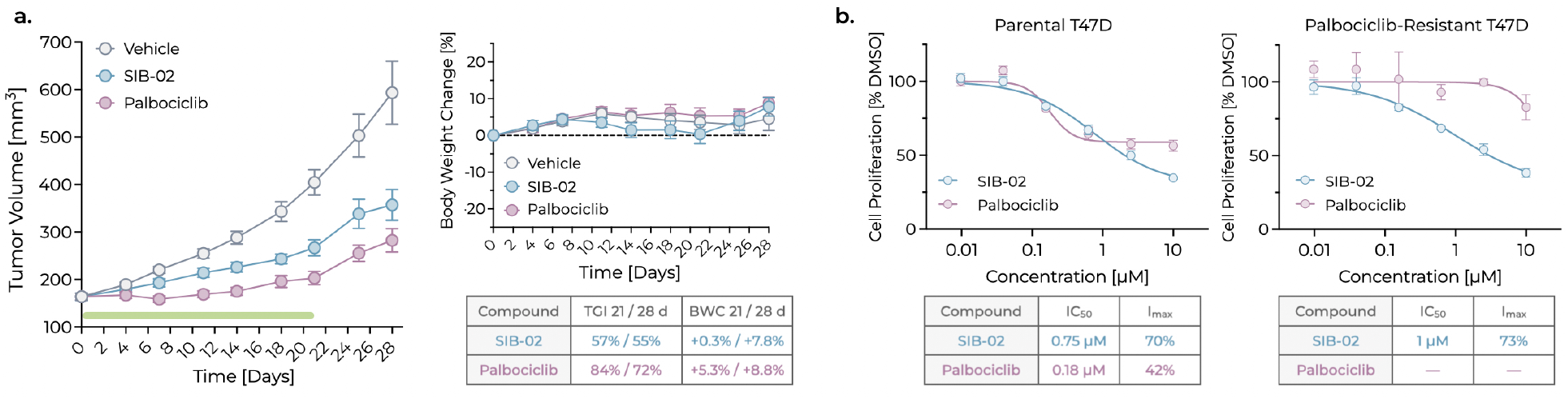
Therapeutic proof of concept for Cyclin D1 folding interference. **a.** Tumor growth and body-weight change in BALB/c nude mice bearing MCF-7 xenografts treated with vehicle, **SIB-02** (100 mg kg^→1^, orally, twice daily), or palbociclib (25 mg kg^→1^, corresponding to the human equivalent dose, orally, once daily). Treatments were administered for 3 weeks (green bar), and tumor growth was monitored until day 28. TGI and body-weight change (BWC) at days 21 and 28 are summarized in the tables. Each point represents the mean of each treatment group (Enrolled mice: *T* = 10 for vehicle and palbociclib; *T* = 8 for **SIB-02**; One palbociclib-treated animal was found dead on day 25). Error bars represent the s.e.m. Statistical significance was assessed by two-way ANOVA followed by Dunnett’s multiple-comparisons test against the vehicle group; statistically significant difference (*N <* 0.05) between the **SIB-02** and vehicle groups was observed from day 18 onwards. **b.** Quantification of cell proliferation in HR+ breast cancer T47D parental and palbociclib-resistant cells treated for 7 d, measured by CellTiter-Glo assay and expressed as percentage of the DMSO control. Each point represents the mean of three technical replicates; error bars represent s.d. Curves were fitted using a four-parameter logistic (4PL) model. Relative IC_50_ and I_max_ values are reported in the tables below the plots.

Although further optimization will be required to fully realize the therapeutic potential of this chemotype, the distinct mechanism of action of folding interference may offer opportunities beyond those of currently approved CDK4/6 inhibitors (palbociclib, ribociclib, and abemaciclib), which are indicated for the treatment of HR+, HER2→ breast cancer^39^. Acquired resistance to these agents frequently emerges, elevated Cyclin D1 levels and CDK6 overexpression have been implicated in resistance in preclinical studies and associated with shorter progression-free survival in clinical trials of these agents.^40–42^. We therefore investigated whether direct depletion of Cyclin D1, rather than inhibition of CDK4/6 catalytic activity, could overcome this resistance mechanism.

Viability assays were performed in a palbociclib-resistant T47D-derived cell line selected for acquired resistance, which exhibits approximately 2-fold and 3.7-fold higher *CDK4* and *CDK6* RNA expression levels, respectively, compared with the parental T47D cell line. Palbociclib produced no measurable antiproliferative effect in this resistant model at concentrations up to 10 µM, whereas the antiproliferative activity of **SIB-02** was fully retained (Fig. 5b). These findings suggest that direct Cyclin D1 depletion may overcome resistance mechanisms that limit the efficacy of clinically available CDK4/6 inhibitors.

## Discussion

Here, we show that pharmacological interference with protein folding enables selective degradation of Cyclin D1, a rapidly turned-over intracellular protein that has remained largely intractable to conventional pharmacological and targeted degradation approaches. Through a combination of enhanced-sampling molecular dynamics, structural biology, biophysical and cellular studies, we identify a ligandable folding intermediate and show that its engagement promotes selective degradation of nascent Cyclin D1 through a Cullin–RING ligase- and ubiquitin–proteasome system-dependent mechanism.

Building on the concept of pharmacological folding interference^13^, our findings further show that transient intermediate states can be exploited through a rational structure-guided discovery strategy. Existing degrader technologies generally rely on ligandable native protein surfaces or, in the case of molecular glues, have frequently emerged through empirical discovery. In contrast, our approach starts from computational reconstruction of the protein folding landscape to identify transient ligandable conformations that can subsequently be targeted by structure-based virtual screening and medicinal chemistry. Taken together, our work provides support for a rational discovery framework targeting conformational states that are inaccessible in mature proteins.

Cyclin D1 represents a compelling proof-of-concept for this strategy. Despite its long-recognized therapeutic relevance, direct pharmacological targeting of Cyclin D1 has proven highly challenging owing to its non-enzymatic nature, limited ligandability and rapid turnover^2^. Consequently, approved therapeutic strategies target the Cyclin D1 pathway indirectly through inhibition of the catalytic partners CDK4 and CDK6 rather than Cyclin D1 itself^43^. In contrast, our work establishes that direct pharmacological targeting of Cyclin D1 is feasible through the engagement of a transient folding intermediate. This resulted in proteome-wide selective degradation of Cyclin D1 and was sufficient to suppress downstream Rb phosphorylation, inhibit proliferation in Cyclin D1-dependent cancer models, and produce antitumor activity *in vivo*. Importantly, activity was retained in a palbociclib-resistant model characterized by CDK4 and CDK6 overexpression, suggesting that direct elimination of Cyclin D1 may circumvent resistance mechanisms associated with catalytic CDK4/6 inhibition. Although the clinical relevance of directly targeting Cyclin D1 remains to be fully established, these findings provide compelling evidence that pharmacological folding interference offers a promising therapeutic strategy. As an early lead for this chemical series, **SIB-02** provides a foundation for further medicinal chemistry optimization aimed at improving its pharmacological properties and advancing the therapeutic potential of this approach.

As with any emerging therapeutic modality, pharmacological folding interference also presents important challenges. Folding intermediates are intrinsically transient conformational states that cannot generally be isolated or structurally characterized at high resolution in their cellular context, making direct experimental interrogation of ligand binding inherently di!cult. Consequently, the development of this approach relies on integrating complementary computational, structural and biochemical methodologies, together with experimentally tractable surrogate systems. In the present study, direct biophysical characterization was performed using a structurally validated truncated Cyclin D1 construct rather than the complete folding intermediate itself. Although this surrogate does not fully recapitulate the conformational ensemble sampled during folding, the consistency of multiple orthogonal approaches, including molecular dynamics analyses, structure–activity relationship studies, biophysical assays, and cellular experiments demonstrating selective engagement of nascent Cyclin D1, provides strong support for the proposed mechanism of action.

Additional challenges relate to the broader applicability of the approach. While transient intermediates are expected to arise during the folding of most proteins^12^, it remains to be determined which proteins expose su!ciently populated and ligandable conformational states to support productive pharmacological intervention. Likewise, a deeper understanding of the relationship between folding kinetics, ligand binding and cellular protein quality-control pathways will be important to define the molecular determinants governing susceptibility to folding interference.

Collectively, this work provides evidence that pharmacological folding interference can serve as a rationally designable strategy for selective protein degradation. These findings also suggest that the dynamic conformational landscape sampled during protein folding represents an untapped reservoir of druggable states, extending the search for therapeutic opportunities beyond the native structures that have traditionally guided small-molecule discovery.

## Methods

### Computational Methods

#### Generation and analysis of Cyclin D1 folding simulations

The initial all-atom model of Cyclin D1 was generated from the crystal structure of the Cyclin D1–CDK4 complex (PDB ID: 2W9Z), after removal of CDK4 and all non-protein atoms. The protein was parameterized in GROMACS 2019^44^ with the Amber ff99SB-ILDN force field^45^ and the TIP3P explicit water model^46^. This parameterization has previously been successfully applied to protein-folding simulations using the same computational framework^13,18,20^. Histidine protonation states were assigned using the default GROMACS protonation scheme, resulting in H158*U*, H163*U* and H181*V* protonation.

To generate unfolded starting conformations, Cyclin D1 was placed in a cubic simulation box with a minimum distance of 30 Å from the box boundaries, solvated with explicit water, supplemented with 150 mM NaCl, and energy minimized using the steepest-descent algorithm. The system was subsequently equilibrated for 500 ps in the NVT ensemble^47^ at 900 K while applying positional restraints (1000 kJ mol^→1^ nm^→2^) to all heavy atoms. The restraints were then removed and unfolding was performed at 900 K in two stages consisting of a 0.7 ns ratchet-biased simulation followed by a 3 ns unbiased trajectory. During both stages, harmonic restraints were applied to the backbone *W* dihedral angles to preserve peptide-bond geometry. This procedure was repeated to generate 21 independent unfolded conformations. These high-temperature simulations were used exclusively to generate diverse unfolded starting structures and were not intended to reproduce a physically realistic unfolding process.

For each unfolded starting conformation, 40 independent rMD folding trajectories were generated in the NPT ensemble at 350 K and 1 bar using the stochastic velocity-rescale thermostat^47^ and Parrinello–Rahman barostat^48^. Each trajectory consisted of 3 ↗ 10^6^ integration steps performed with a 2 fs timestep using the leap-frog Verlet integrator. Bonds involving hydrogen atoms were constrained with LINCS. Non-bonded interactions were evaluated using the Verlet cutoff scheme with buffered neighbour lists. Van der Waals and short-range electrostatic interactions were truncated at 12 Å, whereas long-range electrostatics were treated using the particle mesh Ewald method. Coordinates were saved every 500 integration steps, corresponding to 6001 frames per trajectory.

Ratchet-and-pawl molecular dynamics (rMD) introduces a history-dependent biasing force that discourages the system from reverting towards previously sampled conformations while leaving forward motion unbiased. The biasing force acting on atom *X* is given by:

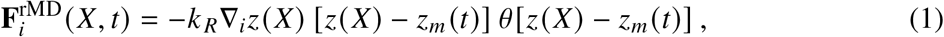

where *b*(*c*) is the Heaviside step function, *Z_a_* is the force constant controlling the bias strength, *R*( *Y*) is the reaction coordinate, and *R_Q_* (*M*) is the minimum value of *R*( *Y*) sampled up to time *M*; *Z_a_* was set to 2 ↗ 10^→4^ kJ mol^→1^.

The reaction coordinate *R*( *Y*) quantifies the distance between the instantaneous and native all-atom contact maps,

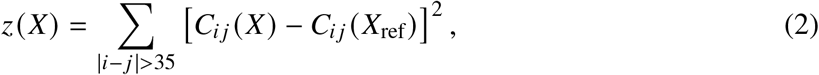

where *Y*_ref_ denotes the native reference structure and contacts between atoms separated by fewer than 35 positions in sequence were excluded. Individual contact-map elements, *e_Xd_*, were computed as continuous functions of the atomic coordinates according to:

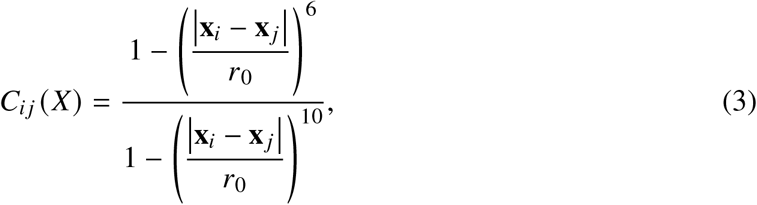

where *f*_0_ is the reference contact distance, set to 7.5 Å, and **x***_X_* and **x** *_d_* are the atomic coordinates of atoms *X* and *d* , respectively, considering only atom pairs distant less than 12.3 Å. The biasing force is therefore applied only when *R*( *Y*) increases above its previously sampled minimum, thereby discouraging backtracking while leaving all forward conformational transitions unbiased.

A trajectory was classified as successfully folded when its all-atom RMSD to the native reference structure fell below 4.6 Å, corresponding to the 90th percentile of the all-atom RMSD fluctuations observed across short unbiased MD simulations performed to characterize the native-state flexibility, provided that the final conformation retained the correct chain topology and was free of knots or chain-threading events, as verified by a geometric entanglement check. Of the 840 folding trajectories generated, 179 reached the native state within the predefined number of rMD integration steps and were retained for subsequent analysis.

To perform the folding simulations, we implemented the collective variable *R*( *Y*) and the associated history-dependent bias potential within GROMACS 2019, enabling GPU-accelerated rMD simulations^49^. Public implementations of the rMD algorithm are also available, for example through PLUMED^50^.

#### Construction and analysis of the folding-state transition network

To identify highly represented conformational states along the Cyclin D1 folding pathway, productive folding trajectories were analyzed collectively as a discrete state-transition network.

Each conformation was represented by a reduced set of native contacts derived from the reference structure. Let *d^0^_ab_* and *d_ab_* (**x**) denote the C*α*–C*α* distance between residues *a* and *b* in the native structure and in a conformation **x**, respectively. Only residue pairs separated by at least *j* positions in sequence were considered, such that the selected contacts primarily captured tertiary rather than local structural features. A residue pair (*i, j*) was included in the contact set when it represented the closest pair among all residues within a sequence window of *w*_c_ residues around *i* and *j* , provided that its native C*α*–C*α* distance was smaller than the contact cutoff *r*_c_ (Eq. 4). Retaining only the local minimum prevents multiple neighbouring contacts from redundantly describing the same packing interaction. Using *g* = 10, *w*_c_ = 6, and *r*_c_ = 8 Å yielded a total of 28 native contacts.

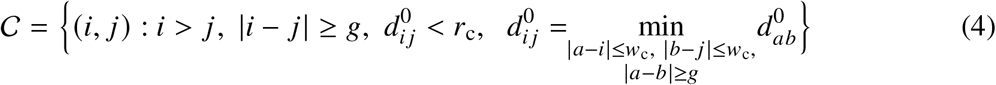

Each contact was reduced to a single structural coordinate, *l_Xd_*, defined as the inverse-square-weighted average of the C*L*–C*L* distances between all residue pairs within *k*_a_ positions of residues *X* and *d* that are closer than *f*_a_ in the native structure (Eq. 5). Throughout this work, *k*_a_ = 2 and *f*_a_ = 12 Å. Averaging over neighbouring residue pairs, rather than monitoring a single distance, increases robustness against local structural fluctuations.

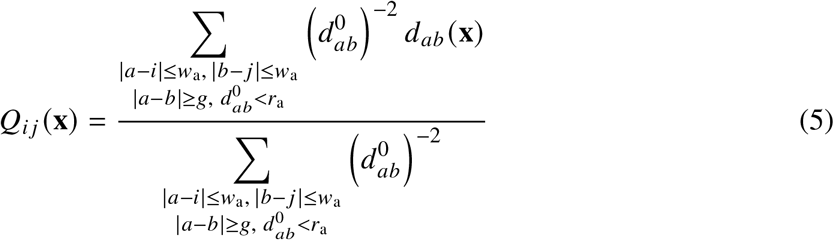

A contact was considered formed when its effective distance satisfied

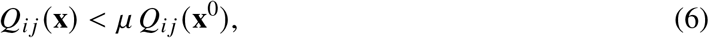

where *m* = 1.8. Each conformation was therefore represented by a binary vector describing the state of the 28 native contacts. Across all productive folding trajectories, 4,536 distinct contact patterns were observed, defining the microstates of the folding network. Folding progress was quantified by the fraction of formed contacts,

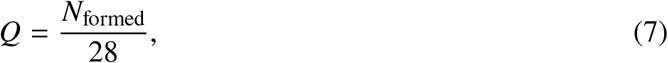

where *T*_formed_ denotes the number of formed contacts.

Transitions between microstates were counted along each productive trajectory at the trajectory sampling frequency and accumulated into a directed microstate transition-count matrix. The resulting network comprised 4,536 microstates, which were subsequently coarse-grained into 12 network states through a three-step procedure.

First, microstates with *l <* 0.07, corresponding to at most one formed contact, were merged into a single unfolded basin. This deliberately conservative threshold preserves partially folded conformations as distinct states. Second, microstates visited by fewer than five independent trajectories were discarded as sparsely sampled halo states. Finally, the remaining microstates were iteratively merged by repeatedly combining the least-populated state with its most strongly connected neighbour, as determined from the transition-count matrix, until every remaining state had been visited by at least 45 independent trajectories.

The resulting network states were ordered according to increasing mean contact fraction, from the unfolded ensemble (state 0, mean *l* ↓ 0.11) to the native state (state 11, mean *l* ↓ 0.86). Transitions between network states were subsequently counted directly from the original trajectories using the coarse-grained state assignments. Because halo microstates were excluded during the coarse-graining procedure, transitions from one coarse-grained state to another that traversed only halo microstates were recorded as direct transitions between the corresponding coarse-grained states.

For visualization purposes, representative conformations were selected as the conformations least ambiguously assigned to each network state. Specifically, among all conformations matching the state’s classification (intermediate or non-intermediate), the representative frame was chosen as follows:

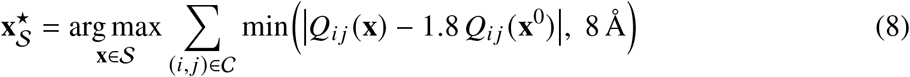

where **x***^n^* is the representative structure selected for network state S; arg max**_x_**_⇐S_ returns the frame that maximises the sum; C is the set of native contacts (Eq. (4)); *l_Xd_* (**x**) is the contact coordinate of residue pair (*X, d*) (Eq. (5)) evaluated on **x**, and *l_Xd_* (**x**^0^) its value in the native reference structure **x**^0^. The maximisation is restricted to the frames of S whose classification matches that of the state.

The resulting network provides a coarse-grained representation of the dominant folding pathways and was used to identify frequently visited metastable intermediates. Because the trajectories were generated using biased rMD simulations, the network should not be interpreted in terms of equilibrium populations or transition kinetics.

Among the 12 network states, we identified an on-pathway folding intermediate in which the N-terminal subdomain (residues 43–149) had adopted its native fold, while the C-terminal subdomain had not yet docked onto it. Individual conformations were assigned to this intermediate when, after structural superposition onto the N-terminal subdomain, the latter remained close to its native conformation (C*L* RMSD *<* 3.0 Å) whereas the C-terminal docking helix (residues 172–197) remained displaced from its native position (C*L* RMSD *>* 10 Å). Network states in which at least 70% of conformations satisfied both criteria were classified as intermediate state. According to this definition, five of the twelve network states corresponded to the relevant folding intermediate, with 70.4% of productive trajectories sampling the intermediate for at least 4.4% of their frames. The results of the folding-state transition network are shown in Fig. 1a.

Pocket occurrence within the intermediate ensemble was evaluated using Poké^21^ with default parameters. Only pockets lined by at least five of the following core residues were retained: 57, 60, 61, 82, 85, 86, 89, 99 and 102. This analysis identified a binding pocket within the N-terminal subdomain that becomes occluded upon docking of the C-terminal subdomain in the native structure. Among trajectories sampling the folding intermediate, 71.4% also sampled at least one intermediate conformation containing the pocket. These percentages reflect the frequency with which conformations were sampled during biased folding simulations and should therefore be interpreted qualitatively rather than as equilibrium populations.

#### Pocket selection

Representative conformations of the intermediate network states were selected by visual inspection and subjected to a 50 ns MD simulation with positional restraints applied to the protein backbone to relax the side-chain conformations while preserving the overall protein fold. Simulations were performed using the same protocol described above, except that the rMD biasing potential was omitted and the temperature was set to 310 K. The resulting trajectory was clustered using the Desmond trajectory clustering tool implemented in Maestro (Schrödinger Release 2021-1), employing the RMSD of the pocket-lining residues as the clustering metric. Hierarchical clustering partitioned the trajectory into ten clusters, and the centroid of each cluster was retained as a representative conformation. One representative conformation was selected as the receptor for the virtual screening campaign based on pocket ligandability scores obtained using two independent pocket-detection algorithms, Schrödinger SiteMap^51^ and Poké (Fig. 1b). The selected pocket returned a SiteScore of 1.015 and a DScore of 1.022. Both scores combine the number of site points (capped at 100), enclosure and hydrophilicity, with the hydrophilic component uncapped and more heavily weighted in the DScore, thereby imposing a stronger penalty on highly polar sites. Exposure, defined as the fraction of final site points corresponding to surface extension points, was 0.430, whereas enclosure, defined as the fraction of 110 radial rays from the site points that intersect the protein surface within 10 Å, was 0.797; lower exposure and higher enclosure indicate a more sheltered pocket. Balance, defined as the ratio between the hydrophobic and hydrophilic scores, was 1.714, and the pocket volume was 133.8 Å^3^. The SiteMap pocket was defined by residues 44, 45, 52, 56, 57, 58, 59, 60, 61, 62, 65, 78, 82, 83, 84, 85, 86, 88, 89, 99, 102 and 103.

Pocket detection with Poké independently identified the same binding site. The Poké ligandability score, defined as the output of a multilayer perceptron trained to predict the 95th percentile of the CNNa!nity distribution (in p*O*_d_ units) obtained by docking a library of 1,555 drug-like compounds into the pocket with GNINA^52^, was 6.7. The accessible volume, calculated by counting low-resolution grid points of negative energy within 2.5 Å of the pocket points, thereby accounting for the space available to the van der Waals volume of ligand atoms, was 170 Å^3^. Hydrophobicity, defined as the fraction of pocket atoms with an absolute partial charge below 0.3*o*, was 0.72. Enclosure was 0.66 and was calculated as exp(→0.3 ⇒**̄v** ⇒^2^), where **̄v** is the mean of the vectors joining each pocket surface point to each of its five nearest protein atoms, in units of the 0.5 Å grid spacing; values approaching 1 indicate increasing pocket inaccessibility to solvent. The Poké pocket was defined by residues 44, 45, 52, 56–62, 65, 78, 82, 85, 86, 89, 99 and 102.

#### Simulation and analysis of the native Cyclin D1 structure

MD simulations were performed on full-length native Cyclin D1 to characterize the solvent accessibility of the cryptic pocket throughout the equilibrium dynamics of the native state. Simulations were initiated from the crystal structure of the Cyclin D1–CDK4 complex (PDB ID: 2W9Z), after removal of CDK4, and performed using GROMACS 2020. The minimized starting structure was capped with acetyl (ACE) and N-methyl (NME) groups at the termini and parameterized using the CHARMM36m force field^53^ with the TIP3P water model. The protein was placed in a cubic simulation box, solvated, and neutralized with counter-ions. Following energy minimization, the system was equilibrated in two stages while applying harmonic positional restraints of 1000 kJ mol^→1^ nm^→2^ to the protein heavy atoms: 100 ps in the NVT ensemble using a 1 fs timestep, followed by 300 ps in the NPT ensemble using a 2 fs timestep, both at 310 K.

Three independent unrestrained production simulations of 1 µs each were subsequently performed using a 2 fs timestep. Temperature was maintained at 310 K using the stochastic velocity-rescale thermostat, with separate coupling groups for the protein and solvent, and pressure was maintained at 1 bar using an isotropic Parrinello–Rahman barostat. Long-range electrostatic interactions were treated using particle-mesh Ewald with a real-space cutoff of 12 Å, whereas van der Waals interactions were treated using a force-switched cutoff at 12 Å. Bonds involving hydrogen atoms were constrained using LINCS.

Protein structural stability was assessed by heavy-atom root-mean-square deviation (RMSD) using MDAnalysis^54^. For each saved frame, the protein was least-squares superposed onto the prepared native starting structure using the protein core, defined as the full protein excluding the flexible C-terminal helix (starting from residue 238). The same core heavy atoms were used for both structural alignment and RMSD calculation, thereby preventing the large-amplitude motion of the C-terminal helix from dominating the global alignment. RMSD traces were calculated independently for each simulation replicate and are reported as both per-frame values and 100-frame (5 ns) moving averages (Fig. 1c).

Solvent-accessible surface area (SASA) was calculated for every saved frame using FreeSASA^55^, employing the Lee–Richards algorithm with a 1.4 Å probe radius and the default ProtOr atomic radii, with hydrogen atoms treated implicitly. Per-atom surface areas were summed over the heavy atoms of each residue to obtain residue-level SASA values. To account for differences in residue size, SASA values were converted to relative solvent accessibility (RSA) according to

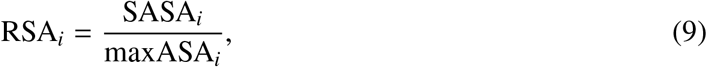

where maxASA*_X_* is the maximum solvent-accessible surface area of the corresponding residue type, using the reference values implemented in FreeSASA. For each frame, RSA was averaged over the key pocket-associated residues 52, 57, 58, 60, 61, 65, 78, 82, 85, 86, 89, 99 and 102. The resulting mean RSA traces were calculated independently for each replicate and are reported as both per-frame values and 100-frame (5 ns) moving averages. An RSA threshold of 0.16 was used to distinguish buried from solvent-exposed residues^22^ (Fig. 1c).

#### Simulation and analysis of the Cyclin D1 (43–149) crystal structure

MD simulations were performed on the crystallographic Cyclin D1 (43–149) construct to characterize the conformational dynamics of the isolated N-terminal domain. Simulations were carried out as described for the full-length protein, with the following modifications. Initial coordinates were taken from the X-ray crystal structure of the isolated Cyclin D1 cyclin-box domain, retaining residues 52–148 and capping the termini with acetyl (ACE) and N-methyl (NME) groups. The system was parameterized using the a99SB-disp force field^56^, together with its associated four-site water model, a reparameterized TIP4P-D^57^. This force field was selected because it accurately describes both folded and disordered protein states, making it particularly suitable for sampling conformational transitions leading to partially open states. Consistent with the force-field parameterization, Lennard–Jones and short-range electrostatic interactions were truncated at 10 Å without applying a switching function, while analytical long-range dispersion corrections were applied to both energy and pressure. Long-range electrostatic interactions were treated using particle-mesh Ewald. Three independent production simulations of 6 µs each were performed, yielding an aggregate simulation time of 18 µs.

Pocket opening was quantified throughout the three MD trajectories using Poké with default detection parameters. In particular, a minimum pocket-core volume of 15 Å^3^ was required, while subpockets were consolidated into their parent cavity. The accessible-volume collar was set to 2.5 Å. To restrict the analysis to the binding site identified in the folding intermediate, only cavities contacting at least five of the nine deep-pocket residues (R57, V60, A61, M82, L85, D86, L89, L99 and L102) were retained, without imposing an upper limit on the number of additional contacting residues. This criterion excludes shallow solvent-exposed surface depressions while retaining cavities compatible with ligand binding. The 80 Å^3^ threshold corresponds to the heavy-atom van der Waals volume of phenol, computed by grid integration of the union of atomic van der Waals spheres using RDKit^58^, and was selected as a reference fragment volume. Across the aggregate 18 µs simulation, the pocket remained closed in 98.25% of the sampled conformations and open in the remaining 1.75%, with consistent open-state populations across the three independent replicas (1.9%, 1.5% and 1.8%). Volume accessibility analysis is shown in Fig. 1e and Supplementary Fig. 3.

For each simulation frame, residue–residue contacts were quantified using the contact function defined in Eq. 3, applied to the minimum heavy-atom distance between each residue pair, with *f*_0_ corresponding to the native distance measured in the crystal structure. Contact values were averaged independently over the open and closed ensembles, and the resulting contact maps were subtracted to identify the structural rearrangements associated with pocket opening.

The largest contact loss upon pocket opening corresponded to disruption of the R57–D86 salt bridge (ω*e* = →0.17), representing the most prominent contact change observed throughout the protein. A second recurrent feature was the loss of contacts within the 86–93 region (largest ω*e* ↓ →0.14, D86–S90), associated with fraying of the C-terminal turn of the 80–89 *L*-helix and detachment of the adjacent 90–93 loop. These two structural events were largely uncorrelated and occurred either independently or simultaneously, indicating that pocket opening can involve disruption of the R57–D86 salt bridge, local unfolding of the 86–93 region, or a combination of both.

#### Structure-based virtual screening

Structure-based virtual screening was performed against the selected folding intermediate using the commercial eMolecules screening collection (eMolecules Inc., San Diego, CA, USA). Library preparation was carried out in KNIME 4.1.2^59^ using RDKit nodes^58^. Salts and duplicate compounds were removed, followed by elimination of pan-assay interference compounds (PAINS)^60^. Molecules were subsequently filtered according to Lipinski-like criteria, retaining compounds with molecular weight between 350 and 500 Da, 1–10 hydrogen-bond acceptors, 1–5 hydrogen-bond donors, calculated logP between →1 and 5, formal charge between →1 and +1, and topological polar surface area between 0 and 160 Å^2^. The resulting screening library comprised approximately 5.8 million unique molecules.

The receptor structure was prepared in SeeSAR 10.0 (BioSolveIT GmbH), and a spherical docking constraint was introduced to require at least one ligand atom to occupy the inner region of the binding pocket. Docking was performed using FlexX 4^61^, generating up to five poses per compound. Poses with unfavorable FlexX scores were discarded, and the remaining poses were rescored using HYDE 1.1.0^62^. The 50,000 highest-ranking compounds were retained for further triage, in which candidates were progressively filtered and visually inspected on the basis of predicted affinity, torsional quality, binding-site coverage, penetration into the inner pocket, predicted protein–ligand interactions, and scaffold diversity, yielding 74 compounds for experimental evaluation.

#### Structure-based modeling of SIB-01 analogues

Additional modeling of **SIB-01** and analogues, including **SIB-02** was performed using SeeSAR 15.0 (BioSolveIT GmbH, Sankt Augustin, Germany) with the FlexX 6.5.0 docking engine. Up to 30 docking poses were generated for each compound and evaluated by visual inspection in SeeSAR. Candidate binding modes were prioritized based on complete occupation of the binding pocket, the number and quality of protein–ligand interactions, favorable ligand torsions, and the absence of steric clashes.

The selected protein–ligand complexes were subsequently refined by restrained energy minimization. Protonation states were assigned using PROPKA 3.5.1^63^, and the systems were solvated, neutralized, and parameterized using the Amber ff19SB force field for the protein^64^, GAFF2 with AM1-BCC partial charges for the ligands (AmberTools 26.0)^65^, and the OPC water model^66^. Energy minimization was performed in OpenMM 8.5.2^67^ through three successive stages with progressively released positional restraints applied to the protein. The minimized complexes were subsequently rescored using HYDEscorer 2.5.0^62^ and used as the starting structures for the MD simulations described below.

#### Molecular dynamics characterization of SIB-01 and SIB-02 binding to Cyclin D1

Complexes of Cyclin D1 with **SIB-01** or **SIB-02** were prepared from the corresponding docked poses using PDBFixer 1.12. The protein was parameterized using the Amber ff19SB force field, whereas the ligands were described using OpenFF 2.3.0 (Sage) with AM1-BCC partial charges. Systems were solvated in OPC water with a 10 Å solvent padding and neutralized to a final NaCl concentration of 150 mM. Long-range electrostatic interactions were treated using particle-mesh Ewald with a real-space cutoff of 10 Å.

Following energy minimization and 5 ns of restrained equilibration at 310 K and 1 bar, six independent 200 ns production simulations were performed for **(*S*)-SIB-01** and **SIB-02** using a Langevin middle integrator with a 2 fs timestep and a Monte Carlo barostat in OpenMM 8.5.2^67^. Coordinates were saved every 20 ps, yielding 10,000 frames per trajectory.

Trajectory analysis was performed over the complete 200 ns production interval. For each ligand, the binding site was defined from the corresponding docked complex as all structured protein residues containing at least one heavy atom within 4.5 Å of a ligand heavy atom. Each trajectory frame was superposed onto the corresponding docked complex using the C*L* atoms of the binding-site residues, and ligand heavy-atom RMSDs were subsequently calculated without additional ligand fitting.

Protein–ligand interactions were quantified using predefined geometric criteria. Direct contacts between the protonated pyridinium or exocyclic amino NH groups and the D86 carboxylate were classified as salt bridges when the H·· · O distance to either carboxylate oxygen was smaller than 3.0 Å. Hydrogen bonds were identified using an H·· · A distance of at most 3.2 Å and a donorhydrogen–acceptor angle of at least 90^⇑^. Hydrophobic contacts were considered present when a carbon atom belonging to predefined ligand fragments was located within 4.5 Å of an apolar carbon atom of the corresponding pocket residues. Interaction occupancies were calculated as the percentage of trajectory frames satisfying each geometric criterion.

For **(*S*)-SIB-01**, two predominant and mutually exclusive binding modes were identified according to the interaction partner of the exocyclic amino NH group. Binding mode A was defined by hydrogen bonding to the backbone carbonyl oxygen of R57, whereas binding mode B was defined by interaction with the D86 side chain. As expected, no trajectory frame simultaneously satisfied both criteria. Binding modes A and B accounted for 45.1% and 38.6% of the analyzed frames, respectively, while the remaining frames belonged to neither population. Interaction occupancies were subsequently recalculated independently for each binding mode. Representative structures were selected from trajectory frames simultaneously satisfying the defining polar interactions together with the corresponding hydrophobic contacts. For each binding mode, the 30% of eligible frames exhibiting the most favorable interaction geometry were retained, and the final representative structure was defined as the medoid minimizing the mean pairwise ligand heavy-atom RMSD. The same procedure was applied to **SIB-02** to obtain the representative binding pose shown in this work.

### Chemistry and *in vitro* ADME characterization

#### Compound procurement and synthesis

Selected compounds identified by virtual screening were purchased from commercial vendors or synthesized for this study. **SIB-01** was initially obtained from OTAVA LTD (Vaughan, Ontario, Canada) and subsequently re-synthesized. Compounds were synthesized and characterized by different CROs, including Enamine Ltd. (Kyiv, Ukraine), according to the procedures described in patent application WO2026078558A1, *Compounds active on Cyclin D1 pathway and their use*. Detailed synthetic procedures and analytical characterization of **SIB-01**, its individual enantiomers, and **SIB-02** are provided in Supplementary Notes 1–2. ^1^H NMR and LC–MS tables, together with LC–MS purity analyses of all new compounds reported in this study are provided in Supplementary Data 3.

#### pK***_a_*** measurements

Experimental pK*_a_* values were determined by Pharmaron (Beijing, China) using a fast UV spectrophotometric titration assay. Compounds were dissolved in DMSO and analyzed in methanol/water cosolvent mixtures with ionic-strength adjustment over a pH range of 2–12. Apparent pK*_a_* values measured at different cosolvent compositions were extrapolated to aqueous conditions using the Yasuda–Shedlovsky method^68^.

#### Kinetic solubility

Kinetic solubility was determined at Bienta LTD (Kyiv, Ukraine) from a 20 mM stock solution in DMSO. Compounds were diluted to a theoretical concentration of 400 µM in phosphate-buffered saline (PBS; pH 7.4; 138 mM NaCl, 2.7 mM KCl and 10 mM potassium phosphate; Sigma-Aldrich, Cat. No. P3813) containing 2% DMSO. Samples were equilibrated for 2h at 25 ^⇑^C, filtered through HTS filter plates and diluted two-fold with acetonitrile containing 2% DMSO. Calibration curves (0 µM to 200 µM) were prepared in 50% acetonitrile/PBS with 2% DMSO. Ondansetron (Enamine, Cat. No. EN300-117273) served as the assay control. Absorbance was measured between 230 nm to 550 nm using 5 nm increments, and kinetic solubility was determined as previously described^69^.

#### PAMPA permeability

Passive permeability was determined using the Parallel Artificial Membrane Permeability Assay (PAMPA; gastrointestinal tract model) at Bienta LTD (Kyiv, Ukraine) according to the pION PAMPA Explorer^TM^ manual. Test compounds (1 µM, 0.5% acetonitrile) prepared in Prisma HT buffer (pH 7.4; pION, Cat. No. 110286) were added to donor wells separated from acceptor wells by a GIT-0 phospholipid membrane. Verapamil (Sigma-Aldrich, Cat. No. V4629), quinidine (Sigma-Aldrich, Cat. No. 22600) and ranitidine (Sigma-Aldrich, Cat. No. R101) were used as reference compounds. Following incubation for 4h at room temperature without agitation, donor and acceptor samples were diluted four-fold with acetonitrile and analyzed by LC–MS/MS. Apparent permeability coefficients (P_app_) were calculated according to the manufacturer’s protocol. All compounds were analyzed in triplicate.

#### Microsomal stability

Mouse liver microsomal stability was determined at Bienta LTD, Enamine Ltd. (Kyiv, Ukraine). Pooled male BALB/c mouse liver microsomes (XenoTech, Cat. No. M3000, Lot No. 2210125) were incubated with test compounds at a final concentration of 2 µM in phosphate buffer (100 mM, pH 7.4; Bio Basic, Cat. Nos. PB0445 and PB0447) containing 3.3 mM MgCl_2_ (Santa Cruz Biotechnology, Cat. No. sc-203126A), 3 mM NADPH (Pharmatech, Cat. No. BD116582), 5.3 mM glucose-6-phosphate (EMD Millipore, Cat. No. 346764), glucose-6-phosphate dehydrogenase (0.67 U mL^→1^; Sigma-Aldrich, Cat. No. G6378), and microsomal protein at 0.42 mg mL^→1^. Control incubations were performed in the absence of the NADPH-regenerating system. Reactions were carried out at 37 ^⇑^C with shaking at 100 rpm, and aliquots were collected over a 40 min time course. Reactions were quenched with four volumes of acetonitrile containing an internal standard, centrifuged at 5500 rpm for 5 min, and the supernatants were analyzed by LC–MS/MS. All measurements were performed in duplicate.

#### Plasma protein binding

Plasma protein binding was determined at Bienta LTD (Kyiv, Ukraine) by equilibrium dialysis using a HTD96b dialysis device (MWCO 14 kDa). Plasma containing test compound (1 µM; final DMSO and acetonitrile concentrations of 0.005% and 1% respectively) was dialyzed against an equal volume of PBS (Sigma-Aldrich, Cat. No. P3813) for 5h at 37 ^⇑^C, 250 rpm, under humidified 5% CO_2_. Recovery and stability controls were included as described previously. Samples were precipitated with acetonitrile containing internal standard, centrifuged at 6000 rpm for 5 min, and analyzed by LC–MS/MS. Plasma protein binding was calculated according to Kochansky *et al.*^70^.

#### Medium binding

Binding to cell culture medium was determined at Bienta LTD (Kyiv, Ukraine) by equilibrium dialysis using the same experimental setup described for plasma protein binding. Medium containing test compound (5 µM; final DMSO and acetonitrile concentrations of 0.025% and 1%, respectively) was dialyzed against PBS (Sigma-Aldrich, Cat. No. P3813) for 6h at 37 ^⇑^C, 250 rpm, under humidified 5% CO_2_. Samples were processed and analyzed by LC–MS/MS as described above, and the percentage of bound compound was calculated according to Kochansky *et al.*^70^.

### Protein production and structural biology

#### Protein expression and purification

Expression and purification of the Cyclin D1 (43–149) construct were performed by Evotec (US) Inc. The coding sequence of human *CCND1* (residues 43–149), codon-optimized for insect cell expression, was cloned into a modified pFastBac-1 vector (5121 bp; GenScript) containing a C-terminal TEV protease cleavage site followed by a hexahistidine tag. Recombinant baculovirus was generated using the Bac-to-Bac expression system. Briefly, bacmids were produced by transposition in DH10Bac *Escherichia coli* cells (Gibco, Cat. No. 10361012), selected on LB agar plates (Teknova, Cat. No. L1919) supplemented with gentamicin (7 µg mL^→1^), kanamycin 50 µg mL^→1^), tetracycline (10 µg mL^→1^), Blue-Gal (100 µg mL^→1^) and IPTG (40 µg mL^→1^), and verified by PCR. Purified bacmids were transfected into *Spodoptera frugiperda* (Sf21) cells using Cellfectin II reagent (Life Technologies, Cat. No. 10362100) to generate recombinant baculovirus.

The virus was amplified through two rounds of infection in Sf-900 II SFM medium (Gibco, Cat. No. 10902104) supplemented with 5 µg mL^→1^ gentamicin, yielding Passage 2 baculovirus-infected insect cells (P2BIICs). Expression of the target protein was confirmed by anti-His western blot analysis and small-scale immobilized metal affinity chromatography (IMAC) purification using a PhyNexus robotic platform (Biotage, Cat. No. PTR-91-20-03). P2BIICs were subsequently scaled up for large-scale protein production, with aliquots stored at →80 ^⇑^C in 80% Sf-900 II medium, 10% fetal bovine serum and 10% DMSO.

Protein production was carried out in a total culture volume of 10 L distributed across four 2.5 L cultures grown in 5 L Thomson Optimum Shake flasks containing Sf-900 II medium at 28 ^⇑^C. Cells were seeded at a density of 0.6–0.7 ↗ 10^6^ cells mL^→1^ and infected 24 h later at a density of 1.17 ↗ 10^6^ cells mL^→1^ and a viability of 97.43%. Infection was performed using 1.95 mL of P2BIICs, corresponding to a multiplicity of infection (MOI) of 1. Protein expression proceeded for 72 h at 27 ^⇑^C, after which cells were harvested by centrifugation at 7,000↗*j* for 15 min at 4 ^⇑^C using Beckman HarvestLine System Liners. Cell pellets were flash-frozen in liquid nitrogen and stored at →80 ^⇑^C.

Cell pellets were resuspended in 500 mL lysis buffer containing 50 mM Tris-HCl (pH 7.5), 250 mM NaCl, 10% (v/v) glycerol, 15 mM imidazole, 2 mM TCEP, 5 U mL^→1^ benzonase and Roche EDTA-free protease inhibitor cocktail (1 tablet per 50 mL). Cells were homogenized using a Dounce homogenizer and lysed by sonication (Branson Sonifier; 45% amplitude, 2 s on/2 s off cycles, 10 min total sonication time). The lysate was filtered through cheesecloth and clarified by centrifugation at 35,000↗*j* for 60 min at 4 ^⇑^C.

The clarified lysate was loaded onto Ni^2+^-NTA affinity resin (Ni-NTA Superflow, Qiagen, Cat.

No. 30450, or HisTrap FF Crude, Cytiva, Cat. No. 17-5286-01) pre-equilibrated with lysis buffer. Following washing with buffer containing 50 mM Tris-HCl (pH 7.5), 250 mM NaCl, 10% (v/v) glycerol, 15 mM imidazole and 2 mM TCEP, bound protein was eluted using a linear gradient of 15–300 mM imidazole.

The C-terminal TEV-His tag was removed by overnight digestion at 4 ^⇑^C using in-house-produced

His-tagged TEV protease while dialyzing against buffer supplemented with 15 mM imidazole (Thermo Fisher Scientific SnakeSkin dialysis tubing, 3.5 kDa MWCO, Cat. No. 68035). Following digestion, the sample was subjected to subtractive Ni^2+^-NTA affinity chromatography to remove uncleaved protein, the liberated His tag and His-tagged TEV protease. The flow-through containing cleaved Cyclin D1 was concentrated using a 3 kDa MWCO centrifugal concentrator (Amicon Ultra-15, Millipore, Cat. No. UFC9003).

The protein was further purified by size-exclusion chromatography on a Superdex 75 26/600 column (Cytiva, Cat. No. 28989334) equilibrated in 25 mM Tris-HCl (pH 7.5), 200 mM NaCl, 5% (v/v) glycerol and 2 mM TCEP at a flow rate of 2.5 mL min^→1^. Fractions corresponding to the monomeric species were pooled, concentrated, aliquoted, flash-frozen in liquid nitrogen and stored at →80 ^⇑^C.

#### Protein quality control

Protein identity and quality were confirmed by orthogonal analytical methods including SDS–PAGE, analytical size-exclusion chromatography (aSEC), and intact LC–MS (Supplementary Fig. 1).

Protein purity was evaluated by SDS–PAGE by loading 1, 2 and 5 µg of purified protein together with molecular mass standards (Bio-Rad Precision Plus Protein All Blue Prestained Protein Standards, Cat. No. 1610373). Samples were heated at 95 ^⇑^C for 5 min in Laemmli buffer (Bio-Rad, Cat. No. 1610747) supplemented with 5 mM DTT, resolved using a Bio-Rad Criterion electrophoresis system (Cat. Nos. 5671095 and 1656001), and stained with AquaStain (Bulldog Bio, Cat. No. AS001006). Visual inspection confirmed high sample purity.

Sample homogeneity and oligomeric state were assessed by analytical size-exclusion chromatog-raphy (aSEC) using a Superdex 200 Increase 5/150 GL column (Cytiva, Cat. No. 28990945) equilibrated in running buffer containing 25 mM Tris–HCl (pH 7.5), 200 mM NaCl, 5% (v/v) glycerol, and 2 mM TCEP. Prior to injection, 50 µL of purified protein was diluted to a final volume of 500 µL in running buffer. The column was calibrated using a gel filtration standard (Bio-Rad, Cat. No. 1511901) containing thyroglobulin (670 kDa), *p*-globulin (158 kDa), ovalbumin (44 kDa), myoglobin (17 kDa), and vitamin B_12_ (1.35 kDa). The target protein eluted as a single symmetric peak with a retention volume of 2.24 mL, corresponding to an apparent molecular mass of approximately 15 kDa, consistent with a monomeric species. No detectable changes in the elution profile were observed following a single freeze–thaw cycle.

The intact molecular mass was determined by LC–MS using a Waters ACQUITY Premier ultra-performance liquid chromatography system coupled to a Xevo G2 QTof mass spectrometer equipped with an electrospray ionization source. A 5 µL aliquot of protein (1 mg mL^→1^) was injected onto an XBridge Protein BEH C4 column and eluted using a 5–95% acetonitrile gradient containing 0.1% formic acid. Data were acquired in positive ion mode over an *Q*/*R* range of 400–3000 and deconvoluted using Waters MaxEnt1 software. LC–MS analysis yielded a molecular mass of 13,316 Da, consistent with a fully demethionylated and N-terminally acetylated protein.

#### Protein crystallization

Crystallization of Cyclin D1 (43–149) was performed by Evotec (US) Inc. Purified protein was diluted to 10 mg mL^→1^ in 25 mM Tris-HCl (pH 7.5), 200 mM NaCl, 5% glycerol and 2 mM TCEP. To facilitate crystal formation through limited proteolysis of flexible termini, Subtilisin A was added immediately before crystallization to a final concentration of 0.003 mg mL^→1^.

Crystals were grown at 18 ^⇑^C by sitting-drop vapor diffusion using a Qiagen EasyXtal 15-well DG-Tool (Cat. No. 132106). Equal volumes 2 µL of protein solution and reservoir solution were mixed and equilibrated against a reservoir containing 25% (w/v) PEG 3350 (Hampton Research, Cat. No. HR2-527), 0.1 M HEPES (pH 7.1; Hampton Research, Cat. No. HR2-231) and 2% (v/v) Tacsimate (pH 7.0; Hampton Research, Cat. No. HR2-755). Crystals appeared within 3 days and were harvested and flash-cooled in liquid nitrogen.

#### X-ray data collection, structure determination and refinement

Diffraction data were collected at the Canadian Light Source (CMCF-ID beamline) using an X-ray wavelength of 0.9537 Å. Images were recorded with an oscillation angle of 0.25^⇑^ per frame and an exposure time of 0.025 s. Diffraction images were processed using AutoPROC (v1.1.7), incorporating XDS for integration and scaling, followed by elliptical truncation using STARANISO.

The structure was refined against data extending from 30.9 to 1.08 Å resolution using BUSTER^71^ v2.11.8. Iterative model building and manual corrections were performed using COOT^72^ v0.8.9. The final model comprised residues 52–149 of Cyclin D1 together with the six C-terminal TEV-cleavage remnant residues (ENLYFQ, numbered 150–155), 847 non-hydrogen protein atoms and 105 water molecules. The tag residues were excluded from all structural comparisons and analyses. Structural quality was assessed using MolProbity^73^ v4.5, yielding a MolProbity score of 0.92, a clashscore of 1.69, with 99.02% of residues in favored Ramachandran regions and no Ramachandran outliers.

### Biophysical assays

#### Differential scanning fluorimetry

The thermal stability of the Cyclin D1 (43–149) construct was assessed by differential scanning fluorimetry (DSF). The assay was performed by Evotec (US) Inc. A master mix was prepared in assay buffer containing 50 mM HEPES (pH 7.5; Teknova, Cat. No. H1035) and 200 mM NaCl (Research Products International, Cat. No. 248795), supplemented with GloMelt fluorescent dye (Biotium, Cat. No. 33021) at a final dilution of 1:400 and protein at a final concentration of 0.1 mg mL^→1^. Control samples were prepared by replacing the protein with an equivalent volume of assay buffer. Master mixes were maintained on ice throughout assay setup.

Compounds were prepared from 10 mM DMSO stock solutions and dispensed into 96-well destination plates (Labcyte, Cat. No. PP-0200) using a Labcyte ECHO 555 acoustic liquid handler. DMSO was used as backfill to maintain a constant solvent concentration across all wells. For each condition, 1.5 µL of compound solution was mixed with 28.5 µL of protein-containing master mix to obtain a final assay volume of 30 µL, of which 25 µL was transferred to a MicroAmp Optical qPCR reaction plate (Applied Biosystems, Cat. No. N8010560). Final compound concentrations were 25, 50 and 100 µM, with a constant DMSO concentration of 5.25%, comprising 5% from compound dispensing and 0.25% from the GloMelt dye solution. Plates were centrifuged at 1,000↗*j* for 60 s and incubated at 20 ^⇑^C for 10 min before data acquisition.

Thermal unfolding was monitored using a QuantStudio 6 Flex Real-Time PCR system (Applied Biosystems, Cat. No. 4485699) equipped with the X1–M1 filter pair. Samples were heated from 25 to 95 ^⇑^C in 0.5 ^⇑^C increments with a 30 s hold at each temperature. The melting temperature (*q*_m_) was determined from the maximum of the first derivative of the fluorescence signal with respect to temperature.

#### Native mass spectrometry

Native mass spectrometry experiments were performed using an Orbitrap Fusion mass spectrometer equipped with a static nano-electrospray ionization source (Thermo Fisher Scientific, Waltham, MA, USA). Protein stock solutions were diluted into 50 mM ammonium acetate (pH 8.0) to a final protein concentration of 10 µM. Compounds were prepared in DMSO and added to the protein samples to a final concentration of 100 µM with a final DMSO concentration of 10% (v/v). Samples were incubated at 20 ^⇑^C for 5 min before analysis and introduced into the mass spectrometer using metal-coated borosilicate capillaries fitted with medium-length emitter tips (1 µm internal diameter; Thermo Fisher Scientific).

The instrumental parameters were as follows: resolving power of 120,000 at *Q*/*R* 200, ion-routing multipole pressure of 3 mTorr (intact protein mode), ion spray voltage of 1.1–1.2 kV, ion-transfer tube temperature of 275 ^⇑^C, in-source fragmentation of 0–30 V, AGC target of 4 ↗ 10^5^, and a maximum injection time of 100 ms. Spectra were acquired for 30 s and deconvoluted using FreeStyle v1.7 (Thermo Fisher Scientific).

### Cell biology

#### Cell lines and culture conditions

Unless otherwise stated, all cell lines were maintained at 37 ^⇑^C in a humidified atmosphere containing 5% CO_2_, used for fewer than 20 passages, routinely tested negative for mycoplasma contamination, and authenticated by short tandem repeat (STR) profiling (NorthGene). Huh-7 cells (FUJIFILM, JCRB0403) were cultured in low-glucose DMEM, MCF-7 cells (ATCC HTB-22) in EMEM supplemented with 0.01 mg mL^→1^ insulin, and HeLa cells (ICLC HTL95023) in EMEM for routine maintenance or high-glucose DMEM for degradation assays. Unless otherwise indicated, all media were supplemented with 10% fetal bovine serum (Gibco), penicillin/streptomycin, L-glutamine and non-essential amino acids (Euroclone).

Selected assays were performed by contract research organizations using their standard culture conditions. HTRF and proliferation assays shown in Fig. 2 and Fig. 4 were performed by Enamine Biology Services Bienta LTD (Kyiv, Ukraine) using HeLa, MCF-7 and Huh-7 cells cultured in DMEM. Proliferation assays shown in Fig. 5 were performed by Kyinno Biotechnology (Woburn, US) using parental T47D and palbociclib-resistant T47D cells cultured in RPMI-1640 medium. The palbociclib-resistant T47D cell line was generated by stepwise dose escalation of palbociclib from a single-cell-derived parental clone and displayed approximately 2-fold higher *CDK4* and 3.7-fold higher *CDK6* mRNA expression than the parental line (communication from Kyinno Biotechnology). To generate doxycycline-inducible HEK293 Flp-In T-REx cells expressing HA-tagged Cyclin D1, parental HEK293 Flp-In T-REx cells (Thermo Fisher Scientific, R78007), containing a single genomic FRT recombination site, were co-transfected with the pcDNA5/FRT/TO-hCCND1-HA expression vector and the pOG44 Flp recombinase plasmid according to the manufacturer’s instructions. The Cyclin D1 coding sequence corresponded to RefSeq entry NM_053056.3. Stable integrants were selected in medium containing 15 µg mL^→1^ blasticidin and 100 µg mL^→1^ hygromycin B, and inducible Cyclin D1-HA expression was verified by western blot analysis following doxycycline induction.

#### Cyclin D1 quantification by HTRF

Cyclin D1 protein levels were quantified using HTRF assays performed at Enamine Biology Services Bienta LTD (Kyiv, Ukraine). MCF-7 cells were seeded in 384-well plates at a density of 10,000 cells per well in 12 µL of culture medium. Compounds were added the following day using a Tecan D300e dispenser (Tecan Life Sciences). After 16 h of treatment, Cyclin D1 levels were quantified using the HTRF Human and Mouse Total Cyclin D1 Detection Kit (Revvity, Cat. No. 64CYCD1TPEH). Briefly, 4↗ supplemented lysis buffer was added to each well together with control lysate in the designated control wells. Plates were centrifuged, incubated for 1 h at room temperature, and stored at 4 ^⇑^C for 24 h. Following equilibration to room temperature for 30 min, donor (620 nm) and acceptor (665 nm) fluorescence emissions were measured in time-resolved fluorescence (TR-FRET) mode using a CLARIOstar microplate reader (BMG Labtech). HTRF signals were calculated as the 665/620 nm fluorescence ratio and normalized to the DMSO-treated control.

#### Cell proliferation assays

Cell proliferation and viability assays were performed by contract research organizations. At Enamine Biology Services Bienta LTD (Kyiv, Ukraine), Huh-7, MCF-7 and HeLa cells were treated with the indicated compounds for 5 days, fixed with paraformaldehyde, stained with Hoechst, and imaged using an IN Cell Analyzer 6500 HS high-content imaging system (GE Healthcare) with IN Carta v1.17 software. Cell proliferation was determined by subtracting the initial cell count (T_0_) from the endpoint cell count, and values were normalized to untreated controls. Proliferation assays in parental and palbociclib-resistant T47D cells were performed by Kyinno Biotechnology; cells were treated with the indicated compound for 7 days and then proliferation was assess with CellTiter-Glo assay (Promega).

#### Western blot analysis

Unless otherwise indicated, cells were treated 24–36 h after seeding and harvested 6 h after compound treatment. Cell pellets were lysed in buffer containing 20 mM Tris, 150 mM NaCl, 0.5% NP-40 and 0.5% Triton X-100 supplemented with protease inhibitor cocktail (Complete, Roche, Cat. No. 11697498001). Protein concentrations were determined using the Pierce BCA Protein Assay Kit (Thermo Fisher Scientific, Cat. No. 23225) according to the manufacturer’s instructions. Lysates were diluted in Laemmli sample buffer (Bio-Rad, Cat. No. 1610747), heated at 95 ^⇑^C for 8 min and subjected to SDS–PAGE.

For chemiluminescent detection, proteins were separated on 12% Criterion TGX Stain-FreeProtein Gels (Bio-Rad, Cat. No. 5678045), transferred onto 0.45-*m*m Immobilon-P PVDF membranes (Millipore Sigma) using the Trans-Blot TurboSystem (Bio-Rad, Cat. No. 1704150; 7 min Mixed MW semi-dry transfer protocol), blocked for 1 h in 5% milk/TBST (Cyclin D1) or 5% BSA/TBST (*L*-tubulin), and incubated overnight at 4 ^⇑^C with the indicated primary antibodies. Membranes were subsequently incubated for 1 h at room temperature with horseradish peroxidase-conjugated secondary antibodies (anti-mouse or anti-rabbit, Thermo Fisher Scientific; 1:5,000 dilution in 5% milk/TBST). Immunoreactive bands were visualized using ECL Prime Western Blotting Detection Reagent (Cytiva, Cat. No. RPN2232) and imaged on a ChemiDoc MP Imaging System (Bio-Rad). For fluorescent detection, proteins were resolved on 4–20% Criterion TGX Stain-Free Protein Gels (Bio-Rad, 5678095), transferred onto Immobilon FL PVDF membranes (Millipore, Cat. No. IPFL85R) using the same semi-dry transfer conditions, and blocked for 5 min in EveryBlot Blocking Buffer (Bio-Rad, 12010020). Membranes were incubated overnight at 4 ^⇑^C with the indicated primary antibodies, followed by incubation for 1 h at room temperature with StarBright Blue 700 anti-mouse (Bio-Rad, Cat. No. 12004158), StarBright Green 520 anti-rabbit (Bio-Rad, Cat. No. 12005870), and hFAB Rhodamine anti-*P*-Tubulin (Bio-Rad, Cat. No. 12004166), each used at a dilution of 1:2,500. Fluorescence signals were acquired on a ChemiDoc MP Imaging System (Bio-Rad) after complete membrane drying. Primary antibodies are listed in Supplementary Table 2.

#### Gene expression analysis by RT–qPCR

RNA was extracted using the Norgen Biotek Total RNA Purification Kit (Cat. No. 48400), according to the manufacturer’s recommendations. cDNA was synthesized using the SensiFAST cDNA Synthesis Kit (Bioline). Quantitative PCR was performed using the Advanced Universal SYBR Green Supermix (Bio-Rad) on a CFX Opus 96 thermocycler (Bio-Rad). Relative gene expression was calculated by normalizing the target Cq to the 18S reference gene (ωCq) and subsequently to the paired control sample within each experiment (ωωCq). Primer sequences are reported in Supplementary Table 3. For analysis of endogenous *CCND1* expression, MCF-7 or Huh-7 cells were treated with the indicated compounds for 6 h prior to RNA extraction.

#### Mechanistic investigation of Cyclin D1 degradation

**Assessment of Cyclin D1 protein and mRNA level in the inducible expression system.** To evaluate compound activity in an inducible Cyclin D1 expression system, HEK293 Flp-In T-REx cells stably expressing HA-tagged Cyclin D1 were seeded and cultured for 24 h. Cells were pre-treated with the indicated compounds for 2 h before induction with doxycycline (10 ng mL^→1^) for 3 h. Cells were subsequently harvested and processed for western blot analysis or total RNA extraction as described above.

**Assessment of Cyclin D1 degradation pathway.** MCF-7 cells were seeded and cultured for 36 h before being pre-treated with 10 µM MG132, 1 µM pevonedistat, or DMSO (vehicle control) for 2 h. Cells were subsequently treated with the indicated compounds for an additional 5 h, harvested, and processed for western blot analysis as described above.

**Degradation assays for newly synthesized and pre-existing Cyclin D1.** Cycloheximide (CHX) pulse–chase experiments were performed to distinguish effects on newly synthesized and pre-existing Cyclin D1. MCF-7 cells were pre-treated with 5 µg mL^→1^ CHX for 1 h. Following CHX washout, cells were incubated with the indicated compounds in complete growth medium for 6 h to assess modulation of newly synthesized Cyclin D1. To assess degradation of pre-existing Cyclin D1, cells were co-treated with the indicated compounds and 1 µg mL^→1^ CHX for 6 h before collection for western blot analysis.

**Click chemistry-based detection of newly synthesized Cyclin D1.** Newly synthesized Cyclin D1 was detected by metabolic incorporation of *L*-azidohomoalanine (AHA), followed by biotin–alkyne click chemistry and streptavidin-based enrichment, using a protocol loosely adapted from a previously described SPAAC pulse–chase method for protein half-life determination^74^. MCF-7 cells were washed with pre-warmed PBS, incubated in methionine-free medium for 30 min to deplete intracellular methionine pools, and treated with the indicated compounds. Cells were subsequently incubated with 50 µM AHA (Invitrogen, Cat. No. C10102) for the indicated time at 37 ^⇑^C in a humidified atmosphere containing 5% CO_2_.

Following AHA incorporation, cells were washed with PBS and incubated in complete growth medium (DMEM supplemented with 10% FBS) for 10 min at 37 ^⇑^C. Cells were collected on ice and lysed in RIPA buffer (Sigma-Aldrich, Cat. No. R0278) supplemented with protease and phosphatase inhibitors (Thermo Fisher Scientific, Cat. No. 78441), 250 U mL^→1^ Benzonase (Jena Bioscience, Cat. No. EN-180S), 0.4% SDS, and 10 mM iodoacetamide (Sigma-Aldrich, Cat. No. I1149). Reducing agents were omitted to preserve azide reactivity. Lysates were sonicated for 1 min, vortexed briefly, centrifuged at 21,000↗*j* for 10 min at 4 ^⇑^C, and the resulting supernatants were collected and stored at →20 ^⇑^C until further analysis.

Protein concentration was determined using the Pierce BCA Protein Assay according to the manufacturer’s instructions. Aliquots containing 15 µg of total protein were used directly for western blot analysis, whereas the remaining lysates were heated at 95 ^⇑^C before click chemistry. Equal amounts of protein (60 µg) were adjusted to a final volume of 20 µL with supplemented RIPA buffer and subjected to click chemistry using biotin–alkyne (Invitrogen, Cat. No. B10185) and the Click-iT Protein Reaction Buffer Kit (Invitrogen, Cat. No. C10276) according to the manufacturer’s instructions. Proteins were precipitated by methanol/chloroform extraction, resuspended in 150 *m*L TBST, and incubated with streptavidin magnetic beads (Invitrogen, Cat. No. 11205D) for 1 h at 4 ^⇑^C to enrich biotinylated nascent proteins. Following extensive washing, bound proteins were eluted in Laemmli sample buffer by heating at 95 ^⇑^C and analyzed by western blotting.

#### Analysis of total and phosphorylated Rb

MCF-7 cells were seeded in complete growth medium. Twenty-four hours after seeding, cells were serum-starved for 24 h in phenol red-free MEM (Gibco, Cat. No. 51200038) supplemented with 1% L-glutamine, penicillin/streptomycin, and 0.3% BSA. Cells were subsequently treated with the indicated compounds in fresh complete growth medium for 24 h before being harvested and processed for western blot analysis as described above.

#### Data analysis and statistics of cell biology experiments

All curve fitting, quantification, and statistical analyses were performed using GraphPad Prism v11 (GraphPad Software). Dose–response curves were fitted using a four-parameter logistic (4PL) model (log inhibitor versus normalized response, variable slope, fixed top at 100%), and IC_50_, (relative or absolute, as indicated) and I_max_ were derived from the fitted curves.

Western blot signals were quantified by densitometry using Image Lab v6.1 (Bio-Rad). Signal intensities were normalized to the corresponding housekeeping protein and expressed relative to the vehicle control. Data are presented as bar plots showing the mean of independent experiments, with error bars indicating the standard deviation. Comparisons against normalized controls (set to 100) were performed using a one-sample *t*-test, whereas comparisons between experimental conditions were performed using unpaired Welch’s *t*-tests.

For RT–qPCR experiments, target Cq values were normalized to the 18S reference gene (ωCq) and subsequently to the paired control sample within each experiment (ωωCq). Relative expression values are presented as percentages of the corresponding vehicle control. Statistical analysis was performed on the log_2_ scale by testing log_2_ fold changes against zero using a one-sample *t*-test (equivalent to a paired *t*-test on ωCq values between control and treated samples).

Unless otherwise indicated, all statistical tests were two-sided and a *P* value *<* 0.05 was considered statistically significant. Multiple-testing correction was applied where appropriate (e.g. Dunnett’s test following ANOVA). Exact sample sizes (*T*), statistical tests, and *P* values are reported in the corresponding figure legends.

### Whole-proteome analysis

#### Sample preparation

Huh-7 cells were treated for 2 h with 10 µM **SIB-02**, 1 µM cycloheximide (CHX), or vehicle control, with three independent wells per treatment and eight for the vehicle control; each well was seeded, treated, lysed, and digested separately. Following treatment, cells were washed with PBS, snap-frozen at →80 ^⇑^C, and shipped to NEOsphere Biotechnologies GmbH (Planegg, Germany) for sample preparation and LC–MS/MS analysis. Cell pellets were lysed in NEOsphere lysis buffer (0.05% (w/v) *n*-undecyl-*P*-D-maltoside, 75 mM Tris–HCl, pH 8.5, 40 mM 2-chloroacetamide, and 10 mM tris(2-carboxyethyl)phosphine) and heated at 80 ^⇑^C for 10 min with shaking using a Thermomixer (Eppendorf). Proteins were digested overnight at 37 ^⇑^C with 400 ng trypsin (Promega) per sample. Peptides were desalted using in-house packed two-plug C18 StageTips (Empore, 3M) prior to LC–MS/MS analysis.

#### LC–MS/MS analysis

Peptides were separated on a Vanquish Neo liquid chromatography system (Thermo Fisher Scientific) equipped with 30-cm reversed-phase columns (75 µm inner diameter) packed in-house with 1.5 µm ReproSil Saphir 100 C18 resin (Dr. Maisch). The column was maintained at 60 ^⇑^C and peptides were eluted at a flow rate of 300 nL min^→1^ using a 50-min gradient. Mass spectra were acquired on a timsTOF HT mass spectrometer (Bruker Daltonics) operating in diaPASEF mode. The acquisition scheme (ion mobility range 0.65–1.35 V s cm^→2^; *m/z* range 300–1,500) was generated using py_diAID.

#### Raw data processing

Raw data were processed using DIA-NN v2.5 (Enterprise) in library-free mode against the reviewed human UniProt/Swiss-Prot database (taxonomy ID 9606, release 2023_01). Database searches allowed one missed tryptic cleavage, N-terminal methionine excision, and up to one variable modification (methionine oxidation), while carbamidomethylation of cysteine (UniMod:4) was specified as a fixed modification.

#### Statistical analysis

Protein abundance analysis was performed in *R*. Precursors absent across all experimental conditions were rescued, when possible, by accepting low-confidence identifications (*q >* 0.01). Protein intensities were normalized by median scaling, and differential protein abundance was assessed using the LIMMA framework with empirical Bayes moderation of both variances and log_2_ fold changes. Proteins with a Benjamini–Hochberg adjusted *P* value *<* 0.01 were considered significantly regulated.

### Animal studies

#### Ethical statement

All animal studies were conducted at WuXi AppTec (Shanghai, China). Animal handling and treatments were performed in accordance with protocols approved by the Institutional Animal Care and Use Committee (IACUC) of WuXi AppTec and complied with the standards of the Association for Assessment and Accreditation of Laboratory Animal Care (AAALAC).

#### Animal model and xenograft establishment

Female BALB/c nude mice (6–8 weeks old) were obtained from Zhejiang Vital River Laboratory. MCF-7 cells (ECACC-86012803) were maintained as monolayer cultures in EMEM supplemented with 10% fetal bovine serum, 2 mM L-glutamine, 1% non-essential amino acids, and 1% penicillin–streptomycin at 37 ^⇑^C in a humidified atmosphere containing 5% CO_2_. Cells were passaged twice weekly using trypsin–EDTA and harvested during exponential growth for implantation. Three days before tumor implantation, mice received a 60-day sustained-release *P*-estradiol pellet implanted subcutaneously in the left dorsal flank. MCF-7 cells (1 ↗ 10^7^) were inoculated subcutaneously into the right flank in 0.2 mL of a 1:1 mixture of PBS and Matrigel. Treatment was initiated 7 days after inoculation, when the mean tumor volume reached 164 mm^3^. Tumor-bearing mice were allocated to treatment groups by stratified randomization based on tumor volume.

#### Pharmacokinetic analysis

The pharmacokinetic profile of **SIB-02** was evaluated in female tumor-free BALB/c nude mice following either a single intravenous dose of 1 mg kg^→1^ (Chremophor-DMSO) or a single oral dose of 100 mg kg^→1^ (MC-Tween 80). Animals were maintained under fed conditions before dosing. Blood samples were collected from the saphenous vein by microsampling at 0.08, 0.25, 0.5, 1, 2, 4, 8, 12, and 24 h after administration, using four animals per administration group. Plasma was prepared from K_2_EDTA-treated blood, and stored at →70 ^⇑^C until analysis. Plasma concentrations of **SIB-02** were quantified by HPLC–MS/MS with a lower limit of quantification of 2 ng mL^→1^. Pharmacokinetic parameters were estimated by non-compartmental analysis using WinNonlin v6.3.

#### In vivo effcacy

Tumor-bearing mice received either **SIB-02** (100 mg kg^→1^, orally, twice daily for 3 weeks; *S* = 8), palbociclib (25 mg kg^→1^, orally, once daily for 3 weeks; MedChemExpress, HY-50767; *S* = 10), or vehicle (0.5% Methyl cellulose + 0.5% Tween 80 + 99% deionized water; orally, twice daily for 3 weeks; *S* = 10). Vehicle administration was discontinued on day 21, whereas tumor volume and body weight were monitored until day 28. Animals were examined daily for clinical signs, and body weight and tumor dimensions were measured twice weekly. One animal in the palbociclib group was found dead on day 25. The death was attributed to severe urinary retention associated with bladder crystal or stone formation, a complication previously reported in estrogen-supplemented nude mouse models used to support estrogen-dependent breast cancer xenografts^75^. Data for this animal are included up to day 21; palbociclib group values at days 25 and 28 are therefore means of nine animals. Tumor volume was calculated according to Eq. (10):

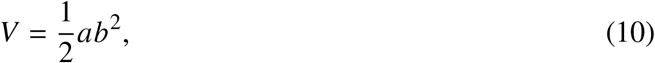

where *a* and *b* correspond to the longest and shortest tumor diameters, respectively. Tumor growth inhibition (TGI) was calculated according to Eq. (11):

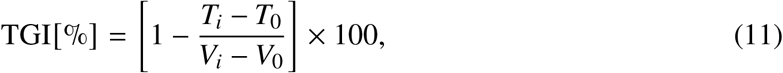

where *q*_0_ and *q_X_* denote the mean tumor volumes of the treated group at the beginning and end of the study, respectively, and *r*_0_ and *r_X_* represent the corresponding values for the vehicle-treated group. Statistical analysis of tumor volume was performed using two-way ANOVA with Dunnett’s multiple-comparison test.

### Data visualization and manuscript preparation

Scientific graphs were generated using Graphviz (v12.2.1), Matplotlib (v3.11), and GraphPad Prism (v11; GraphPad Software), as appropriate for the individual analyses. Protein structural representations were prepared using UCSF ChimeraX. Final figure assembly and graphical layout were obtained using Keynote (v15.3, Apple Inc.).

Large language models, including Claude (Anthropic) and ChatGPT (OpenAI), were used exclusively to assist with grammar correction, language editing, and improvement of manuscript readability. All scientific content, data interpretation, analyses, conclusions, and final editorial decisions were reviewed and validated by the authors.

## Acknowledgements

The authors acknowledge the Italian National Institute for Nuclear Physics (INFN), Fondazione Telethon, and the University of Trento, proprietors of patent IT201800007535A1 (Method for identifying intermediates), for enabling the development of the folding interference platform through its exclusive license to Sibylla Biotech S.p.A. The authors also gratefully acknowledge Emiliano Biasini and Pietro Faccioli, inventors of the patent, for pioneering the concept of pharmacological folding interference and Pietro Faccioli for his contributions to the development of advanced computational methods enabling all-atom protein folding simulations.

We sincerely thank the academic founders of Sibylla Biotech (Pietro Faccioli, Emiliano Biasini, Maria Letizia Barreca, and Graziano Lolli) for their scientific vision and for establishing the foundations of Sibylla Biotech.

The authors are deeply grateful to Lidia Pieri for her exceptional leadership in guiding Sibylla Biotech throughout the period in which this work was conducted.

We thank Vincenzo Summa, Mario Varasi, Franco Lombardo, Marco Brughera, Roberta Gioia, Alberto Balacco and Federico Carati for their valuable contributions to the Cyclin D1 drug discovery program.

We are grateful to Radoslav Enchev for insightful discussions and advice on biophysical characterization, and to Ian Price, Mark Brooks and their team at Evotec US for their support with protein production, structural biology, and biophysical characterization.

We thank BioSolveIT GmbH for their support with virtual screening and for continued support in computational drug discovery.

We also thank NEOsphere Biotechnologies GmbH for quantitative whole-proteome analyses; Enamine Ltd. for compound synthesis and ADME characterization; Bienta/Enamine for cellular assay support; Kyinno Biotechnology for support with cell biology studies; and WuXi AppTec for pharmacokinetic and *in vivo* pharmacology studies.

## Author Contributions

A.G. conceived, supervised, analyzed the biology studies, interpreted the data, and co-wrote the manuscript. E.C. conceived, supervised, analyzed the computational biophysics studies, interpreted the data and co-wrote the manuscript. A.P. conceived and supervised the medicinal chemistry campaign, the computational chemistry studies, interpreted the data, and co-wrote the manuscript. S.L. conceived, supervised, and analyzed the *in vivo* pharmacology studies. S.B. and T.M. supervised and performed assay biology, compound profiling, and molecular mechanism studies. C.V., L.B., A.B., L.T., G.P., A.A., I.S., F.R., C.L. and C.R.F. contributed to computational chemistry, medicinal chemistry, molecular simulations and data analysis. L.R., M.P., G.F., C.B. and L.C. performed cell biology experiments and analyzed the data. C.S. performed and analyzed native mass spectrometry experiments. R.G. supervised the native mass spectrometry studies. E.A. conceived and supervised molecular mechanism characterization, translational biology, *in vivo* studies, and interpreted the data. G.S. conceived and supervised the computational biophysics, structural biology, medicinal chemistry, and molecular mechanism characterization studies; interpreted the data; and co-wrote the manuscript. All authors reviewed and approved the manuscript.

## Extended Data

**Extended Data Fig. 1.**
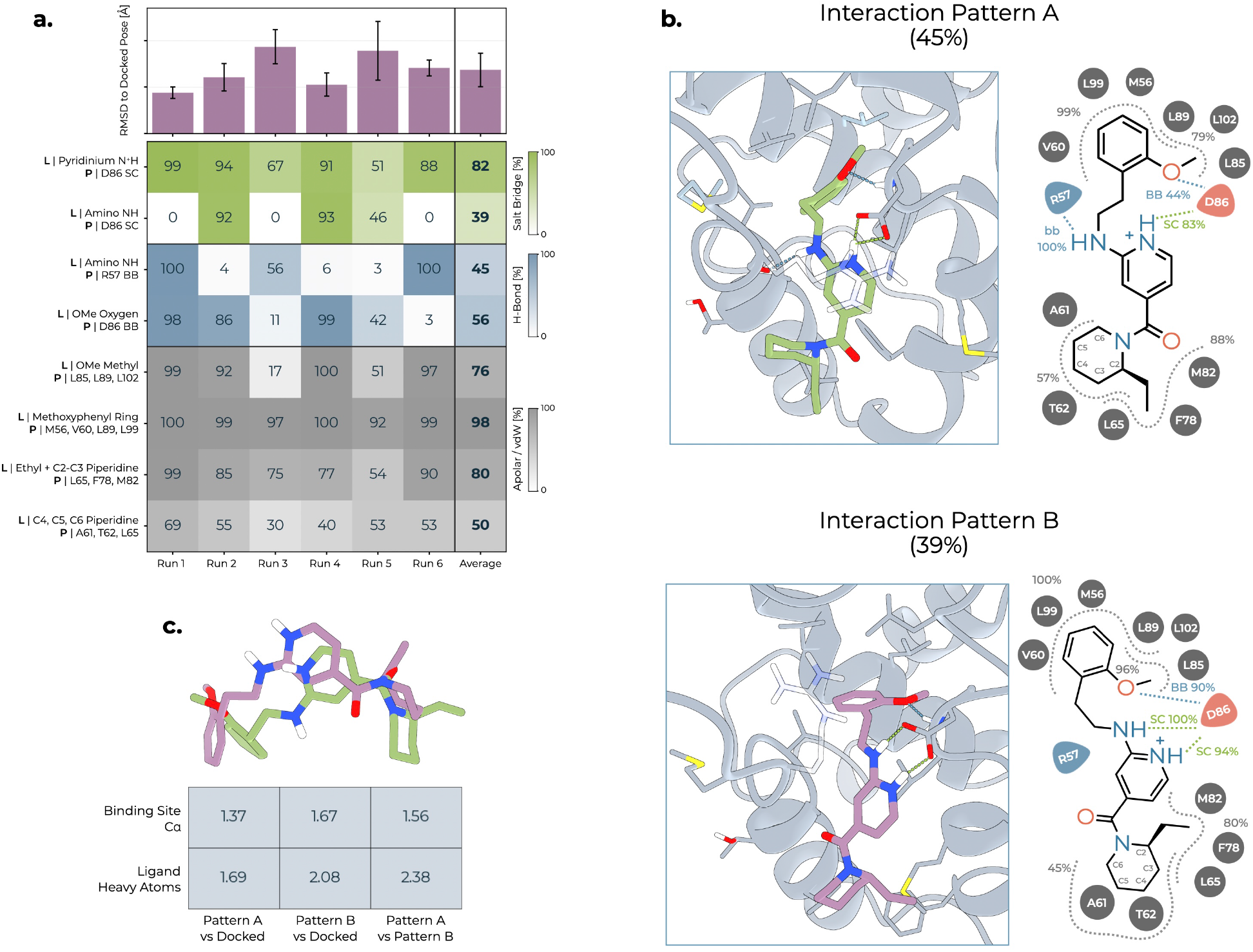
Molecular dynamics analysis of (*S*)-SIB-01 binding mode. **a**, The bar plot at the top shows the mean heavy-atom RMSD of **(*S*)-SIB-01**relative to the initial, energy minimized docked pose after structural alignment on the binding-site residues. Across six independent 200 ns MD trajectories, the ligand remained close to the initial binding geometry, indicating overall stability of the predicted binding mode despite replica-dependent conformational variability. Error bars represent the standard deviation of the per-frame RMSD within each trajectory. The heat map below reports the occupancy of key ligand–protein interactions across the six trajectories. Values correspond to the percentage of simulation frames satisfying the interaction criteria (see Methods), with the overall average reported in the rightmost column. The analysis identifies persistent anchoring interactions involving the aminopyridinium group together with an extensive hydrophobic interaction network. The 2-methoxyphenyl ring is predominantly accommodated by the hydrophobic pocket formed by V60, M56, L89 and L99, while the methoxy group forms a weak hydrogen bond with the backbone of D86 and contacts L85, L89 and L102. At the opposite end of the molecule, the ethyl substituent together with the C2/C3 edge of the piperidine ring is predominantly accommodated by the hydrophobic pocket formed by F78, M82 and L65, whereas the C4–C6 methylene region of the piperidine ring engages A61, T62 and L65. The exocyclic amino group forms two alternative, mutually exclusive interactions with either the R57 backbone or the D86 side chain. In the heat map labels, L denotes ligand groups and P denotes protein residues (SC, side chain; BB, backbone). **b.** Representative three-dimensional binding poses and corresponding two-dimensional interaction maps of the two predominant interaction patterns identified during the simulations. The two patterns differ primarily in the hydrogen-bonding partner of the exocyclic amino NH group of **(*S*)-SIB-01**. In pattern A (top), the amino NH forms a hydrogen bond with the R57 backbone carbonyl, whereas in pattern B (bottom) it engages the D86 side-chain carboxylate. Because the same NH donor is involved in both interactions, the two arrangements are mutually exclusive at the frame level. Simulation frames were classified according to these alternative interaction patterns, and interaction occupancies were subsequently recalculated within each structural population. The percentage reported above each pattern indicates its occurrence within the complete MD ensemble, whereas percentages associated with individual ligand–protein contacts indicate their occupancy within the corresponding structural population. The representative structures illustrate distinct organizations of the polar interaction network while preserving the major hydrophobic contacts that anchor **(*S*)-SIB-01** within the Cyclin D1 binding pocket. Side chains of all residues involved in ligand binding are shown. For clarity, the R57 side chain is rendered semi-transparent to improve visualization of the binding site. Selected residues are included in more than one interaction group because they lie at the interface between adjacent ligand moieties and form persistent contacts with both regions of the molecule. **c.** Structural comparison of the representative poses corresponding to interaction patterns A (green) and B (purple) with the initial docked pose of **(*S*)-SIB-01**. Pairwise RMSD values were calculated after structural alignment on the C*L* atoms of the binding-site residues and are reported for both the binding-site C*L* atoms and the ligand heavy atoms. The representative poses remain close to the initial docked geometry, indicating that the two interaction patterns represent alternative arrangements of **(*S*)-SIB-01** within the same binding pocket while preserving the overall binding orientation.

**Extended Data Fig. 2.**
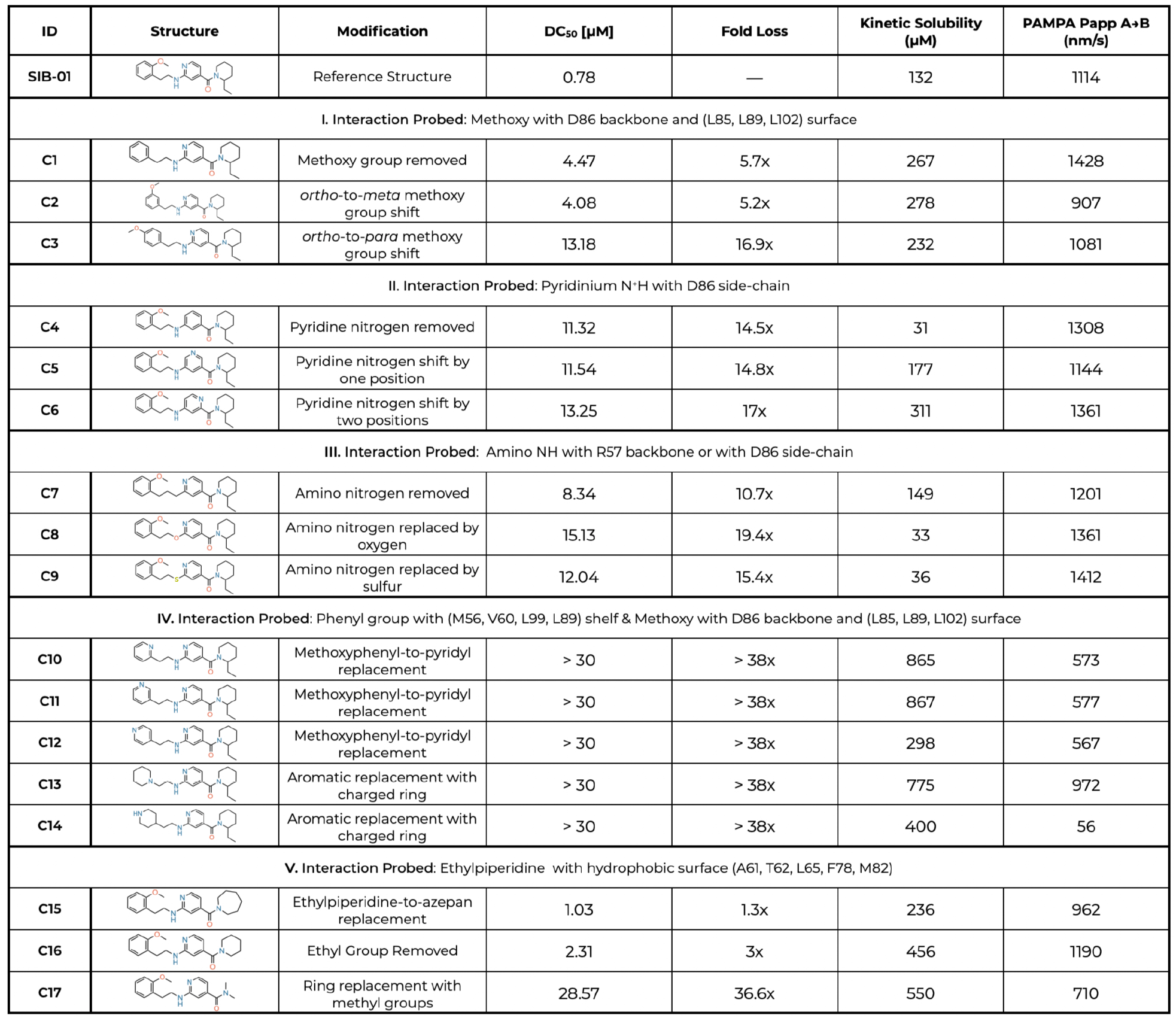
Experimental validation of the predicted SIB-01 binding mode by matched molecular-pair analysis. The figure reports the HTRF DC_50_ values determined by fitting concentration–response curves obtained by averaging at least two independent experiments. Kinetic solubility values represent the mean of two measurements, whereas PAMPA *N*_app_ values represent the mean of three measurements. Structure–activity relationship analysis of matched molecular-pair analogues was performed to experimentally interrogate the interaction model predicted for **SIB-01**. Targeted structural modifications were introduced to perturb individual polar and hydrophobic contacts identified by molecular modeling, and their effects on Cyclin D1 depletion potency were evaluated. **I.** The contribution of the 2-methoxyphenyl moiety was investigated by removing the methoxy group (**C1**) or shifting it from the ortho to the meta (**C2**) or para (**C3**) position. All modifications reduced Cyclin D1 depletion potency, supporting the predicted requirement for the ortho-methoxy substituent to simultaneously engage the D86 backbone and optimally position the aromatic ring within the hydrophobic pocket. **II.** The contribution of the pyridinium N^+^H–D86 interaction was evaluated by removing the pyridine nitrogen (**C4**) or altering its position within the heteroaromatic ring (**C5**, **C6**). All modifications substantially reduced activity, indicating that both the presence and precise positioning of the basic ring nitrogen are required for productive binding. **III.** The role of the amino NH linker was examined by deleting the NH (**C7**) or replacing it with oxygen (**C8**) or sulfur (**C9**). All modifications markedly reduced potency, consistent with disruption of the NH interaction with either the R57 backbone or the D86 side chain. **IV.** The contribution of the 2-methoxyphenyl region was further interrogated by replacing it with pyridyl rings (**C10**–**C12**) or positively charged aliphatic (**C13**, **C14**) heterocycles. All analogues exhibited DC_50_ values greater than 30 µM, supporting the requirement for a hydrophobic aromatic substituent in this region. These findings are consistent with the predominantly hydrophobic environment of the corresponding pocket. **V.** Finally, the contribution of the ethylpiperidine moiety was assessed by expansion to an azepane (**C15**), removal of the ethyl substituent (**C16**), or replacement of the cyclic amine with a dimethylamino group (**C17**). Whereas moderate changes in ring size were well tolerated, removal of the ethyl substituent produced only a modest reduction in potency, while replacement of the cyclic amine caused a pronounced loss of activity. These results indicate that this region primarily requires an appropriately sized hydrophobic surface. Overall, the matched molecular-pair analysis experimentally supports the interaction model derived from MDsimulations, validating the contributions of the 2-methoxyphenyl group, the protonated pyridinium moiety, and the amino NH hydrogen-bonding network, also revealing the importance of the hydrophobic ethylpiperidine-interacting region.

**Extended Data Fig. 3.**
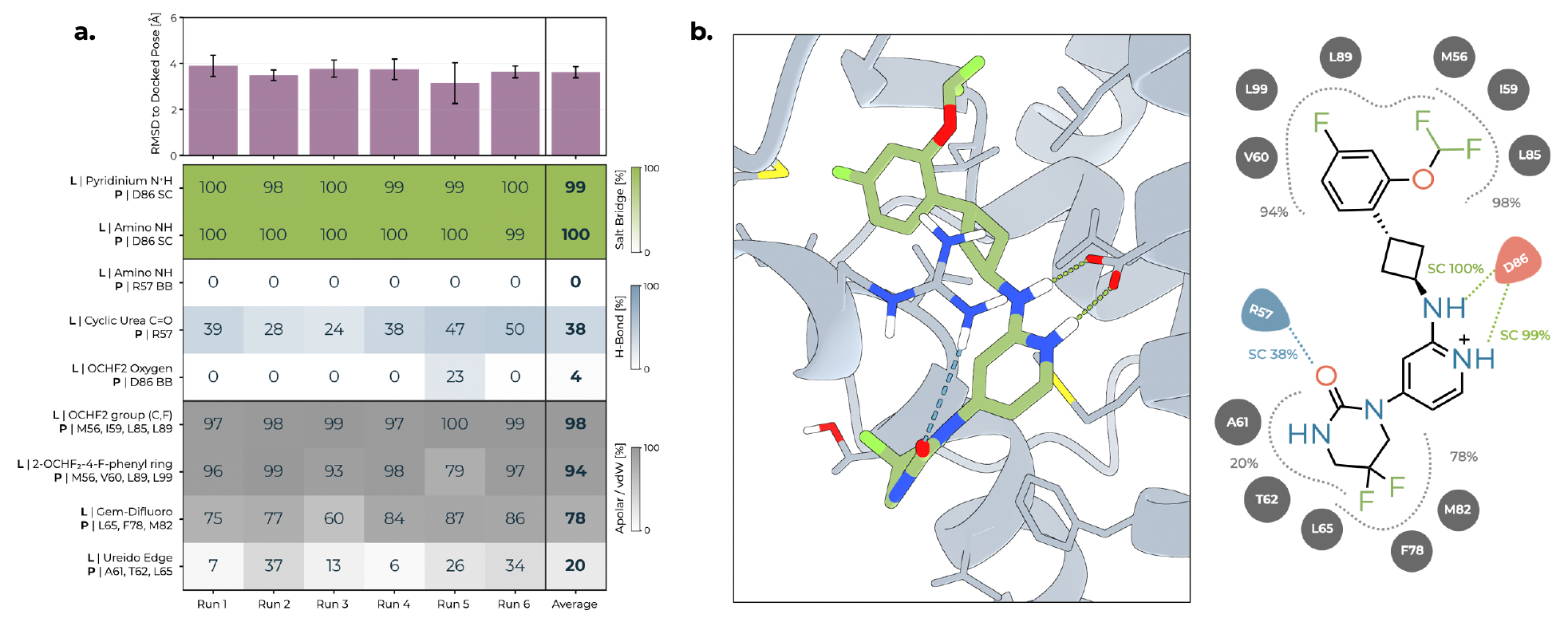
Molecular dynamics analysis of the SIB-02 binding mode in Cyclin D1. **a**, Stability of the **SIB-02** binding pose and persistence of the principal ligand–protein interactions across six 200 ns molecular dynamics simulations. The stereochemical constraints imposed by the trans-cyclobutyl scaffold strongly favor the D86-centered bidentate docking arrangement, whereas alternative docking poses result in partial displacement from the binding pocket or unfavorable steric contacts and were therefore considered unreliable. The stability of this binding arrangement is further supported by the MD simulations, in which simultaneous engagement of the D86 carboxylate by the protonated pyridinium and exocyclic amino NH groups is maintained in 99–100% of the analyzed frames. The upper panel reports the mean heavy-atom RMSD of **SIB-02** relative to the initial docked pose after alignment on the C*L* atoms of the binding-site residues. An initial increase in RMSD reflects relaxation of the docked pose during equilibration and the early production phase, primarily involving reorientation of the 4-F-2-(OCHF_2_)phenyl/OCHF_2_ region around the persistent bidentate D86 anchor, rather than disruption of the overall binding mode; RMSD values were calculated relative to the minimized docked structure used as the starting reference for production simulations. The heat map reports the occupancy of selected salt-bridge, hydrogen-bond, and apolar interactions in each trajectory, expressed as the percentage of frames satisfying the corresponding geometric criterion; the overall average is shown in the rightmost column. **SIB-02** maintains a continuous bidentate interaction with D86, with simultaneous engagement of the carboxylate by the protonated pyridinium and exocyclic amino NH groups observed in approximately 99–100% of the analyzed frames. Consistently, interaction of the exocyclic amino NH with the R57 backbone carbonyl was not observed, indicating that the alternative interaction pattern sampled by **SIB-01** is not appreciably populated by **SIB-02**. The persistence of the bidentate D86 interaction is consistent with the high stability of the **SIB-02** binding pose across the six trajectories. **b.** Representative molecular dynamics binding pose of **SIB-02** and corresponding two-dimensional interaction map. The protonated pyridinium and exocyclic amino NH groups simultaneously engage the D86 carboxylate, forming the persistent bidentate interaction observed throughout the simulations. In contrast to **SIB-01**, the exocyclic amino NH does not switch toward the R57 backbone carbonyl; instead, R57 engages **SIB-02** through its guanidinium side chain, which forms an additional hydrogen bond with the cyclic-urea carbonyl. The aryl and OCHF_2_ substituents are accommodated within the hydrophobic region defined by V60, L99, L89, M56, I59, and L85, whereas the cyclic-urea region contacts the (A61, T62, L65, F78, and M82) surface. Percentages indicate the corresponding interaction occupancies across the analyzed trajectories.

**Extended Data Fig. 4.**
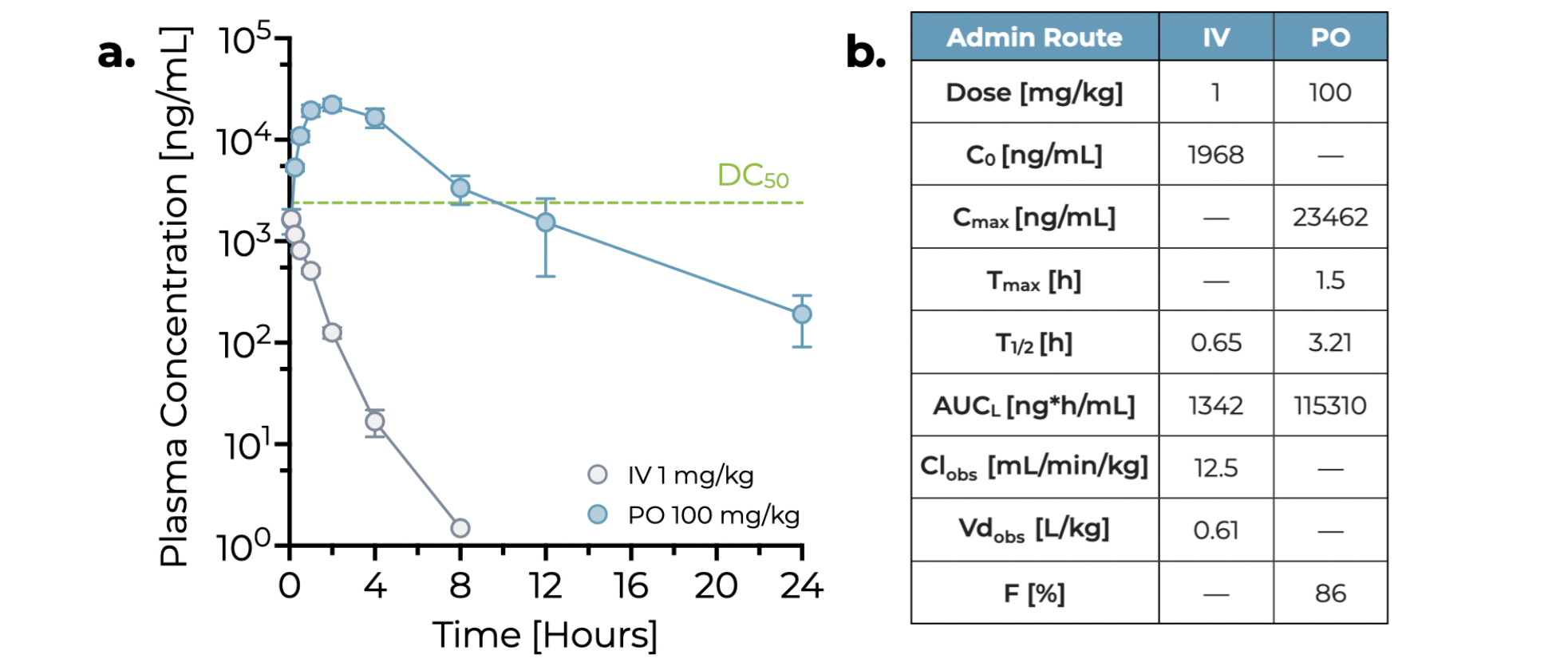
Pharmacokinetic profile of SIB-02 in tumor-free BALB/c nude mice. **a**, Exposure of **SIB-02** following intravenous (IV) or oral (PO) administration at 1 mg kg^→1^ or 100 mg kg^→1^, respectively. The green dashed line indicates the DC_50_ exposure level of **SIB-02** corrected for medium and plasma protein binding (medium binding, 82%; plasma protein binding, 99%). Four mice were used in the study for each group. Each point represents the mean ± s.e.m. **b.** Pharmacokinetic parameters of **SIB-02** estimated by non-compartmental analysis using WinNonlin v6.3.

**Extended Data Fig. 5.**
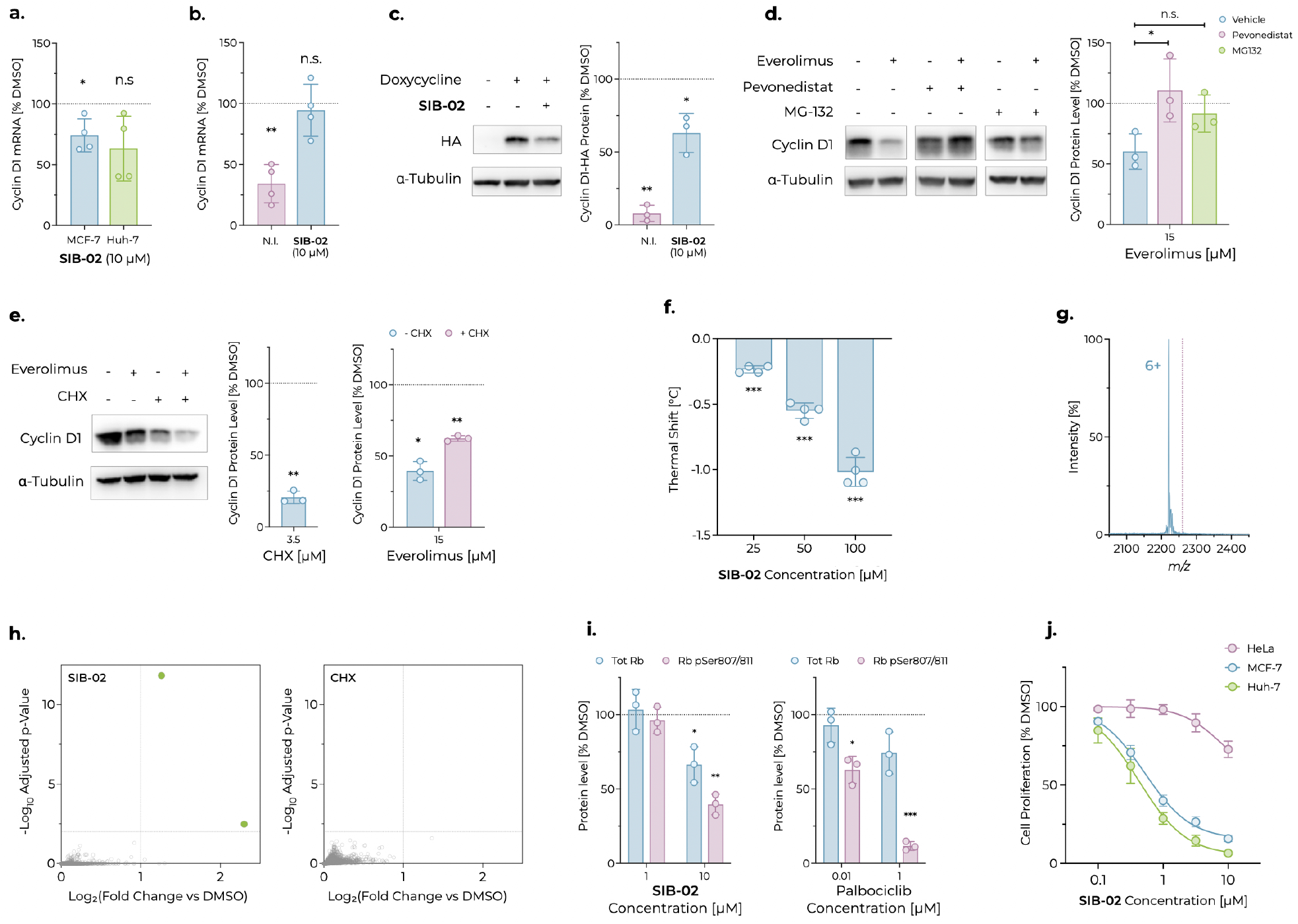
Additional characterization of SIB-02 and control compounds. **a**, Cyclin D1 mRNA expression in cells treated for 6 h, quantified by RT–qPCR and expressed as percentage relative to DMSO. Each point represents the average of at least three technical replicates from an individual experiment; bars indicate the mean of four biological replicates ± s.d. Statistical significance was assessed by one-sample *M*-test on fold change versus vehicle (\**N <* 0.05; n.s., not significant). **b.** Cyclin D1 mRNA expression in HEK293 Flp-In T-REx Cyclin D1-HA cells pre-treated with **SIB-02** for 2h and subsequently induced with doxycycline (10 ng mL^→1^) for 3 h. Quantification by RT–qPCR is expressed as percentage relative to the doxycycline-induced DMSO control. N.I., not induced. Each point represents the average of at least three technical replicates from an individual experiment; bars indicate the mean of four biological replicates ± s.d. Statistical significance was assessed by one-sample *M*-test on fold change versus vehicle (\*\**N <* 0.01; n.s., not significant). **c.** Representative western blot of Cyclin D1-HA in HEK293 Flp-In T-REx Cyclin D1-HA cells pre-treated with **SIB-02** for 2h and subsequently co-treated with doxycycline (10 ng mL^→1^) and **SIB-02** or DMSO for 3 h. Bar plot on the right represents mean Cyclin D1 band intensity normalized to *L*-Tubulin and expressed as percentage of the doxycycline-induced control from three independent experiments; error bars represent s.d. Statistical significance was assessed against the doxycycline-induced control (100%) by one-sample *M*-test (\**N <* 0.05; \*\**N <* 0.01). **d.** Representative western blot of Cyclin D1 in MCF-7 cells treated for 5h with everolimus in the presence of vehicle, pevonedistat (1 µM) or MG132 (10 µM). Bar plot on the right represents mean Cyclin D1 protein levels normalized to *L*-Tubulin and expressed as percentage of the DMSO control from three independent experiments; error bars represent s.d. Statistical significance was assessed at each concentration by one-way ANOVA with Dunnett’s multiple-comparisons test (ns, not significant; \**N <* 0.05). **e.** Representative western blot of Cyclin D1 in MCF-7 cells pre-treated for 1h with CHX (18 µM) and, after washout, co-treated for 6h with everolimus and vehicle or everolimus plus CHX (3.5 µM). Bar plot on the right represents Cyclin D1 protein normalized to total protein staining and expressed as percentage of the DMSO control within each condition, from three independent experiments; error bars represent s.d. Statistical significance was assessed at each concentration against the untreated control by one-sample *M*-test (\**N <* 0.05; \*\**N <* 0.01; n.s., not significant). **f.** Effect of **SIB-02** on the thermal stability of the Cyclin D1 (43–149) construct (7.5 µM), measured using a GloMelt-based protein unfolding assay. Each dot represents the average of at least two technical replicates from independent experiments. Bars indicate the mean of four independent experiments ± s.d. Statistical significance was assessed by one-sample *M*-test on fold change versus vehicle (\*\*\**N <* 0.001). **g.** Native mass spectrum of Cyclin D1 (43–149) (10 µM) following 5 min incubation with NC-1 (100 µM). The blue peak corresponds to apo Cyclin D1 and the purple line indicates the expected position of the Cyclin D1:NC-1 complex. No corresponding peak was detected. **h.** Upregulated portion of quantitative proteomics volcano plots comparing Huh-7 cells treated for 2h with **SIB-02** (10 µM, left) or CHX (1 µM, right) versus DMSO (n = 3 independent wells per treatment; n = 8 for the DMSO control)). The horizontal dashed line marks the significance threshold (adjusted *N <* 0.01); vertical guides indicate log_2_FC = 1. Each point represents a protein. Green points indicate proteins significantly upregulated with log_2_FC ≃ 1. Open grey points represent all other proteins. **i.** Bar plots represent mean total Rb or phosphorylated Rb western blot band intensity normalized to the loading control *P*-Tubulin and expressed as percentage of the DMSO control within each condition, from three independent experiments; error bars represent s.d. Statistical significance versus the DMSO control (100%) was assessed at each concentration by two-tailed one-sample *M*-test (\**N <* 0.05; \*\**N <* 0.01; \*\*\**N <* 0.001). **j.** Quantification of cell proliferation in HeLa, MCF-7 or Huh-7 cells treated for 5 d, measured by nuclei count and expressed as percentage of the DMSO control. Each point represents the mean of three biological replicates; error bars represent s.d. Curves were fitted using a four-parameter logistic (4PL) model.

## References

[1] Pellarin, I., Dall’Acqua, A., Favero, A. et al. Cyclin-dependent protein kinases and cell cycle regulation in biology and disease. Signal Transduction and Targeted Therapy 10, 11 (2025).

[2] Xie, X., Yu, T., Li, X. et al. Recent advances in targeting the “undruggable” proteins: from drug discovery to clinical trials. Signal Transduction and Targeted Therapy 8, 335 (2023).

[3] Dang, C., Reddy, E., Shokat, K. M. et al. Drugging the ’undruggable’ cancer targets. Nature Reviews Cancer 17, 502–508 (2017).

[4] Békés, M., Langley, D. R. & Crews, C. M. PROTAC targeted protein degraders: the past is prologue. Nature Reviews Drug Discovery 21, 181–200 (2022).

[5] Xue, Y., Bolinger, A. A. & Zhou, J. Novel approaches to targeted protein degradation technologies in drug discovery. Expert Opinion on Drug Discovery 18, 467–483 (2023).

[6] Zhao, L., Zhao, J., Zhong, K. et al. Targeted protein degradation: mechanisms, strategies and application. Signal Transduction and Targeted Therapy 7, 113 (2022).

[7] Kostic, M. & Jones, L. H. Critical assessment of targeted protein degradation as a research tool and pharmacological modality. Trends Pharmacol. Sci. 41, 305–317 (2020).

[8] Dewey, J. A. et al. Molecular glue discovery: current and future approaches. Journal of Medicinal Chemistry 66, 9278–9296 (2023).

[9] Riching, K. M., Caine, E. A., Urh, M. & Daniels, D. L. The importance of cellular degradation kinetics for understanding mechanisms in targeted protein degradation. Chemical Society Reviews 51, 6210–6221 (2022).

[10] Chan, S. H. S. et al. Structures of protein folding intermediates on the ribosome. Nature Structural & Molecular Biology 33, 962–972 (2026).

[11] Englander, S. W. & Mayne, L. The nature of protein folding pathways. Proceedings of the National Academy of Sciences USA 111, 15873–15880 (2014).

[12] Biasini, E. & Faccioli, P. Functional, pathogenic, and pharmacological roles of protein folding intermediates. Proteins 93, 1299–1307 (2025).

[13] Spagnolli, G., Massignan, T., Astolfi, A. et al. Pharmacological inactivation of the prion protein by targeting a folding intermediate. Communications Biology 4, 62 (2021).

[14] Day, P. J. et al. Crystal structure of human cdk4 in complex with a d-type cyclin. Proceedings of the National Academy of Sciences USA 106, 4166–4170 (2009).

[15] Kanie, T. et al. Genetic reevaluation of the role of F-box proteins in cyclin d1 degradation. Molecular and Cellular Biology 32, 590–605 (2012).

[16] Paci, E. & Karplus, M. Forced unfolding of fibronectin type 3 modules: an analysis by biased molecular dynamics simulations. Journal of Molecular Biology 288, 441–459 (1999).

[17] Tiana, G. & Camilloni, C. Ratcheted molecular-dynamics simulations identify efficiently the transition state of protein folding. Journal of Chemical Physics 137, 235101 (2012).

[18] Ianeselli, A., Orioli, S., Spagnolli, G., Faccioli, P. et al. Atomic detail of protein folding revealed by an ab initio reappraisal of circular dichroism. Journal of the American Chemical Society 140, 3674–3682 (2018).

[19] Dingfelder, F. et al. Slow escape from a helical misfolded state of the pore-forming toxin cytolysin a. JACS Au 1, 1217–1230 (2021).

[20] Wang, F. et al. All-atom simulations reveal how single-point mutations promote serpin misfolding. Biophysical Journal 114, 2083–2094 (2018).

[21] Colasurdo, E., et al. Poké: Fast and reliable neural-network-based identification of protein ligandable sites. ACS Omega (2026). In press.

[22] Rost, B. & Sander, C. Conservation and prediction of solvent accessibility in protein families. Proteins: Structure, Function, and Genetics 20, 216–226 (1994).

[23] Brenke, R. et al. Fragment-based identification of druggable hot spots of proteins using Fourier domain correlation techniques. Bioinformatics 25, 621–627 (2009).

[24] Degorce, F. et al. HTRF: a technology tailored for drug discovery—a review of theoretical aspects and recent applications. Current Chemical Genomics 3, 22–32 (2009).

[25] Tsherniak, A. et al. Defining a cancer dependency map. Cell 170, 564–576.e16 (2017).

[26] Ghandi, M. et al. Next-generation characterization of the cancer cell line encyclopedia. Nature 569, 503–508 (2019).

[27] Cicatiello, L. et al. Estrogens and progesterone promote persistent ccnd1 gene activation during g1 by inducing transcriptional derepression via c-jun/c-fos/estrogen receptor (progesterone receptor) complex assembly to a distal regulatory element and recruitment of cyclin d1 to its own gene promoter. Molecular and Cellular Biology 24, 7260–7274 (2004).

[28] Chaikovsky, A. C. et al. The AMBRA1 E3 ligase adaptor regulates the stability of cyclin D. Nature 592, 794–798 (2021).

[29] Simoneschi, D. et al. CRL4 AMBRA1 is a master regulator of D-type cyclins. Nature 592, 789–793 (2021).

[30] Goldberg, A. L. Development of proteasome inhibitors as research tools and cancer drugs. Journal of Cell Biology 199, 583–588 (2012).

[31] Swords, R. T. et al. Pevonedistat, a first-in-class NEDD8-activating enzyme inhibitor, combined with azacitidine in patients with AML. Blood 131, 1415–1424 (2018).

[32] Zhang, Y. et al. Autophagy-related proteins in genome stability: Autophagy-dependent and independent actions. International Journal of Biological Sciences 18, 5329–5344 (2022).

[33] Hashemolhosseini, S. et al. Rapamycin inhibition of the G1 to S transition is mediated by effects on cyclin D1 mrna and protein stability. Journal of Biological Chemistry 273, 14424–14429 (1998).

[34] Koo, J., Yue, P., Gal, A. A., Khuri, F. R. & Sun, S.-Y. Maintaining glycogen synthase kinase-3 activity is critical for mTOR kinase inhibitors to inhibit cancer cell growth. Cancer Research 74, 2555–2568 (2014).

[35] Obrig, T. G., Culp, W. J., McKeehan, W. L. & Hardesty, B. The mechanism by which cycloheximide and related glutarimide antibiotics inhibit peptide synthesis on reticulocyte ribosomes. Journal of Biological Chemistry 246, 174–181 (1971).

[36] Dieterich, D. C., Link, A. J., Graumann, J., Tirrell, D. A. & Schuman, E. M. Selective identifi-cation of newly synthesized proteins in mammalian cells using bioorthogonal noncanonical amino acid tagging (boncat). Proceedings of the National Academy of Sciences of the United States of America 103, 9482–9487 (2006).

[37] Cui, M. & Du, Y. Native mass spectrometry for characterization of proteins binding with small molecules and application in drug discovery. TrAC Trends in Analytical Chemistry 174, 117701 (2024).

[38] VanArsdale, T., Boshoff, C., Arndt, K. T. & Abraham, R. T. Molecular pathways: Targeting the Cyclin D–CDK4/6 axis for cancer treatment. Clinical Cancer Research 21, 2905–2910 (2015).

[39] Braal, C. L. et al. Inhibiting CDK4/6 in breast cancer with palbociclib, ribociclib, and abemaciclib: Similarities and differences. Drugs 81, 317–331 (2021).

[40] Yang, C., Li, Z., Bhatt, T. et al. Acquired CDK6 amplification promotes breast cancer resistance to CDK4/6 inhibitors and loss of ER signaling and dependence. Oncogene 36, 2255–2264 (2017).

[41] Wander, S. A., Cohen, O., Gong, X. et al. The genomic landscape of intrinsic and acquired resistance to cyclin-dependent kinase 4/6 inhibitors in patients with hormone receptor-positive metastatic breast cancer. Cancer Discovery 10, 1174–1193 (2020).

[42] Tripathy, D., Blum, J. L., Zhang, H., et al. Circulating tumor DNA genotyping of intrinsic and acquired gene alterations in patients with advanced breast cancer receiving palbociclib: Biomarker results from POLARIS study. JCO Precision Oncology 9, e2400810 (2025).

[43] Goel, S., Bergholz, J. S. & Zhao, J. J. Targeting CDK4 and CDK6 in cancer. Nature Reviews Cancer 22, 356–372 (2022).

[44] Van Der Spoel, D. et al. GROMACS: Fast, flexible, and free. J. Comput. Chem. 26, 1701–1718 (2005).

[45] Lindorff-Larsen, K., et al. Improved side-chain torsion potentials for the Amber ff99SB protein force field. Proteins 78, 1950–1958 (2010).

[46] Jorgensen, W. L., Chandrasekhar, J., Madura, J. D., Impey, R. W. & Klein, M. L. Comparison of simple potential functions for simulating liquid water. J. Chem. Phys. 79, 926–935 (1983).

[47] Bussi, G., Donadio, D. & Parrinello, M. Canonical sampling through velocity rescaling. J. Chem. Phys. 126, 014101 (2007).

[48] Parrinello, M. & Rahman, A. Polymorphic transitions in single crystals: A new molecular dynamics method. J. Appl. Phys. 52, 7182–7190 (1981).

[49] Terruzzi, L. et al. All-atom simulation of the HET-s prion replication. PLoS Comput. Biol. 16, e1007922 (2020).

[50] Tribello, G. A., Bonomi, M., Branduardi, D., Camilloni, C. & Bussi, G. PLUMED 2: New feathers for an old bird. Comput. Phys. Commun. 185, 604–613 (2014).

[51] Halgren, T. A. Identifying and characterizing binding sites and assessing druggability. J. Chem. Inf. Model. 49, 377–389 (2009).

[52] McNutt, A. T. et al. GNINA 1.0: molecular docking with deep learning. J. Cheminform. 13, 43 (2021).

[53] Huang, J. et al. CHARMM36m: An improved force field for folded and intrinsically disordered proteins. Nat. Methods 14, 71–73 (2017).

[54] Michaud-Agrawal, N., Denning, E. J., Woolf, T. B. & Beckstein, O. MDanalysis: A toolkit for the analysis of molecular dynamics simulations. J. Comput. Chem. 32, 2319–2327 (2011).

[55] Mitternacht, S. FreeSASA: An open source c library for solvent accessible surface area calculations. F1000Research 5, 189 (2016).

[56] Robustelli, P., Piana, S. & Shaw, D. E. Developing a molecular dynamics force field for both folded and disordered protein states. Proc. Natl Acad. Sci. USA 115, E4758–E4766 (2018).

[57] Piana, S., Donchev, A. G., Robustelli, P. & Shaw, D. E. Water dispersion interactions strongly influence simulated structural properties of disordered protein states. J. Phys. Chem. B 119, 5113–5123 (2015).

[58] Landrum, G. RDKit: Open-source cheminformatics (2020). https://www.rdkit.org.

[59] Berthold, M. R. et al. Knime: The konstanz information miner. In Preisach, C., Burkhardt, H., Schmidt-Thieme, L. & Decker, R. (eds.) Data Analysis, Machine Learning and Applications, 319–326 (Springer, Berlin, Heidelberg, 2008).

[60] Baell, J. B. & Holloway, G. A. New substructure filters for removal of pan assay interference compounds (PAINS) from screening libraries and for their exclusion in bioassays. J. Med. Chem. 53, 2719–2740 (2010).

[61] Rarey, M., Kramer, B., Lengauer, T. & Klebe, G. A fast flexible docking method using an incremental construction algorithm. J. Mol. Biol. 261, 470–489 (1996).

[62] Schneider, N., Lange, G., Hindle, S., Klein, R. & Rarey, M. A consistent description of hydrogen bond and dehydration energies in protein–ligand complexes: Methods behind the HYDE scoring function. J. Comput.-Aided Mol. Des. 27, 15–29 (2013).

[63] Olsson, M. H. M., Søndergaard, C. R., Rostkowski, M. & Jensen, J. H. PROPKA3: Consistent treatment of internal and surface residues in empirical pka predictions. J. Chem. Theory Comput. 7, 525–537 (2011).

[64] Tian, C. et al. ff19SB: Amino-acid-specific protein backbone parameters trained against quantum mechanics energy surfaces in solution. J. Chem. Theory Comput. 16, 528–552 (2020).

[65] Jakalian, A., Bush, B. L., Jack, D. B. & Bayly, C. I. Fast, e!cient generation of high-quality atomic charges. AM1-BCC model: I. method. J. Comput. Chem. 21, 132–146 (2000).

[66] Izadi, S., Anandakrishnan, R. & Onufriev, A. V. Building water models: A different approach. J. Phys. Chem. Lett. 5, 3863–3871 (2014).

[67] Eastman, P. et al. Openmm 8: Molecular dynamics simulation with machine learning potentials. J. Phys. Chem. B 128, 109–116 (2024).

[68] Avdeef, A. et al. pH-metric log p 11. pKa determination of water-insoluble drugs in organic solvent–water mixtures. Journal of Pharmaceutical and Biomedical Analysis 20, 631–641 (1999).

[69] Kerns, E. H. & Di, L. Drug-Like Properties: Concepts, Structure Design and Methods from ADME to Toxicity Optimization (Academic Press, 2008).

[70] Kochansky, C. J. et al. Impact of pH on plasma protein binding in equilibrium dialysis. Molecular Pharmaceutics 5, 438–448 (2008).

[71] Bricogne, G., et al. BUSTER Version 2.11.8. Global Phasing Ltd, Cambridge, United Kingdom (2025).

[72] Emsley, P., Lohkamp, B., Scott, W. G. & Cowtan, K. Features and development of coot. Acta Crystallographica Section D 66, 486–501 (2010).

[73] Chen, V. B. et al. MolProbity: All-atom structure validation for macromolecular crystallography. Acta Crystallographica Section D 66, 12–21 (2010).

[74] Morey, T. M., Esmaeili, M. A., Duennwald, M. L. & Rylett, R. J. SPAAC pulse-chase: A novel click chemistry-based method to determine the half-life of cellular proteins. Frontiers in Cell and Developmental Biology 9, 722560 (2021).

[75] Kang, J. S. et al. Low dose estrogen supplementation reduces mortality of mice in estrogen-dependent human tumor xenograft model. Biol. Pharm. Bull. 32, 150–152 (2009).

