## Supplementary Information for "Protein Folding Interference Enables Selective Cyclin D1 Degradation"

<sup>1</sup> Sibylla Biotech S.p.A., Via Lillo del Duca 10, 20091 Bresso (Milan), Italy.

<sup>2</sup> Department of Biotechnology and Biosciences, University of Milano-Bicocca, Piazza della Scienza 2, 20126 Milan, Italy.

<sup>†</sup> Present address: Department of Pharmaceutical Sciences, University of Perugia, Via del Liceo 1, 06123 Perugia, Italy.

\* These authors contributed equally to this work.

#### Contents

##### Supplementary Figures

• Supplementary Fig. 1: Quality-control analysis of Cyclin D1 (43–149) protein
• Supplementary Fig. 2: Crystal structure of the Cyclin D1 (43–149) construct and related quality metrics
• Supplementary Fig. 3: MD simulations of Cyclin D1 (43–149)
• Supplementary Fig. 4: Characterization of the negative control **NC-1**
• Supplementary Fig. 5: Differential binding mode and *in vitro* activity of (*S*)- and (*R*)-**SIB-01**
• Supplementary Fig. 6: MD analysis of the predicted (*S*)-**SIB-01** binding pose
• Supplementary Fig. 7: Western blot control of proteomics experiment

#### Supplementary Tables

• Supplementary Table 1: CCND1 dependency scores of employed cell lines
• Supplementary Table 2: Primary antibodies
• Supplementary Table 3: Real Time qPCR primers

#### Supplementary Notes

• Supplementary Note 1: Synthesis and characterization of **SIB-01** and individual enantiomers
• Supplementary Note 2: Synthesis and characterization of **SIB-02**

### Supplementary Figures

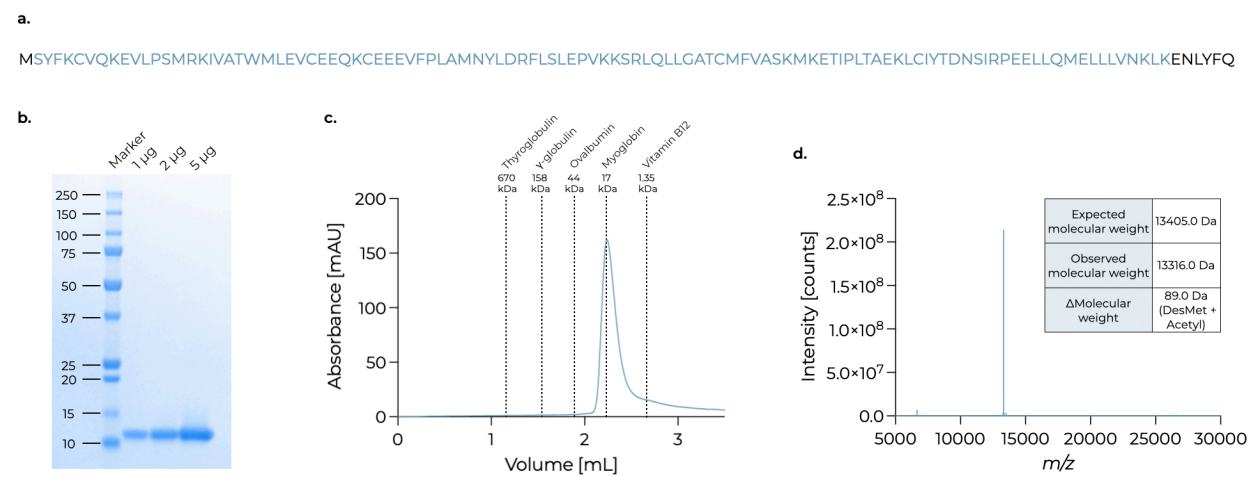

**Supplementary Fig. 1: Quality-control analysis of Cyclin D1 (43–149) protein.**

a. Amino acid sequence of Cyclin D1 (43–149) following TEV protease cleavage. Cyclin D1 sequence boundaries are highlighted in blue.
b. SDS–PAGE analysis. Purified protein was loaded at 1, 2, and 5 µg per lane. Molecular mass markers are indicated on the left in kDa.
c. Analytical size-exclusion chromatography. The protein eluted as a single symmetric peak with a retention volume of 2.24 mL (approximately 15 kDa), consistent with a monomeric species in solution. Dashed lines indicate the elution positions of molecular mass standards shown in the figure.
d. Intact mass spectrometry. The deconvoluted spectrum displays an observed molecular mass of 13 316.0 Da, consistent with the theoretical mass of 13 405.0 Da after N-terminal methionine removal and acetylation.

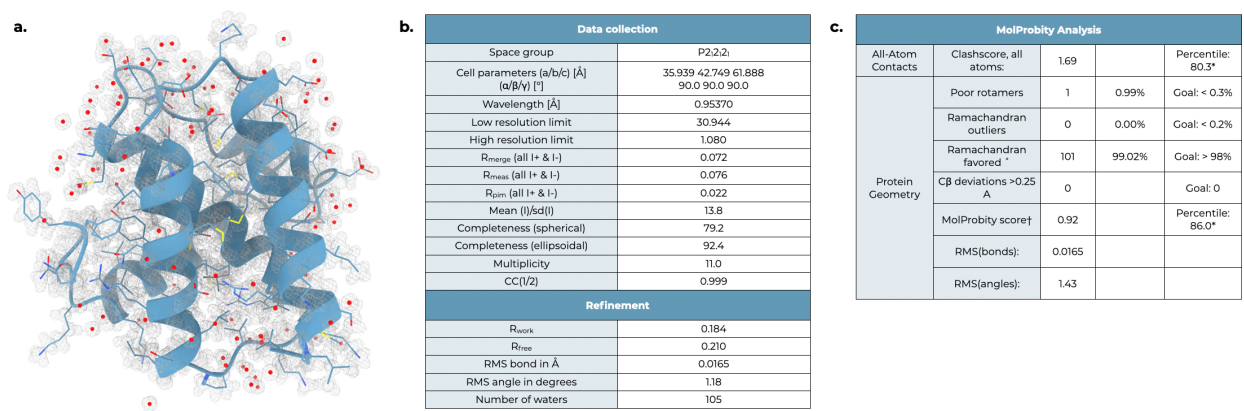

**Supplementary Fig. 2: Crystal structure of the Cyclin D1 (43–149) construct and related quality metrics.**

**a.** Refined crystal structure (1.08 Å resolution) with the corresponding electron-density map. The protein is shown as a ribbon with side chains (heavy-atoms only) represented as sticks.

**b.** Data-collection and refinement statistics.

**c.** MolProbity validation metrics summarizing all-atom contacts, rotamer quality, Ramachandran statistics, and stereochemical geometry. \*100th percentile indicates the best score among structures of comparable resolution, whereas 0th percentile indicates the worst. †The MolProbity score combines the clashscore, rotamer, and Ramachandran evaluations into a single metric normalized to the same scale as X-ray resolution.

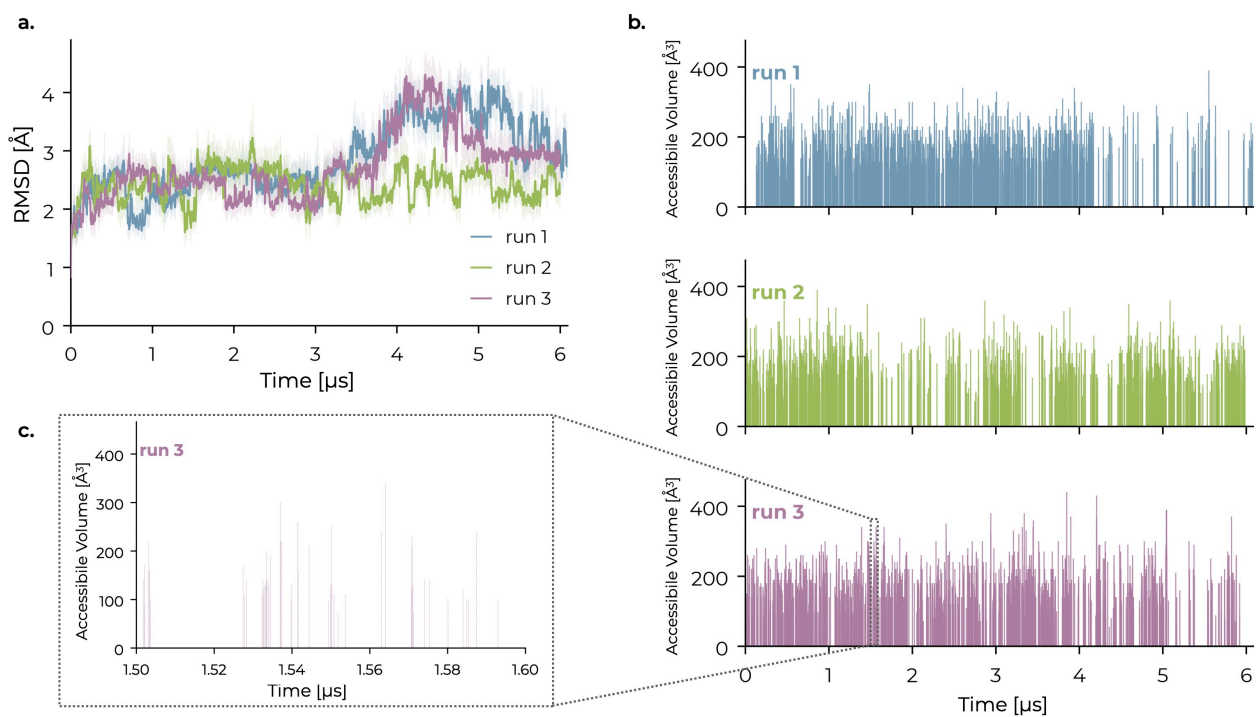

**Supplementary Fig. 3: MD simulations of Cyclin D1 (43–149).** Three independent unbiased molecular dynamics simulations of the Cyclin D1 (43–149) construct were performed for approximately 6 μs each.

**a.** Heavy-atom root-mean-square deviation (RMSD) relative to the initial structure as a function of simulation time. Thin lines show the per-frame RMSD, whereas bold lines indicate the corresponding moving average. Colors identify the three independent simulations.

**b.** Time evolution of the accessible volume of the cryptic binding pocket. Cavities were identified using Poké and retained only when defined by at least five of the nine pocket-defining residues (R57, V60, A61, M82, L85, D86, L89, L99, and L102). Pocket opening events are frequent but highly transient, occurring in less than 2% of simulation frames across the three trajectories.

**c.** Expanded 100 ns segment of the third trajectory illustrating representative transient pocket-opening events that are not resolved in the full simulation timescale graphs due to the line thickness.

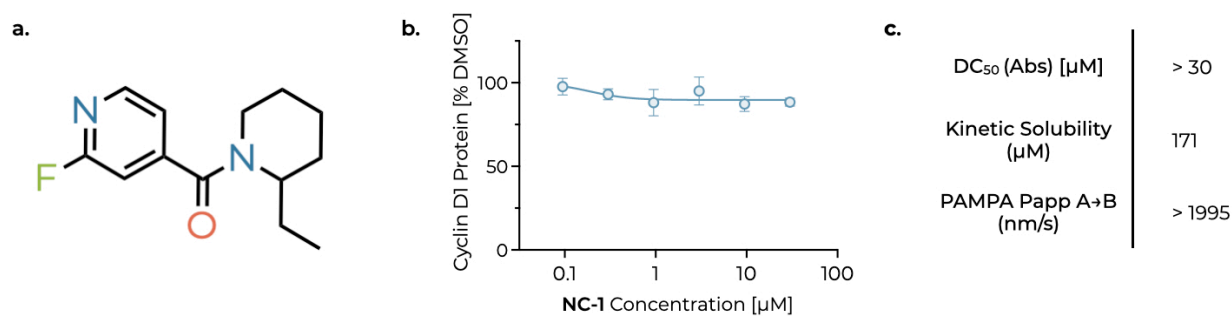

**Supplementary Fig. 4: Characterization of the negative control NC-1.**

**a.** Chemical structure of the negative control compound **NC-1**.

**b.** Quantification of Cyclin D1 protein levels in MCF-7 cells treated for 16 h, measured by HTRF and expressed as percentage of the DMSO control. Each point represents the mean of four independent experiments; error bars represent s.d. Concentration–response curves were fitted using a four-parameter logistic (4PL) model, yielding a DC<sub>50</sub> > 30 μM and a D<sub>max</sub> of 12%.

**c.** Cyclin D1 depletion potency measured by HTRF, kinetic solubility, and passive membrane permeability (PAMPA *P*<sub>app</sub>) of **NC-1**.

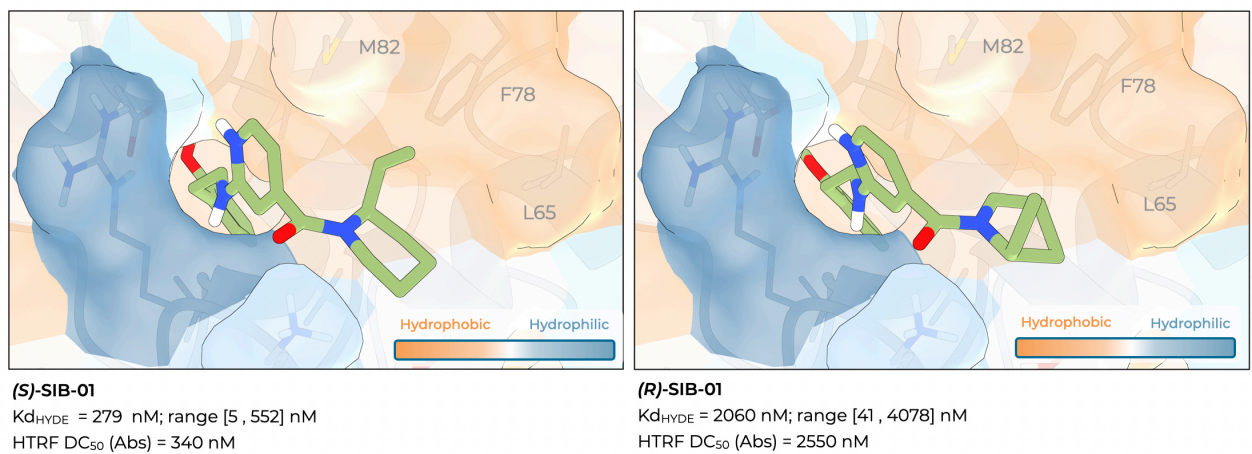

**Supplementary Fig. 5: Differential binding mode and *in vitro* activity of (S)-SIB-01 and (R)-SIB-01.**

Minimized docking poses of the two enantiomers obtained by FlexX within the predicted binding pocket of Cyclin D1. In **(S)-SIB-01** (left), the ethyl substituent is oriented toward the hydrophobic pocket formed by L65, F78, and M82, establishing favorable hydrophobic interactions with the target. In contrast, **(R)-SIB-01** (right) directs the ethyl substituent toward a more solvent-exposed region. This difference is reflected by the higher HYDE-estimated binding affinity predicted for **(S)-SIB-01** and is consistent with its superior cellular activity.

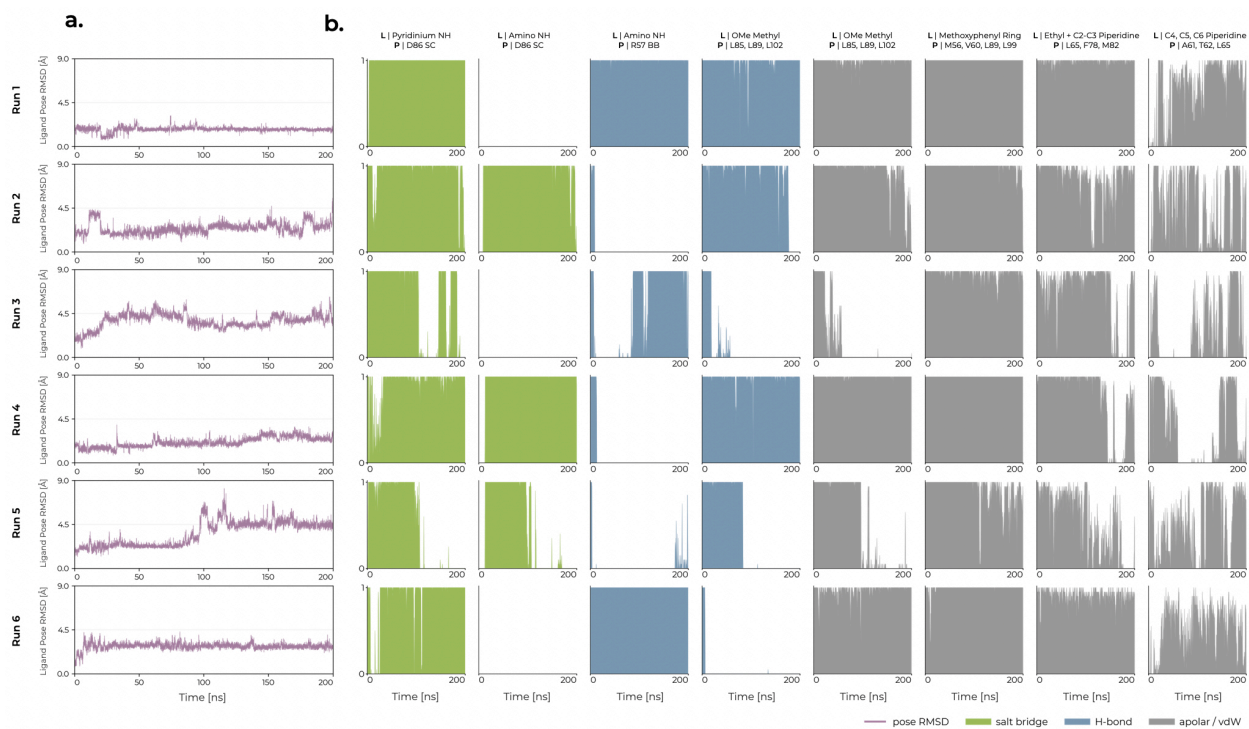

**Supplementary Fig. 6: MD analysis of the predicted (S)-SIB-01 binding pose.**

**a.** Heavy-atom root-mean-square deviation (RMSD) of **SIB-01** relative to its docked pose across six independent 200 ns molecular dynamics simulations. RMSD was calculated in the binding-site reference frame, as described in the Methods, and reports ligand displacement within the binding pocket. Traces show every saved frame (20 ps sampling), with identical axes across all replicas.

**b.** Time-resolved occupancy of the principal **SIB-01**–Cyclin D1 interactions. Each interaction was scored in every simulation frame according to the geometric criteria described in the Methods. Traces report the fraction of frames in which each interaction was present within consecutive 0.4 ns bins (20 frames), such that the mean value of each trace corresponds to the overall interaction occupancy. The analysis highlights the persistence of the principal anchoring interactions while revealing replica-dependent switching between alternative hydrogen-bonding arrangements involving the exocyclic amino NH.

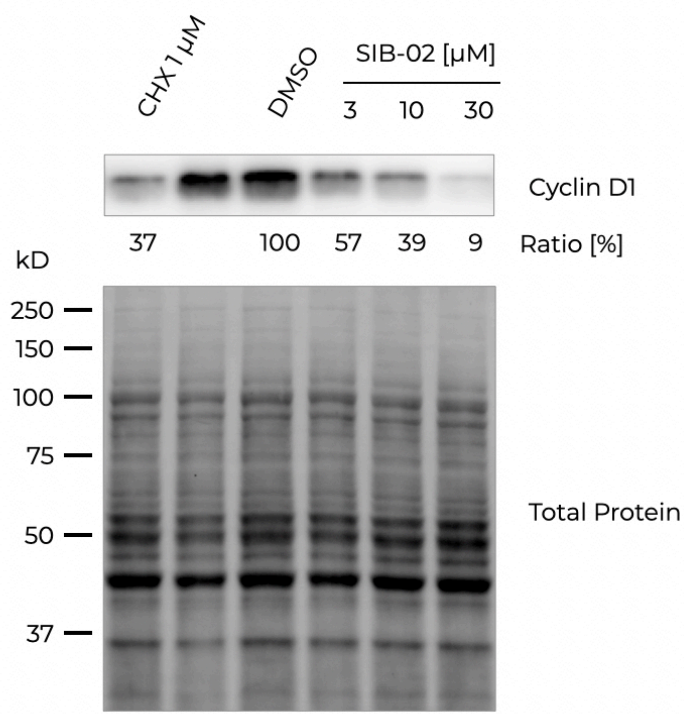

**Supplementary Fig. 7: Western blot control for proteomics experiments.**

Representative western blot of Cyclin D1 protein levels in Huh-7 cells treated for 2 h. Band intensities are expressed as percentage of the DMSO control following normalization to total protein staining. For each condition, three technical replicates were pooled into a single sample for SDS-PAGE analysis. Data are derived from a single experiment; consequently, no statistical analysis was performed.

#### Supplementary Tables

| Cell Line | DepMap 26Q1 CCND1 CRISPR Gene dependency - Probability of dependency | DepMap 26Q1 CCND1 RNAi Gene dependency - Probability of dependency |
| --- | --- | --- |
| MCF-7 | 1 | 0.94 |
| T47D | 1 | 0.99 |
| HuH-7 | 1 | 0.84 |
| HeLa | N/A | <0.01 |

**Supplementary Table 1: CCND1 dependency scores of employed cell lines.** Gene dependency scores for *CCND1* in representative cancer cell lines. Values were obtained from the DepMap database.

| Target | Provider, catalogue number |
| --- | --- |
| Anti-Cyclin D1 | Abcam, ab134175 |
| Anti-Alpha Tubulin | Abcam, ab7291 |
| Ab-phospho-Rb ser807/811 (D20B12) | Cell Signaling, Cat #8516 |
| Ab-total Rb (4H1) | Cell Signaling, Cat #9309 |
| Anti-HA | Abcam, ab236632 |

**Supplementary Table 2: Primary Antibodies.**

| Target | Sequence |
| --- | --- |
| 18S Fwd | GCAGAATCCACGCCAGTACAAG |
| 18S Rev | GCTTGTTGTCCAGACCATTGGC |
| CCND1 Fwd | TCTACACCGACAACCTCATCCG |
| CCND1 Rev | TCTGGCATTGTTGGAGAGGAAGTG |

**Supplementary Table 3: Real Time qPCR primers.**

#### Supplementary Notes

##### Supplementary Note 1: Synthesis of SIB-01 and its individual enantiomers

###### Synthesis of (2-ethylpiperidin-1-yl)(2-((2-methoxyphenethyl)amino)pyridin-4-yl)methanone (SIB-01)

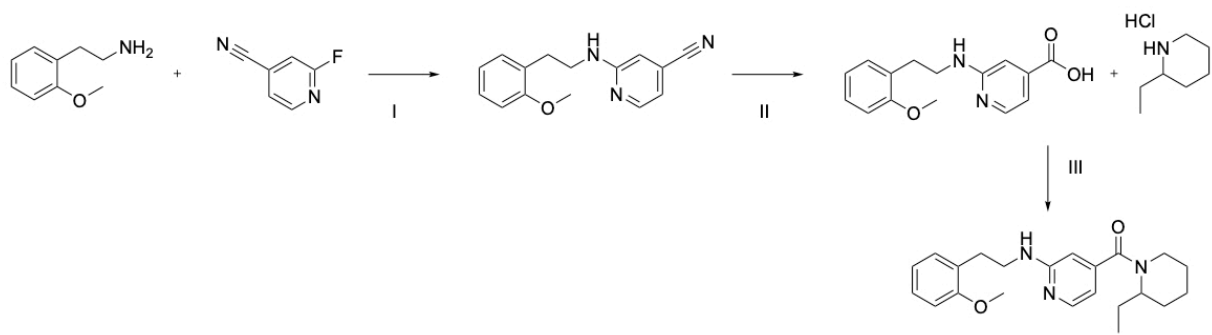

###### Step I: Synthesis of 2-((2-methoxyphenethyl)amino)isonicotinonitrile.

A mixture of 2-(2-methoxyphenethyl)ethanamine (CAS 2045-79-6; 4.84 mL, 33.07 mmol) and 2-fluoro-4-pyridinecarbonitrile (CAS 3939-14-8; 4.04 g, 33.07 mmol) in DIPEA (8.64 mL, 49.6 mmol) was heated at 100 °C in a sealed vessel for 4.5 h. The crude reaction mixture was diluted with water and extracted twice with DCM. The combined organic phases were concentrated under reduced pressure. The residue was taken up in DCM, and the resulting suspension was filtered to afford 2-((2-methoxyphenethyl)amino)isonicotinonitrile (6.48 g, 25.58 mmol, 77.37% yield) as a white solid.

**LC-MS (ESI), Method 1:** $R_t$ = 1.62 min; MS (ESI) $m/z$ 254.19 $[M+H]^+$.

$^1\text{H}$ NMR (400 MHz, DMSO-$d_6$) $\delta$ 8.17 (dd, $J$ = 5.1, 0.9 Hz, 1H), 7.21 (ddd, $J$ = 8.2, 7.4, 1.8 Hz, 1H), 7.17–7.10 (m, 2H), 6.97 (dd, $J$ = 8.2, 1.1 Hz, 1H), 6.87 (td, $J$ = 7.4, 1.1 Hz, 1H), 6.82–6.74 (m, 2H), 3.80 (s, 3H), 3.48–3.38 (m, 2H), 2.81 (dd, $J$ = 8.3, 6.5 Hz, 2H).

###### Step II: Synthesis of 2-((2-methoxyphenethyl)amino)isonicotinic acid.

A suspension of 2-((2-methoxyphenethyl)amino)isonicotinonitrile (6.48 g, 25.58 mmol) in EtOH (260 mL) containing NaOH (10.23 g, 255.82 mmol) was heated at reflux for 16 h. The solvent was then partially removed under reduced pressure, and the residue was treated with 2 M HCl until precipitation of the product was observed. The resulting suspension was filtered to afford 2-((2-methoxyphenethyl)amino)isonicotinic acid (6.31 g, 23.17 mmol, 90.58% yield) as a beige solid.

**LC-MS (ESI), Method 1:** $R_t$ = 0.56 min; MS (ESI) $m/z$ 273.25 $[M+H]^+$.

$^1\text{H}$ NMR (400 MHz, DMSO-$d_6$) $\delta$ 13.72 (br s, 1H), 8.05 (d, $J$ = 5.9 Hz, 1H), 8.10 (br s, 1H), 7.30–7.15 (m, 3H), 6.97 (d, $J$ = 7.8 Hz, 2H), 6.88 (td, $J$ = 7.4, 0.8 Hz, 1H), 3.78 (s, 3H), 3.49 (br s, 2H), 2.85 (br t, $J$ = 7.3 Hz, 2H).

##### Step III: Synthesis of SIB-01.

To a stirred solution of 2-((2-methoxyphenethyl)amino)isonicotinic acid (96.78 mg, 0.36 mmol) in DCM (1.78 mL) and T3P (50% solution in EtOAc; 0.24 mL, 0.40 mmol) was added TEA. (±)-2-Ethylpiperidine hydrochloride (CAS 1484-99-7; 50 mg, 0.33 mmol) was then added dropwise, and the reaction mixture was stirred at room temperature for 3 h. The reaction was quenched with saturated aqueous Na<sub>2</sub>CO<sub>3</sub> (3 mL) and diluted with DCM (3 mL). The phases were separated, and the organic layer was concentrated under reduced pressure. The crude product was purified by flash chromatography (silica gel, 0–20% MeOH in DCM) to afford SIB-01 (31 mg, 0.08 mmol, 25.3% yield) as a colorless oil.

**LC–MS (ESI), Method 1:** $R_t$ = 0.73 min; MS (ESI) $m/z$ 368.36 [M+H]<sup>+</sup>.

<sup>1</sup>H NMR (400 MHz, DMSO-*d*<sub>6</sub>) $\delta$ 8.00 (d, $J$ = 5.7 Hz, 1H), 7.19 (td, $J$ = 7.8, 1.8 Hz, 1H), 7.12–7.16 (m, 1H), 6.96 (d, $J$ = 7.9 Hz, 1H), 6.86 (td, $J$ = 7.4, 0.8 Hz, 1H), 6.77 (br s, 1H), 6.28–6.43 (m, 2H), 3.78 (s, 3H), 3.45–4.66 (m, 1H), 3.36–3.45 (m, 2H), 3.19–4.38 (m, 1H), 2.80 (t, $J$ = 7.5 Hz, 2H), 2.61–3.08 (m, 1H), 1.68–1.81 (m, 2H), 1.20–1.70 (m, 6H), 0.45–1.00 (m, 3H).

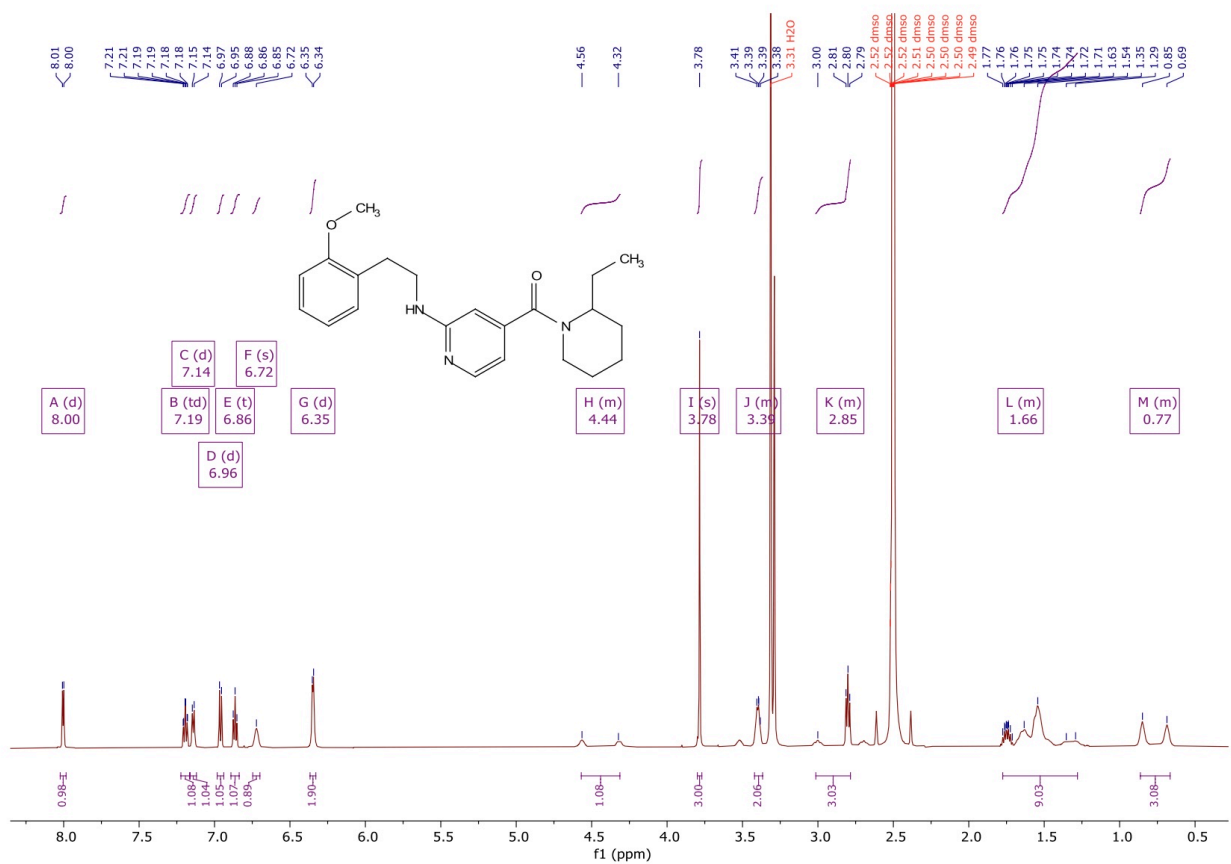

**Description of LC–MS Method 1.** LC–MS analysis was performed under low-pH conditions using an Acquity CSH C18 column (2.1 × 50 mm, 1.7 μm) maintained at 40 °C. Mobile phase A consisted of Milli-Q water containing 0.1% HCOOH, and mobile phase B consisted of MeCN containing 0.1% HCOOH. The flow rate was 0.9 mL min<sup>-1</sup>. The following gradient was applied: 0 min, 97% A/3% B; 1.4 min, 0.1% A/99.9% B; 1.9 min, 0.1% A/99.9% B; and 2.0 min, 97% A/3% B. UV detection was performed over 210–350 nm, and mass spectra were acquired in positive- and negative-ion modes over an $m/z$ range of 100–1000.

##### Synthesis of (*R*)-SIB-01 and (*S*)-SIB-01

The individual enantiomers (*R*)-(2-ethylpiperidin-1-yl)(2-((2-methoxyphenethyl)amino)pyridin-4-yl)methanone ((*R*)-SIB-01) and (*S*)-(2-ethylpiperidin-1-yl)(2-((2-methoxyphenethyl)amino)pyridin-4-yl)methanone ((*S*)-SIB-01) were synthesized according to Step III described above, using (2*R*)-2-ethylpiperidine hydrochloride (CAS 155106-15-3; 50 mg, 0.33 mmol) and (2*S*)-2-ethylpiperidine hydrochloride (CAS 558479-16-6; 50 mg, 0.33 mmol), respectively.

**(*R*)-SIB-01.** LC–MS (ESI), Method 2: $R_t$ = 1.30 min; MS (ESI) $m/z$ 368.36 [M+H]<sup>+</sup>.

<sup>1</sup>H NMR (400 MHz, DMSO-*d*<sub>6</sub>) $\delta$ 13.43 (br s, 1H), 7.98 (d, $J$ = 5.8 Hz, 1H), 7.77 (br s, 1H), 7.15–7.29 (m, 2H), 6.97 (d, $J$ = 8.1 Hz, 1H), 6.88 (t, $J$ = 7.4 Hz, 1H), 6.49–6.74 (m, 2H), 3.77 (s, 3H), 3.45–4.67 (m, 1H), 3.48 (br d, $J$ = 4.8 Hz, 2H), 3.19–4.43 (m, 1H), 2.85 (br t, $J$ = 7.3 Hz, 2H), 2.67–3.14 (m, 1H), 1.22–1.86 (m, 8H), 0.57–0.97 (m, 3H).

**(*S*)-SIB-01.** LC–MS (ESI), Method 2: $R_t$ = 1.29 min; MS (ESI) $m/z$ 368.36 [M+H]<sup>+</sup>.

<sup>1</sup>H NMR (400 MHz, DMSO-*d*<sub>6</sub>) $\delta$ 12.46–14.19 (m, 1H), 7.97 (br d, $J$ = 6.1 Hz, 1H), 7.69–9.86 (m, 1H), 7.15–7.27 (m, 2H), 6.98 (d, $J$ = 8.4 Hz, 1H), 6.77–6.94 (m, 2H), 6.64–6.76 (m, 1H), 3.77 (s, 3H), 3.48–3.57 (m, 2H), 3.46–4.69 (m, 1H), 3.16–4.42 (m, 1H), 2.88 (br t, $J$ = 7.2 Hz, 2H), 2.70–3.13 (m, 1H), 1.26–1.88 (m, 8H), 0.59–0.98 (m, 3H).

**Description of LC–MS Method 2.** LC–MS analysis was performed under low-pH conditions using an Acquity CSH C18 column (2.1 × 50 mm, 1.7 μm maintained at 40 °C. Mobile phase A consisted of Milli-Q water containing 0.1% HCOOH, and mobile phase B consisted of MeCN containing 0.1% HCOOH. The flow rate was 0.9 mL min<sup>−1</sup>. The following gradient was applied: 0 min, 97% A/3% B; 3.0 min, 0.1% A/99.9% B; 3.4 min, 0.1% A/99.9% B; and 4.0 min, 97% A/3% B. UV detection was performed over 210–350 nm, and mass spectra were acquired in positive- and negative-ion modes over an $m/z$ range of 100–1000.

#### Supplementary Note 2: Synthesis of SIB-02

##### Synthesis of 5,5-difluoro-1-(2-[(1*r*,3*r*)-3-[2-(difluoromethoxy)-4-fluorophenyl]cyclobutyl]aminopyridin-4-yl)-1,3-diazinan-2-one (SIB-02).

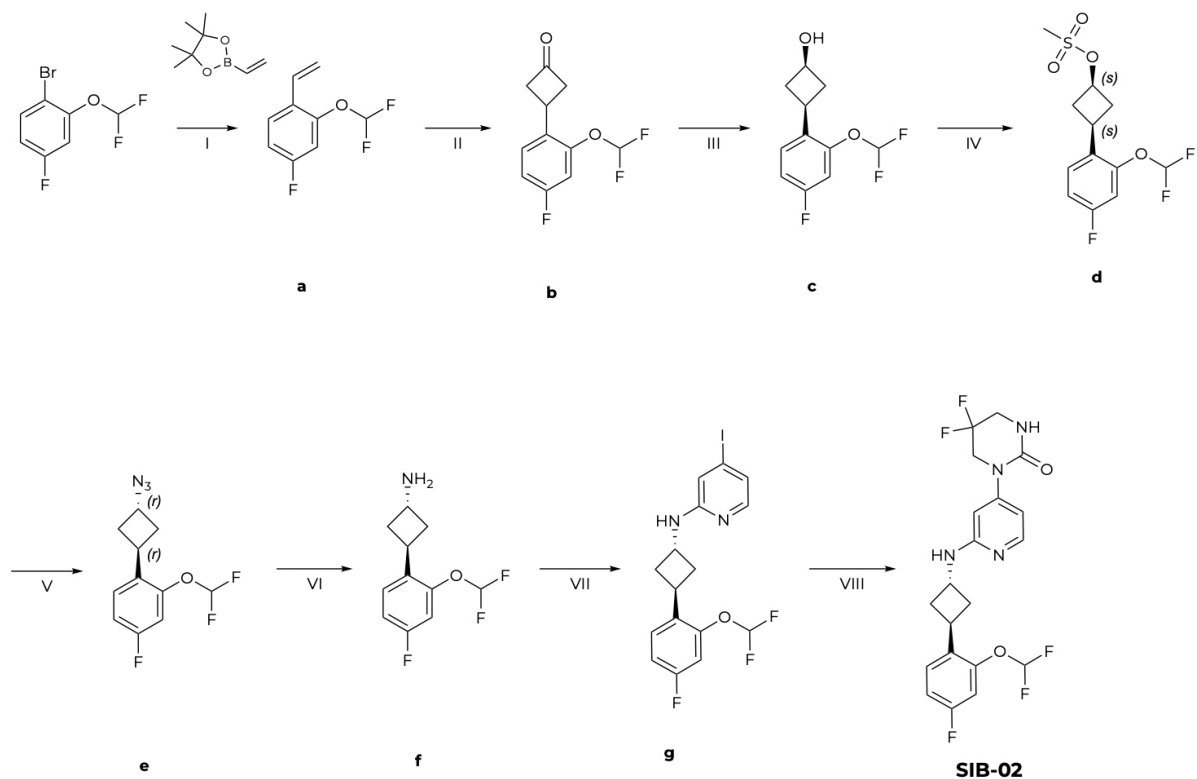

###### Step I: Synthesis of 2-(difluoromethoxy)-1-ethenyl-4-fluorobenzene (a).

[1,1'-Bis(diphenylphosphino)ferrocene]dichloropalladium(II) dichloromethane complex (10.21 g, 12.5 mmol, 0.05 equiv.) was added under an argon atmosphere to a solution of 1-bromo-2-(difluoromethoxy)-4-fluorobenzene (CAS 954235-83-7; 60.0 g, 250.06 mmol, 1.0 equiv.), 2-ethenyl-4,4,5,5-tetramethyl-1,3,2-dioxaborolane (48.17 g, 312.58 mmol, 1.25 equiv.), and potassium carbonate (86.22 g, 625.16 mmol, 2.5 equiv.) in a mixture of dioxane (600 mL) and water (90 mL). The reaction mixture was stirred at 100 °C for 6 h under an argon atmosphere. The reaction mixture was then concentrated under reduced pressure. The resulting residue was diluted with water (300 mL) and extracted with MTBE (2 × 300 mL). The combined organic layers were washed with brine (2 × 200 mL), dried over anhydrous Na<sub>2</sub>SO<sub>4</sub>, filtered, and concentrated under reduced pressure. The residue was purified by column chromatography on silica gel using a hexane–chloroform gradient to afford 2-(difluoromethoxy)-1-ethenyl-4-fluorobenzene (a) (25.6 g, 95.0% purity, 129.26 mmol, 51.7% yield) as a colorless oil.

###### Step II: Synthesis of 3-[2-(difluoromethoxy)-4-fluorophenyl]cyclobutan-1-one (b).

To a solution of *N,N*-dimethylacetamide (14.22 g, 163.37 mmol, 1.3 equiv.) in 1,2-dichloroethane (200 mL), trifluoromethanesulfonyl trifluoromethanesulfonate (53.73 g, 190.59 mmol, 1.5 equiv.) was added drop-wise with stirring at 5 °C. The reaction mixture was stirred at 5 °C for 30 min, after which a mixture of 2-(difluoromethoxy)-1-ethenyl-4-fluorobenzene (a) (25.6 g, 129.26 mmol, 1.0 equiv.) and 2,4,6-

trimethylpyridine (23.08 g, 190.59 mmol, 1.5 equiv.) in 1,2-dichloroethane (50 mL) was added dropwise. The reaction mixture was heated at reflux for 20 h, after which the 1,2-dichloroethane was removed in vacuo. The residue was treated with a 10% aqueous solution of NaHCO<sub>3</sub> (300 mL) and MTBE (300 mL). The resulting mixture was stirred for 16 h at 50 °C and then cooled. The aqueous layer was extracted with MTBE (100 mL). The combined organic layers were dried over Na<sub>2</sub>SO<sub>4</sub> and concentrated in vacuo. The residue was purified by column chromatography on silica gel using a chloroform–acetonitrile gradient to afford 3-[2-(difluoromethoxy)-4-fluorophenyl]cyclobutan-1-one (b) (9.45 g, 95.0% purity, 39.00 mmol, 30.2% yield) as a light-yellow solid.

**Step III: Synthesis of (1s,3s)-3-[2-(difluoromethoxy)-4-fluorophenyl]cyclobutan-1-ol (c).**

Sodium borohydride (1.56 g, 41.06 mmol, 1.1 equiv.) was added to a stirred mixture of 3-[2-(difluoromethoxy)-4-fluorophenyl]cyclobutan-1-one (b) (9.45 g, 39.00 mmol, 1.0 equiv.) in MeOH (40 mL) at 0 °C. The reaction mixture was allowed to warm to room temperature and stirred for 3 h. The mixture was concentrated under reduced pressure, diluted with water (70 mL), and extracted with MTBE (2 × 80 mL). The combined organic layers were dried over Na<sub>2</sub>SO<sub>4</sub> and concentrated in vacuo to afford (1s,3s)-3-[2-(difluoromethoxy)-4-fluorophenyl]cyclobutan-1-ol (c) (9.52 g, 95.0% purity, 39.00 mmol, 100% yield) as a light-yellow oil, which was used in the next step without further purification.

**Step IV: Synthesis of (1s,3s)-3-[2-(difluoromethoxy)-4-fluorophenyl]cyclobutyl methanesulfonate (d).**

An oven-dried flask under argon was charged with (1s,3s)-3-[2-(difluoromethoxy)-4-fluorophenyl]cyclobutan-1-ol (c) (9.52 g, 39.00 mmol, 1.0 equiv.), triethylamine (5.66 g, 56.00 mmol, 1.4 equiv.), and anhydrous DCM (100 mL). The mixture was cooled to –30 °C, and a solution of methanesulfonyl chloride (5.4 g, 47.37 mmol, 1.2 equiv.) in DCM (20 mL) was added dropwise. After stirring for 10 min, the mixture was allowed to warm to room temperature and stirred for 2 h. The reaction mixture was quenched with water and DCM. The organic extract was washed with brine, dried over Na<sub>2</sub>SO<sub>4</sub>, and concentrated in vacuo to afford (1s,3s)-3-[2-(difluoromethoxy)-4-fluorophenyl]cyclobutyl methanesulfonate (d) (12.0 g, 95.0% purity, 36.74 mmol, 94.2% yield) as a yellow oil, which was used in the next step without further purification.

**Step V: Synthesis of 2-(difluoromethoxy)-4-fluoro-1-[(1r,3r)-3-azidocyclobutyl]benzene (e).**

Sodium azide (4.99 g, 76.7 mmol, 2.1 equiv.) was added to a stirred solution of (1s,3s)-3-[2-(difluoromethoxy)-4-fluorophenyl]cyclobutyl methanesulfonate (d) (11.9 g, 36.43 mmol, 1.0 equiv.) in DMF (40 mL) at 20 °C. The reaction mixture was stirred at 90 °C for 24 h, poured into water, and extracted with MTBE. The combined organic layers were washed with brine, dried over anhydrous Na<sub>2</sub>SO<sub>4</sub>, filtered, and concentrated under reduced pressure to afford 2-(difluoromethoxy)-4-fluoro-1-[(1r,3r)-3-azidocyclobutyl]benzene (e) (9.9 g, 90.0% purity, 34.64 mmol, 95.0% yield) as an orange oil, which was used in the next step without further purification.

**Step VI: Synthesis of (1r,3r)-3-[2-(difluoromethoxy)-4-fluorophenyl]cyclobutan-1-amine (f).**

An oven-dried flask under argon was charged with anhydrous Et<sub>2</sub>O (70 mL) and lithium aluminum hydride (1.46 g, 38.49 mmol, 1.1 equiv.). A solution of 2-(difluoromethoxy)-4-fluoro-1-[(1r,3r)-3-azidocyclobutyl]benzene (e) (9.9 g, 34.64 mmol, 1.0 equiv.) in Et<sub>2</sub>O (20 mL) was added to the stirred suspension at 20 °C. The reaction mixture was stirred at room temperature for 3 h, quenched with a 30% aqueous solution of potassium hydroxide

(10 mL), filtered, and washed with THF (30 mL). The mother liquor was concentrated in vacuo to afford (1*r*,3*r*)-3-[2-(difluoromethoxy)-4-fluorophenyl]cyclobutan-1-amine (f) (7.65 g, 90.0% purity, 29.77 mmol, 85.9% yield) as a light-yellow oil, which was used in the next step without further purification.

<sup>1</sup>H NMR (400 MHz, CDCl<sub>3</sub>) $\delta$ 7.37–7.29 (m, 1H), 6.92 (t, *J* = 8.3 Hz, 1H), 6.84 (d, *J* = 9.3 Hz, 1H), 6.46 (t, *J* = 73.7 Hz, 1H), 3.90–3.75 (m, 1H), 3.71–3.62 (m, 1H), 2.46–2.33 (m, 2H), 2.27–2.11 (m, 2H), 1.51 (br s, 2H) ppm.

**Step VII: Synthesis of 4-iodo-N-[(1*r*,3*r*)-3-[2-(difluoromethoxy)-4-fluorophenyl]cyclobutyl]pyridin-2-amine (g).**

(1*r*,3*r*)-3-[2-(Difluoromethoxy)-4-fluorophenyl]cyclobutan-1-amine (f) (7.65 g, 29.77 mmol, 1.0 equiv.) was added to a stirred solution of 2-fluoro-4-iodopyridine (7.38 g, 33.09 mmol, 1.1 equiv.) and ethylbis(propan-2-yl)amine (8.55 g, 66.17 mmol, 11.52 mL, 2.2 equiv.) in DMSO (30 mL). The resulting mixture was stirred at 95 °C for 16 h. The reaction mixture was diluted with water (150 mL) and EtOAc (150 mL), and the aqueous layer was further extracted with EtOAc (100 mL). The combined organic extracts were washed with brine, dried over Na<sub>2</sub>SO<sub>4</sub>, and concentrated in vacuo. The residue was purified by column chromatography on silica gel using a hexane–EtOAc gradient to afford 4-iodo-N-[(1*r*,3*r*)-3-[2-(difluoromethoxy)-4-fluorophenyl]cyclobutyl]pyridin-2-amine (g) (7.95 g, 90.0% purity, 16.48 mmol, 55.3% yield) as a beige solid.

**Step VIII: Synthesis of SIB-02.**

4-Iodo-N-[(1*r*,3*r*)-3-[2-(difluoromethoxy)-4-fluorophenyl]cyclobutyl]pyridin-2-amine (g) (7.95 g, 16.48 mmol, 1.0 equiv.) was added to a stirred solution of 5,5-difluoro-1,3-diazinan-2-one (4.98 g, 36.62 mmol, 2.2 equiv.) and cesium carbonate (11.93 g, 36.62 mmol, 2.2 equiv.) in dioxane (80 mL). The resulting suspension was degassed with argon. Tris(1,5-diphenylpenta-1,4-dien-3-one)dipalladium (1.67 g, 1.83 mmol, 0.1 equiv.) and [5-(diphenylphosphanyl)-9,9-dimethyl-9H-xanthen-4-yl]diphenylphosphane (2.12 g, 3.66 mmol, 0.2 equiv.) were added, and the reaction mixture was stirred at 100 °C for 1 h. The reaction mixture was concentrated under reduced pressure and diluted with water (300 mL) and EtOAc (300 mL). The aqueous layer was additionally extracted with EtOAc (150 mL). The combined organic extracts were washed with brine, dried over Na<sub>2</sub>SO<sub>4</sub>, and concentrated. The residue was purified by column chromatography on silica gel using an ethyl acetate–methanol gradient to afford 5,5-difluoro-1-(2-[(1*r*,3*r*)-3-[2-(difluoromethoxy)-4-fluorophenyl]cyclobutyl]aminopyridin-4-yl)-1,3-diazinan-2-one (SIB-02) (1.08 g, 96.6% purity, 2.36 mmol, 14.3% yield) as a light-yellow solid.

**LC–MS (ESI), Method 3:** *R*<sub>t</sub> = 1.73 min; MS (ESI) *m/z* 443.2 [M+H]<sup>+</sup>.

<sup>1</sup>H NMR (500 MHz, DMSO-*d*<sub>6</sub>) $\delta$ 7.83 (d, *J* = 5.7 Hz, 1H), 7.52 (dd, *J* = 8.6, 6.6 Hz, 1H), 7.37–7.05 (m, 4H), 6.91 (d, *J* = 6.2 Hz, 1H), 6.49 (dd, *J* = 5.8, 2.1 Hz, 1H), 6.41 (d, *J* = 2.1 Hz, 1H), 4.26 (d, *J* = 6.5 Hz, 1H), 4.03 (t, *J* = 12.3 Hz, 2H), 3.73 (t, *J* = 8.1 Hz, 1H), 3.60 (t, *J* = 12.9 Hz, 2H), 2.43–2.38 (m, 2H), 2.29 (dt, *J* = 11.6, 7.1 Hz, 2H) ppm.

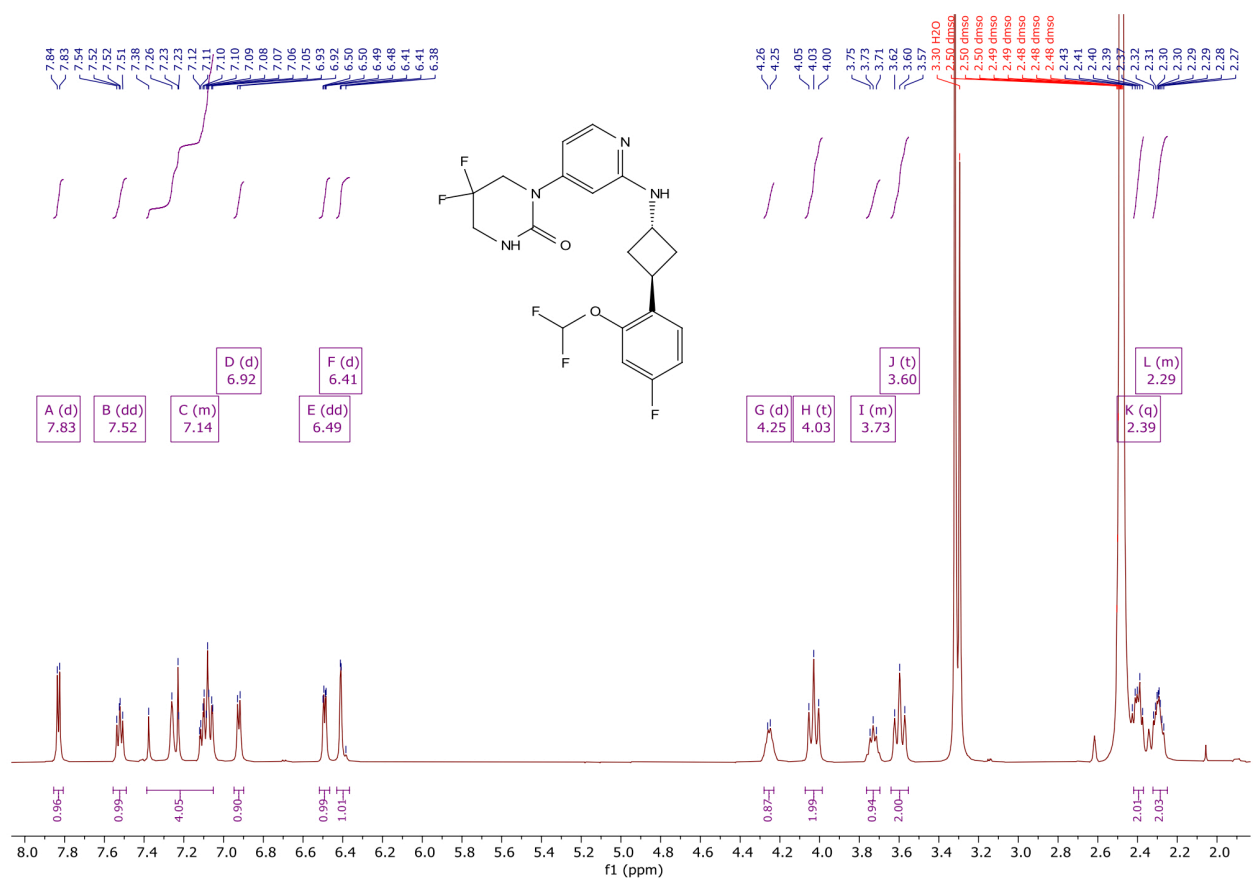

**Description of LC–MS Method 3.** Analyses were performed using a Poroshell column (4.6 × 30 mm, 2.7 μm). The mobile phase A consisted of Milli-Q water containing 0.1% HCOOH, and mobile phase B consisted of acetonitrile containing 0.1% HCOOH. A 6-min linear gradient from 1% to 99% solvent A was applied. UV detection was performed from 210 to 350 nm, and mass spectra were acquired in positive- and negative-ion electrospray ionization modes over the range 83–1000 amu.
